# From lab to field: analyses of genome-edited bacterial blight resistant rice

**DOI:** 10.1101/2025.05.26.656110

**Authors:** Eliza P.I. Loo, José Huguet-Tapia, Michael Selvaraj, Melissa Stiebner, Britta Killing, Marcel Buchholzer, Van Schepler-Luu, Thomas Hartwig, Sandra Valdéz Gutierrez, Madlen I. Rast-Somssich, Boris Szurek, Joe Tohme, Paul Charraviaga, Frank F White, Bing Yang, Wolf B. Frommer

**Affiliations:** Heinrich Heine University Düsseldorf, Faculty of Mathematics and Natural Sciences, Institute for Molecular Physiology, Düsseldorf, Germany; Cluster of Excellence on Plant Sciences (CEPLAS), Heinrich Heine University, 40225 Düsseldorf, Germany; Department of Plant Pathology, University of Florida Gainesville, FL 32611, USA; The Alliance of Bioversity International and International Center for Tropical Agriculture (CIAT), 763537 Cali, Colombia; International Rice Research Institute, DAPO Box 7777, Metro Manila, Philippines; Independent Research Group, Max Planck Institute for Plant Breeding Research, Cologne, Germany; Division of Plant Sciences, Bond Life Sciences Center, University of Missouri, Columbia, MO 65211, USA; Plant Health Institute of Montpellier (PHIM), Université Montpellier, IRD, CIRAD, INRAE, Institut Agro, Montpellier, France; Department of Plant Pathology, University of Florida, Gainesville, FL, USA; Donald Danforth Plant Science Center, St. Louis, MO 63132, USA; Institute for Transformative Biomolecules, ITbM, Nagoya, Japan

**Keywords:** *Xanthomonas oryzae pv. oryzae*, TAL effector, bacterial blight, genome editing, field trial, transgene analysis, whole genome sequencing, regulations, rice, Africa, India, yield

## Abstract

Bacterial blight (BB) of rice, caused by *Xanthomonas oryzae* pv. *oryzae* (Xoo) is one of the major drivers of yield losses in Africa and Asia. Xoo secretes TAL-effectors (TALe) that induce host SWEET sucrose uniporter by binding to the effector binding element (EBE) of *SWEET* promoters, likely required for Xoo reproduction and virulence. We had multiplex edited the EBEs of three SWEET genes to prevent TALe binding, producing genome-edited (GE’d) rice mega-varieties (IR64, Ciherang-Sub1 for Asia, and Komboka for Africa) that were resistant to a wide spectrum of Xoo strains. Here, we report comprehensive analyses of the GE’d lines, including evaluation of agronomic performance in multi-location multi-season experimental field plots under different fertilization regimes, and tests for the presence/absence of foreign DNA/transgene in the offspring of GE’d lines (IR64-BC1T6, Ciherang-Sub1-BC1T5, Komboka-T3). Various strategies were evaluated, including herbicide tolerance, PCR, DNA gel blotting, whole genome sequencing (WGS), and specific tests stipulated by country-specific biosafety guidelines. Different WGS technologies were evaluated and also used to identify heritability of the edits, single nucleotide polymorphisms (SNPs), and insertions/deletions (indels) that might have resulted from somaclonal variation and potential GE-induced off-target mutations. Complete genome reference sequences for the parental lines IR64, Ciherang-Sub1, and Komboka are provided. In the field experiments, the GE’d lines did not show performance defects. Together, the results indicate that select GE lines do not contain foreign DNA or transgene fragments and fulfill the requirements for treatment equivalent to classical breeding lines in countries such as India and Kenya.

## INTRODUCTION

In 2023, 800 million tons of rice (*Oryza sativa L.*) were produced, providing >20% of the total calories consumed worldwide (www.fao.org/faostat/en/#data/QCL). Still, in the same year, approximately 760 million of the >8 billion people on the planet faced hunger (http://www.who.int/publications/m/item/the-state-of-food-security-and-nutrition-in-the-world-2024). With the global population projected to reach around 10 billion by 2050, global food production will need to increase by nearly 40% to ensure adequate nutrition for everyone. (www.un.org/development/desa/pd/content/World-Population-Prospects-2022). Limited resources, national conflicts, and climate change are likely to exacerbate world hunger; therefore, ensuring food security is an urgent need. Pathogens and pests reduce yield and quality of agricultural production, causing substantial losses in addition to threatening food security and impacting a large number of small-scale food producers in Asia and Africa. Globally, yield losses for rice are estimated at 25 - 41%; highest losses are associated with food-scarce regions with endemic and emerging pests and diseases (Savary *et al*., 2019). Losses are severe in Southeast Asia and increasingly damaging in West African countries (Niño-Liu *et al*., 2006); and despite implementation of successful breeding programs, BB causes yield losses in the range of 8% in the Indo-Gangetian plain (Savary *et al*., 2019). Recent outbreaks in Tanzania and Madagascar caused by the inadvertent introduction of strains of Asian Xoo (Xoo^Asia^) strains are impacting rice production in East Africa (Raveloson *et al*., 2023; Schepler-Luu *et al*., 2023). Virulence of Xoo critically depends on Transcription Activation Like effector (TALe) proteins that are injected into host cells where they bind to effector binding elements (EBEs) and, in turn, induce transcription of at least one of three SWEET sugar uniporter genes (*SWEET11a, SWEET13* or *SWEET14*)(Chen *et al*., 2012, 2010; Oliva *et al*., 2019; Streubel *et al*., 2013; L. B. Wu *et al*., 2022). Notably, BB resistance had been found in a diverse germplasm due to nucleotide polymorphisms in EBEs in *SWEET* promoters: *xa13* (*SWEET11a*), *xa25* (*SWEET13*) (Liu *et al*., 2011; Zhou *et al*., 2015), and *xa47t* (*SWEET14*)(Hutin *et al*., 2015; Zaka *et al*., 2018). The resistance can be broken by strains that target either an alternate EBE in the same SWEET, or another sucrose transporting SWEET (Eom *et al*., 2019).

Theoretically, marker-assisted breeding could be used to combine *xa13*, *xa25,* and *xa41t*; this would, however, require challenging crosses between diverse germplasm, and importantly, since the SWEET14 promoter contains multiple EBEs, resistance would not cover all known strains/TALe. A more promising approach would entail multiplex editing of all known EBEs in the three *SWEET* promoters with CRISPR/Cas9 to confer broad-spectrum resistance to virtually any known rice cultivar in Asia and Africa (Bezrutczyk *et al*., 2018; Eom *et al*., 2019; Ni *et al*., 2021; Oliva *et al*., 2019; Ramasamy *et al*., 2025; Rifhani *et al*., 2023; Schepler-Luu *et al*., 2023). Multiplex editing had been used for the foundational breeding lines (mega varieties), including IR64, a rice variety widely grown in South East Asia, Ciherang-Sub1 (Bina Dhan11), a submergence-tolerant variety used widely in Indonesia, India and Bangladesh, and Komboka, a popular Kenyan variety (Ng’endo *et al*., 2022; Pramudyawardani *et al*., 2020; Toledo *et al*., 2015). The resulting genome-edited (GE’d) lines in the three elite backgrounds showed resistance to representative extant isolates and did not exhibit obvious defects in growth or habit in greenhouse or screenhouse trials. As of May 2025, twenty-one countries have guidelines exempting transgene-free GE’d crops from genetically modified organism (GMO) regulations (Buchholzer and Frommer, 2023)(www.editagenome.org). Here, we present comprehensive analyses of BB-resistant GE’d elite lines (IR64, Ciherang-Sub1, and Komboka) to lay the basis for transfer to small-scale food producers in Asia and Africa (Eom *et al*., 2019; Oliva *et al*., 2019; Schepler-Luu *et al*., 2023). A key prerequisite for classifying a GE’d crop as equivalent to classical breeding lines is the effective removal of foreign DNA/transgene fragments by segregation. However, a proof of removal is challenging since proving absence is not possible. After careful testing for the effective removal of foreign DNA/transgenes used for editing by segregation, GE’d lines are eligible for transfer to interested government organization in interested countries. Moreover, we show here in several experimental field plots performed in the absence of disease pressure that the GE’d IR64 and Ciherang-Sub1 lines do not show performance defects.

## RESULTS

As a basis for further dissemination, GE’d elite lines in three genetic backgrounds were evaluated for successful outcrossing of the transgenes and for field performance.

### Monitoring the presence of vector sequences in GE’d IR64 and Ciherang-Sub1 lines by PCR

Progenies from eleven GE’d IR64 lines (BC1T4) and eight GE’d Ciherang-Sub1 lines (BC1T3) resistant to Asian and African Xoo strains, previously validated to not carry the Cas9 gene, were transferred from IRRI to CIAT (Eom *et al*., 2019; Oliva *et al*., 2019). The GE’d IR64 and Ciherang-Sub1 lines underwent a multilayered pipeline for transgene analysis of GE’d plants (Figure 1). PCR was performed to screen for the presence of T-DNA in the GE’d lines using primer pairs specific to the CaMV35S promoter, gRNAs, ZmUBI10 promoter, and parts of *Cas9* (Table S1). PCR amplicons were not detected in any of the GE’d IR64 and Ciherang-Sub1 lines. Eleven GE’d IR64 and 8 GE’d Ciherang-Sub1 lines were advanced two generations before they were subjected to DNA gel blotting and short-read WGS (Figure S1a, described in detail below). Concurrently, a parental T2 GE’d IR64-C3 line was transferred from IRRI to the University of Missouri and advanced independently of the lines at CIAT. Since transformation could lead to insertion of vector backbone fragments, the PCR screens for T-DNA were not considered fully conclusive.

**Figure 1:**
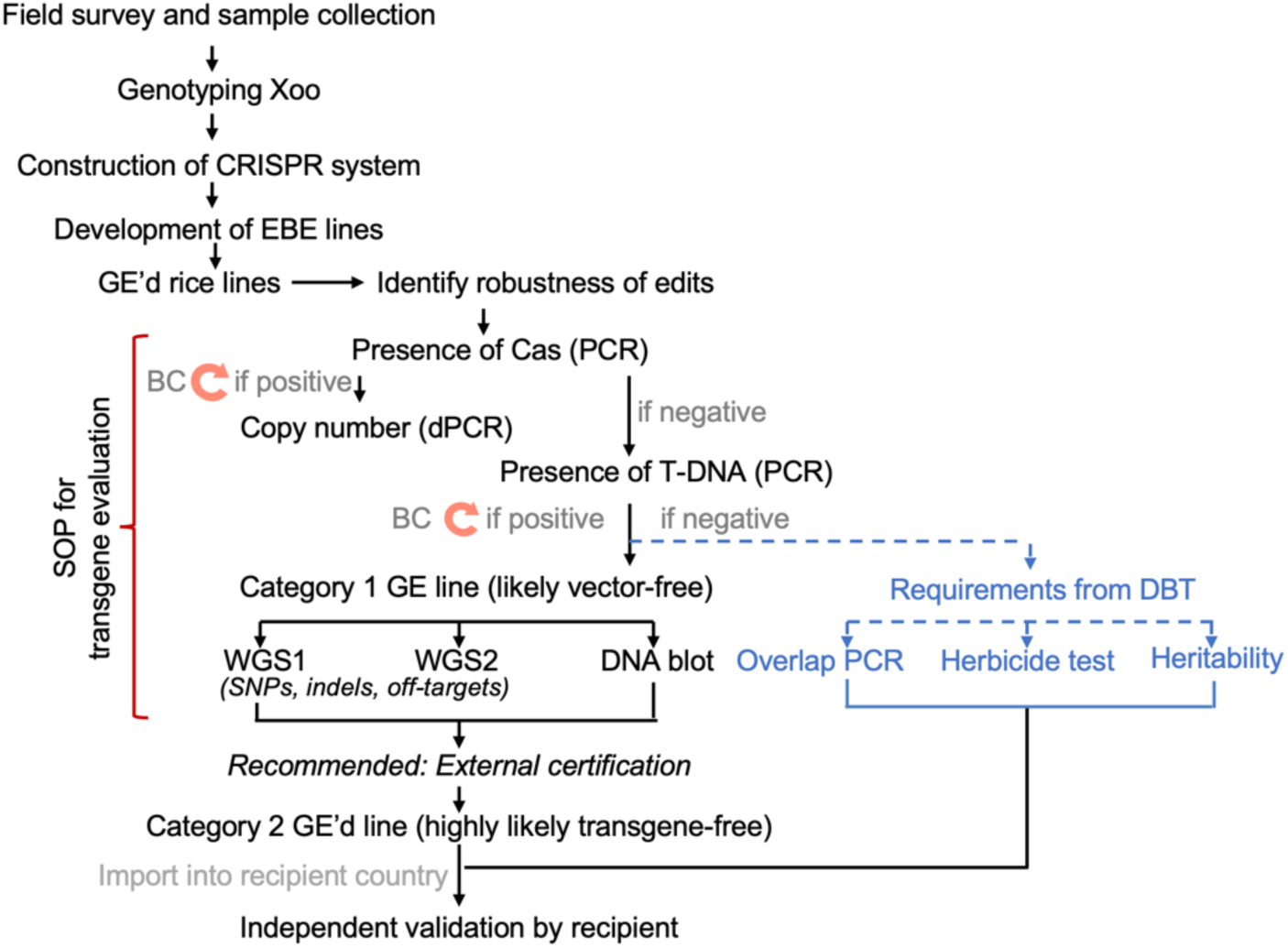
Transgene evaluation pipeline for validation of elimination of transgenes. After identifying lines with an optimal combination of edits, F2 lines are analyzed for the presence of transgenes. In the first step, lines are tested for the presence of Cas9 (or sequences close to the right border of the T-DNA) using RT-PCR and validated primers (including negative and positive controls). In parallel, it is useful to determine the copy number, e.g., by digital PCR (Schepler-Luu *et al*., 2023). Lines that do not appear to contain Cas9 are subjected to DNA gel blots using fragments covering the vector, overlap PCR (max amplicon size 500 bp, minimal overlap 50 bp) as well as tests for the loss of the selectable marker (here hygromycin resistance) used for transformation. Lines are subjected to WGS in two rounds using independent enzymatic and mechanical fragmentation (provided Illumina or technologies that require fragmentation are used). Alternatively, and preferably, WGS is performed using long read WGS technology with adequate coverage. It may be useful to obtain external validation and certification that the lines are transgene free based on certified GM tests. If in one of the steps indications for the presence of a transgene are identified, lines will be backcrossed (BC) and subsequently have to reenter the analysis pipeline.

### DNA hybridization analyses for detecting vector sequences

To detect potential vector fragment integration, DNA gel blots were performed using eight overlapping DNA fragments as probes for detecting IRS1132 in six GE’d IR64 and two GE’d Ciherang-Sub1 lines (Figure S2, Table S1). Wild-type (WT) plants of the respective cultivars served as negative controls. Parental lines carrying vector integration (IR64-136b for IR64 lines and CS-3d for Ciherang-Sub1 lines) and WT plant DNA spiked with vector DNA of various copy number reconstructions were used as positive controls. In controls, DNA gel blots using all eight probes detected the presence of vector spiked into the genomic DNA of WT plants at the sensitivity of ≤0.5 copies of vector per genome (Figure S2). The probes did not detect vector sequences in the eight GE’d lines, providing support for the hypothesis of the absence of vectors at the levels of 0.5 copies per genome (Figures S2). Notably, however, we assume that the DNA gel blots cannot detect small insertions (<30bp), and the sensitivity may depend on the length of the hybridizing sequence.

### Whole Genome Sequencing of GE’d IR64 and Ciherang-Sub1 lines

As a potentially more reliable alternative to detecting vector integration, WGS may be suitable for identifying transgene fragments. Genomic DNA of 11 GE’d IR64 lines and eight GE’d Ciherang-Sub1 lines were analyzed by short-read paired-end WGS (Figure S1a). The first protocol used enzymatic digestion of the genomic DNA in library preparation for sequencing. Since enzymatic digestion of genomic DNA exhibits restriction site bias and may introduce indels, resulting in incomplete coverage (Knierim *et al*., 2011; Sato *et al*., 2019), a second round of WGS was performed using mechanical shearing via sonication. Leaf samples were pooled from two individual plants as a biological replicate. Genome sequences of GE’d IR64 and Ciherang-Sub1 line were mapped to the vector pBY02UbiOsCas9_gSWEET11_13_14 (hereafter, IRS1132). We applied stringent inspection criteria to classify reads mapped to the vector as ‘true’ genome integration if:

i) Mapped reads were not singletons/orphan reads, i.e., mapped reads must be paired. Since pair-end WGS was employed, each sequenced fragment should produce two reads. Singletons were defined here as sequencing fragments with only one of the reads successfully aligned to the vector.
ii) For chimeric reads, soft-clipped reads had to map to the rice genome. For discordant read pairs, one pair had to map to the genome, and the other pair to the vector.
iii) Mapped reads did not correspond to the native rice-derived U3 promoter (on the vector driving gRNA) or EBE (on the vector as gRNA).
iv) Mapped reads were consistently found in all biological replicates in both WGS protocols.

Based on the criteria listed above, vector integration was not detected in six GE’d IR64 and two Ciherang-Sub1 lines, namely, IR64-5d, IR64-9a, IR64-134b, IR64-136a, IR64-7b, IR64-C3, CH-3b, and CH-4a (see Supplemental Methods for details, Figures 2, S3, Table S2). An insertion of 273 bp derived from *Cas9*, i.e., a partial insertion without adjacent T-DNA sequences, was found on chromosome 12 of IR64-7a. To validate that singletons were not caused by small insertions, the genomes of a representative IR64 line, i.e., IR64-C3 (BC2T5), that had been subjected to short-read and long-read (without barcoding/multiplexing) WGS were compared. Notably, none of the singletons in IR64-C3 sequenced via short-read sequencing that mapped to the vector IRS1132 were found in the genome derived from long-read sequencing (Figures 2, S3, Table S2). Since samples were pooled prior to short-read sequencing, we posit that reads mapped to the vector may be caused by index hopping, i.e., an artifact generated by misassignment of reads during demultiplexing (Kircher *et al*., 2012).

**Figure 2:**
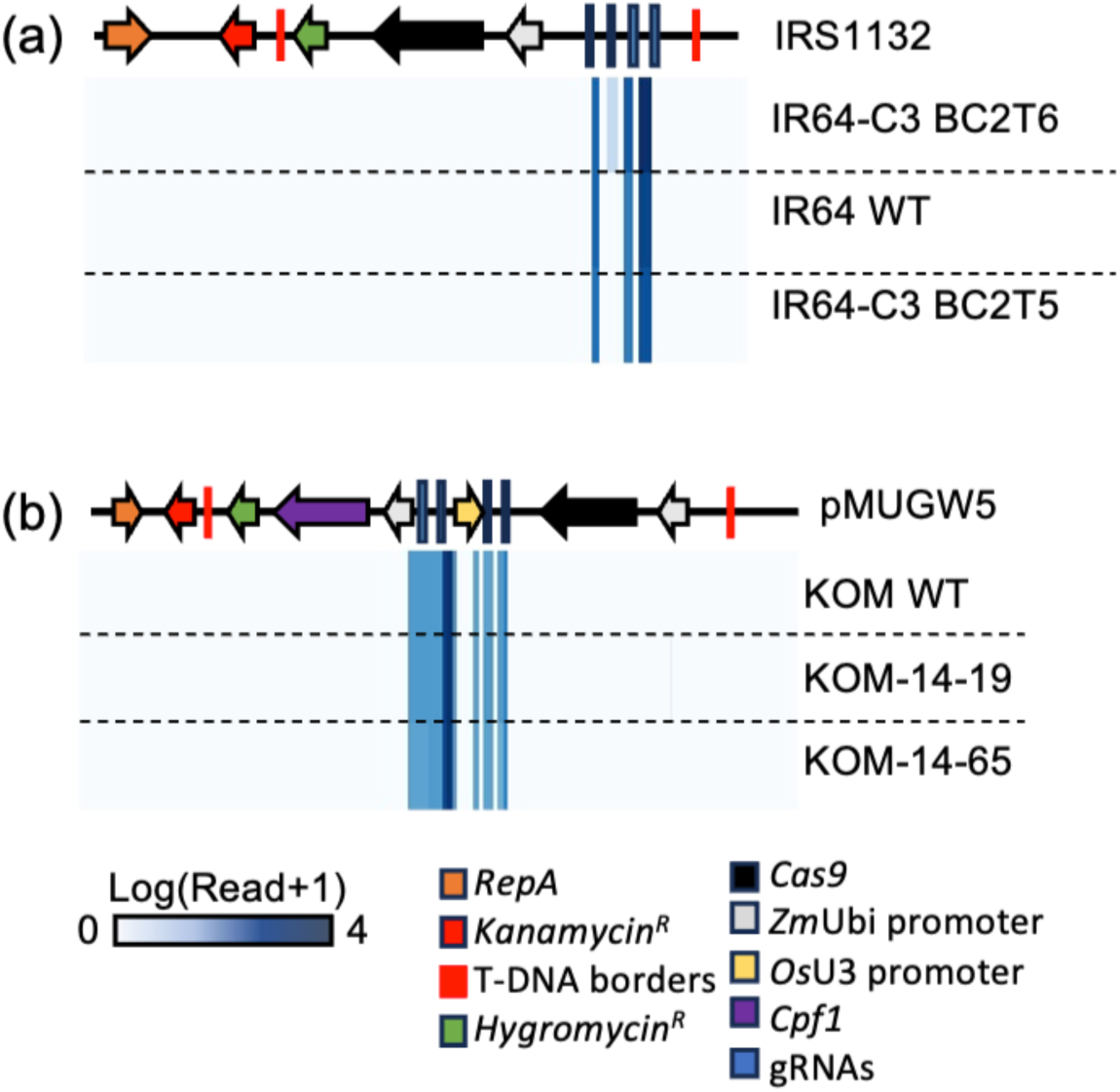
Analyses for vector integration in GE’d rice lines. A) Analysis for the presence of vector sequences by WGS. (a) Alignment of GE’d IR64 and Cihernag-Sub1, and (b) Komboka) against vector sequences used for genome editing (IRS1132 for IR64 and Ciherang-Sub1, pMUGW5 for Komboka). Linearized IRS1132 and pMUGW5 indicated on the top of respective genome alignments. Legends indicate components in vectors IRS1132 and pMUGW5. Blue scale bar indicates number of reads in the sequenced genome of GE’d lines mapped to respective vectors.

### Identification of SNPs, indels, and off-target mutations in GE’d IR64 and Ciherang-Sub1 lines

*Agrobacterium*-mediated transformation of rice involves extended tissue culture phases known to induce somaclonal variation (Miyao *et al*., 2012). In addition, new ‘natural’ mutations occur in the progeny. To evaluate new mutations, the genomes of the GE’d lines were inspected for SNP and indels. To avoid assessing SNPs/indels generated due to sequencing errors, only indels/SNPs consistently detected in all replicates from both WGS protocols (i.e., consensus indels/SNPs) were considered (Liu *et al*., 2021). Indels/SNPs in a representative GE’d line, IR64-C3 were mapped across three generations, including IR64-C3 T1, the T3 generation backcrossed once with WT IR64 (IR64-C3 BC1T3), and a fifth-generation progeny from two backcrosses (IR64-C3 BC2T5). In the T1 generation, 303 indels/SNPs were evenly distributed across all chromosomes (Figures 3a, S4a). In IR64-C3 BC1T3, the number of indels/SNPs was reduced to 131 and further reduced to 120 in IR64-C3 BC2T5, indicating successful reduction of somaclonal variation. The natural mutation rate was estimated to be 5.4 x 10^-8^ per rice generation (Tang *et al*., 2018). The GE’d IR64 and Ciherang-Sub1 lines have been cultivated for seven and six generations, respectively, after *Agrobacterium*-mediated transformation. Therefore, the theoretical number of ‘natural’ SNP/indel expected after seven generations of cultivation was 163 and 139 after six generations. The observed number of SNPs/indels in IR64-9a BC1T6 was 242, 270 in IR64-136a BC1T6, and 226 in IR64-7b, whereas 48 SNPs/indels in CS-3a BC1T5 and 74 SNPs/indels in CS4 BC1T5 were within the expected range (Figures S4b-c).

**Figure 3:**
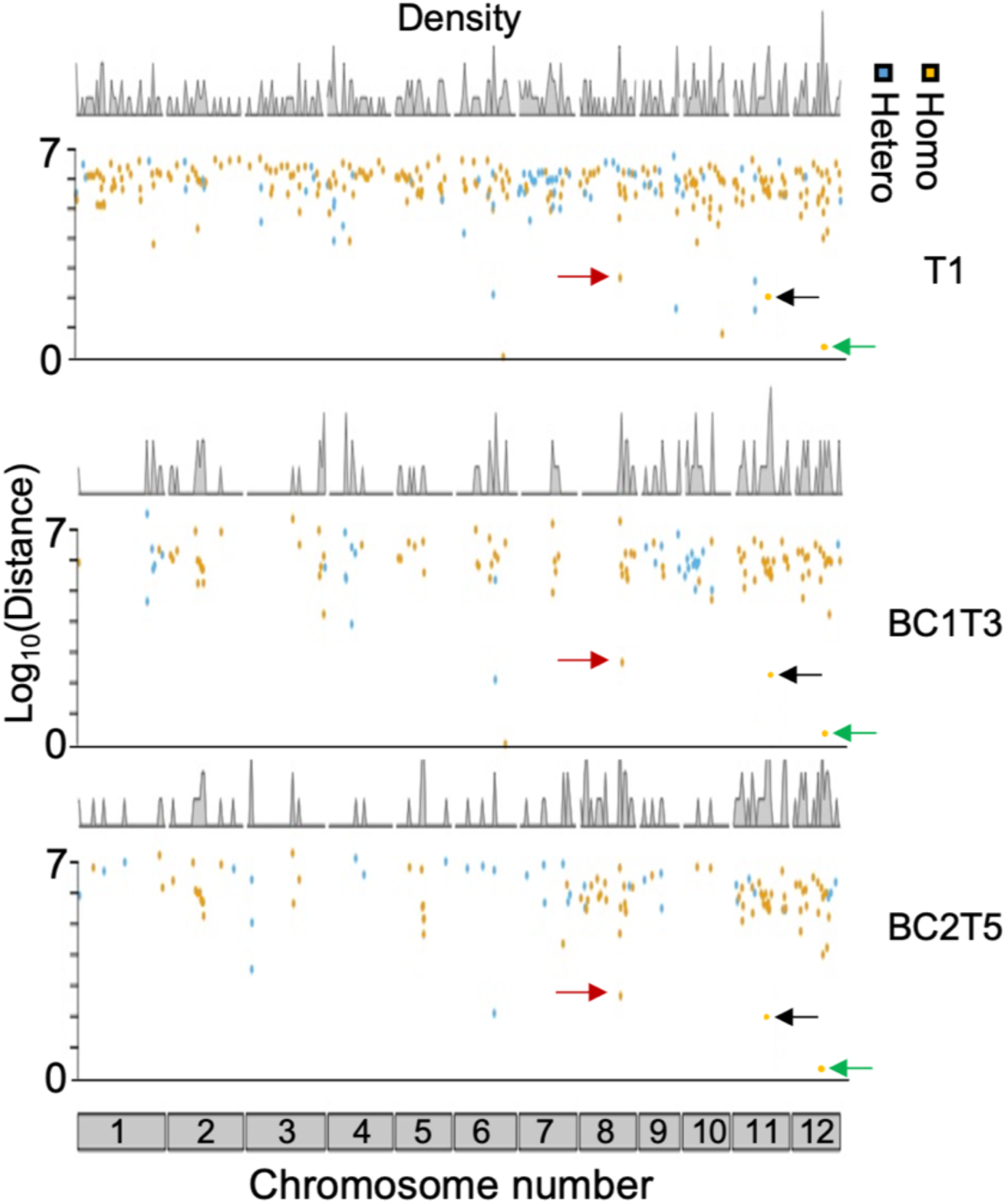
Analysis of SNPs, indels, and potential off-target mutations in GE’d rice lines. Density and distribution of SNPs detected in each chromosome of IR64 C3 (BC2T5) and its parental genomes (T1 and BC1T1). Colors of dots indicate type of SNP, i.e. Homo = homozygous SNP, Hetero = heterozygous SNP. Location of SNPs, i.e., chromosome number indicated on the X-axis. Red arrows indicate mutation introduced in EBE of PthXo1, black arrows indicate mutation introduced in EBEs of PthXo3, AvrXa7, and TalF, green arrows indicate mutation introduced in EBE of PthXo2.

A range between 10 and 56 indels, and 38 and 214 SNPs were identified in the genomes across all GE’d lines (Figure S4b). Indels and SNPs were traced to their respective genes/chromosomal locations. Between 31 and 176 SNPs were found in intergenic regions (>75%); 5-30 non-intergenic SNPs were present in introns (3-24%), and <20 SNPs fell into exons (<10.5%). Exonic SNPs were further inspected for potential disruption of gene function. Between two and five exonic SNPs were silent mutations; 1-16 were missense mutations. Between 9 and 42 indels were in intergenic regions, 0-13 indels in introns, and one indel in the exon of LOC_Os02g32570.1 (NF2 family N-terminal domain-containing protein), resulting in an in-frame deletion in IR64-136a (Table S3). Lines IR64-7b, CS-3b, and CS-4a, carried one indel each that results in a frameshift in g9701 (in IR64-7b; new gene of unknown function found by reannotation), LOC_Os01g13760.1 (in CS-3b; heat shock protein 40)(Sarkar *et al*., 2013), and LOC_Os1g50320.2 (in CS-4; suppressor-of-white-apricot (SURP) module family protein)(Fan *et al*., 2024; Nameki *et al*., 2022) (Figure S3b, Table S3).

### Potential off-target mutations detected by WGS in GE’d IR64 and Ciherang-Sub1 lines

Cas9 tolerates up to three mismatches to the gRNA and can cause off-target mutations. We analyzed the genome sequences of the GE’d lines for potential off-target mutations. *Bona fide* off-target mutations were considered only for mismatches for the sequence corresponding to the guide RNA plus five bp up- or downstream (Hsu *et al*., 2013). All four gRNAs were randomly (*in silico*) mutated with 1-5 mismatches, and the genomes of GE’d lines were subsequently screened for potential mutated gRNA targets. Two potential Cas9 off-targets were detected in the genome of IR64-136a, and one potential off-target in the genomes of IR64-9a and IR64-7b. The potential off-target mutations, i.e., one-bp insertion in chromosome 1 position 6314428 in IR64-9a, IR64-136a, and IR64-7b; and eight-bp deletion in chromosome 5 position 15429580 in IR64-136a, are likely only in the event where the gRNA targeting the EBE of PthXo2 undergoes three random mutations (Table S4). All potential off-targets were traced to intergenic regions.

### Analysis of vector integration, SNPs/indels, and off-target mutations for GE’d Komboka lines

Progenies of GE’d Komboka lines previously validated to be resistant to Asian and African Xoo (Schepler-Luu 2023), i.e., eight T3 lines and four T2 lines, were subjected to our multilayer pipeline for transgene analysis (Figure 1). Of the eight T3 lines and five T2 lines screened via PCR using primers specific to *Cas9* fragment, amplicons were not detected in all eight T3 lines and three T2 lines (Figure S1b). KOM-14-19 and KOM14-65 were advanced to T3 and T4, respectively, for WGS validation for the absence of vector integration. To avoid the possibility of index hopping, GE’d KOM14-19 and KOM14-65, along with WT plants, were axenically grown and whole-genome sequenced using long-read WGS without multiplexing. Vector integration was not detected in GE’d Komboka lines based on the criteria used for evaluating GE’d IR64 and Ciherang-Sub1 lines (Figure 2b). SNPs and indels analyses indicated that KOM-14-19 carried one indel that results in a frameshift in LOC_Os03g57720.1, encoding a putative aldehyde oxidase 2)(J. Wu *et al*., 2022) (Figure S4B-C, Table S3). Cas9 off-target analysis did not reveal potential off-targets in GE’d Komboka lines (Table S4).

### Tests to fulfill specific requirements set by the Department of Biotechnology (DBT), India

To classify GE’d crops as conventional breeding materials, and as a prerequisite for import, the DBT guidelines stipulate two essential tests for the presence of exogenously introduced DNA in the genome of GE’d plants: (i) successful removal of the selection/screenable markers, and (ii) overlap PCR on genomic DNA for vector sequences used for generating the DNA modification. T-DNA-containing T3 parents, i.e., IR64-136b for IR64, CS-3d for Ciherang-Sub1, and T2 Komboka parent KOM-18-68 were used as positive parental controls to test for herbicide resistance. In contrast to controls containing the T-DNA, WT and GE’d lines without detectable vector integration, as determined by WGS analyses, did not grow on media supplemented with hygromycin (Figure S5a). Accordingly, overlap PCR using 49 (IRS1132 in Ciherang-Sub1), 50 (IRS1132 in IR64) and 59 (pMUGW5 in Komboka) primer pairs (Table S1) did not produce amplicons in WT and GE’d lines but expected amplicons (length ≤500 bp; ≥ 50 bp overlap) were observed in two positive controls, i.e., (i) a segregating parental line harboring full or part of the vector in its genome, and (ii) genomic DNA of the WT plant spiked with the vectors diluted 1000-fold (Figure S5c-e). In addition to foreign DNA/transgene analysis, DBT also requires evidence that the edited traits are heritable. The EBEs from two generations of each GE’d line were sequenced at least twice and aligned to ensure that the EBEs were inherited. The edited EBEs were identical across all generations of the GE’d line tested (Table S5), indicating the heritability of the edited trait. The information generated for transfer to India was sufficient to allow for the import of the edited lines into Kenya (KALRO).

### Genome-edited rice lines did not exhibit yield penalties in experimental field plots (EFP)

To evaluate whether the *Agrobacterium*-mediated editing process impacted growth or yield, as assessed under open-field conditions, EFPs were performed in the absence of BB disease pressure. Opportunities for EFP were limited due to regulations for genetically modified organisms. EFPs were performed at CIAT in Palmira, Valle de Cauca, Colombia, due to the availability of permitted and suitable infrastructure and the absence of BB. Eight GE’d IR64 and six Ciherang-Sub1 lines qualified for EFPs after quarantine measures were advanced to BC1T5 for agronomic performance testing (Figure S1a). Unmanned aerial vehicle (drone)-based multispectral analyses were performed in EFPs across two planting seasons. The first trial took place during the 2020 dry season (June 2020 to October 2020), and the second trial during the 2021 wet season (January 2021 to May 2021). Normalized Difference Vegetation Indices (NDVI) and Normalized Difference Red Edge Indices (NDRE) were used as quantitative measurement proxies for canopy development and chlorophyll levels (Figure 4a) (Huang *et al*., 2021). NVDI and NDRE values range between 1 and -1, with higher values indicating optimal vegetation scores. Both the WT and GE’d lines showed high NDVI value ranges of 0.8 - 0.91 in the dry season, indicating high photosynthetic capacity (Tables S6, S7). NDVI values for GE’d and WT plants in the wet season of 2021 were lower compared to NVDI values from the dry season of 2020, ranging from 0.33 - 0.83. The NDRE values for the GE’d and WT lines ranged from 0.29 and 0.55 during the dry season and between 0.2 and 0.5 during the wet season (Tables S6, S7). The lower NDVI and NDRE values obtained in the wet season may be a consequence of the overall reduced sunlight (Figure S6a). NDVI and NDRE indices showed no significant differences in the overall photosynthetic capacity and canopy development between GE’d and WT plants across both planting seasons (Figure 4a, Tables S6, S7). Additionally, eleven agronomic traits were quantified in three EFPs, including two trials in Colombia (dry season from June 2020 to October 2020, and wet season from January 2021 to May 2021), and one trial in the Philippines (wet season from December 2020 to March 2021) (Tables 1, S8). Across the three EFPs, agronomics trait evaluation indicated that the GE’d IR64 and Ciherang-Sub1 cultivars did not exhibit penalties in agronomic traits and total yield (Tables 1, S8). Reduced 1000-grain weight was observed from GE’d IR64 lines, an average of 27.3 g compared to 29 g from WT plants in the dry season, and an average of 26.4 g compared to 28.7 g from WT plants in the wet season. However, bulk grain weight, grain yield per m^2^, single plant grain yield, and yield per hectare of GE’d IR64 lines were not different relative to WT IR64 controls (Table S8). Principal component analysis clustered the rice lines based on geographical and seasonal harvest, and not by genotypic differences (Figure 4b). Drone-based multispectral and agronomic performance analyses were performed to further evaluate whether stress conditions might impact the performance of the GE’d lines with reduced nitrogen. Neither NDVI and NDRE at 1- to 2-week intervals nor agronomic traits of plants grown with or without N-fertilizer supplementation (0, 90 and 180 kg N ha^-1^; basal nitrogen content of 60 kg ha^-1^) had differences between GE’d and WT plants (Figure S6b, Tables S9, S10). PCA analysis did not separate between the different GE’d lines, and between GE’d and respective WT lines (Figure S6b).

**Figure 4:**
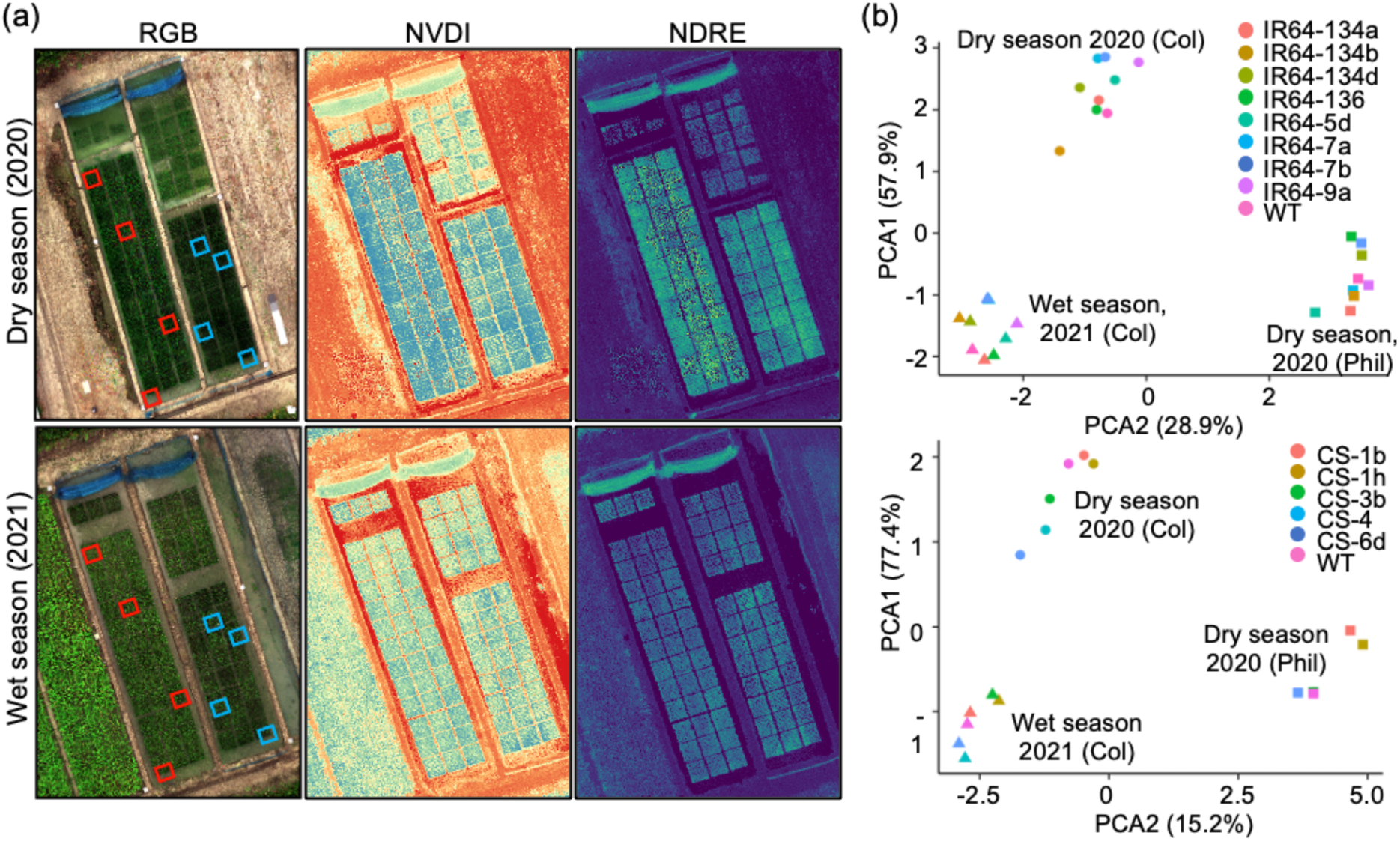
Agronomic performance of GE’d IR64 and Ciherang-Sub1 lines are not significantly different from parental WT plants. (a) Red-Green-Blue (RGB) and multispectral images of the EFPs conducted in Colombia in the dry season of 2020 and wet season of 2021. Boxed in red indicates WT IR64 plants, and boxed in blue indicates WT Ciherang-Sub1 plants. NVDI-Normalized Difference Vegetation Indices, NDRE-Normalized Difference Red Edge Indices. (b) Principal component analysis of WT and GE’d IR64 (top) and Ciherang-Sub1 (bottom) lines from three EFPs in two geographically separated fields and two harvest seasons. Colors indicate independent GE’d or WT lines, shapes indicate the location of EPF: Col-Colombia, Phil-The Philippines.

**Table 1:** Agronomic traits of GE’d and WT plants of IR64 and Ciherang-Sub1 cultivars during the 2021 wet season in Palmira, Valle del Cauca, Colombia.

| Genotype | 100% Flowering (DAP) | Bulk Grain weight (g) | Dry Biomass (g) | Grain Yield (kg/m<sup>2</sup>) | Harvest Index | Panicle Length (cm) | Panicle Number | Plant height (cm) | Single Plant grain Yield (g) | Yield (Ton/Ha) |
| --- | --- | --- | --- | --- | --- | --- | --- | --- | --- | --- |
| IR64-134c | 75.5 ± 0.5 | 1658.75 ± 77.7 | 30.78 ± 1 | 0.83 ± 0.04 | 0.52 ± 0.02 | 24.31 ± 0.1 | 12.18 ± 0.2 | 82.43 ± 0.9 | 35.49 ± 1.7 | 4.15 ± 0.3 |
| IR64-7b | 79.5 ± 8.5 | 1561.5 ± 125.2 | 32.78 ± 4.1 | 0.98 ± 0.1 | 0.55 ± 0.06 | 24.34 ± 0.4 | 12.68 ± 0.7 | 76.79 ± 1.1 | 39.07 ± 4.1 | 3.9 ± 0.2 |
| IR64-136a | 71.75 ± 0.9 | 1485 ± 64.9 | 35.24 ± 3.2 | 0.89 ± 0.13 | 0.5 ± 0.08 | 24.1 ± 0.3 | 14 ± 0.6 | 75.89 ± 1.1 | 33.62 ± 5.1 | 3.71 ± 0.4 |
| IR64-134b | 73.25 ± 0.75 | 1458.5 ± 151.3 | 28.55 ± 5.18 | 0.84 ± 0.07 | 0.54 ± 0.05 | 23.98 ± 0.3 | 12.75 ± 1.2 | 76.71 ± 1.1 | 28.32 ± 2.4 | 3.65 ± 0.2 |
| IR64-134a | 71.25 ± 0.75 | 1396.25 ± 71.6 | 32 ± 4.7 | 0.71 ± 0.04 | 0.48 ± 0.03 | 24.41 ± 0.4 | 11.14 ± 0.7 | 79.64 ± 0.9 | 29.44 ± 1.8 | 3.49 ± 0.3 |
| IR64-9a | 70.25 ± 0.25 | 1440.25 ± 110.8 | 30.79 ± 3.3 | 0.74 ± 0.08 | 0.49 ± 0.05 | 25.05 ± 0.6 | 12.86 ± 0.8 | 82 ± 0.1 | 32.35 ± 3.4 | 3.6 ± 0.4 |
| IR64-7a | 72 ± 0.8 | 1475 ± 172.8 | 28.77 ± 2.5 | 0.81 ± 0.05 | 0.53 ± 0.03 | 24.63 ± 0.3 | 11.43 ± 0.9 | 82.71 ± 1 | 29.56 ± 1.8 | 3.69 ± 0.1 |
| IR64-5d | 71 ± 1 | 1517.5 ± 48.1 | 32.53 ± 1.6 | 0.74 ± 0.04 | 0.48 ± 0.02 | 24.91 ± 0.2 | 11.86 ± 0.1 | 81.54 ± 0.9 | 29.46 ± 1.5 | 3.79 ± 0.2 |
| IR64 WT | 74.75 ± 1.4 | 1403 ± 70.3 | 30.35 ± 3 | 0.74 ± 0.01 | 0.49 ± 0.01 | 24.41 ± 0.3 | 12.61 ± 0.6 | 81.61 ± 1.3 | 33.21 ± 0.4 | 3.51 ± 0.2 |
| CS-6d | 78 ± 0.7 | 1486.75 ± 62.8 | 24.25 ± 2.1 | 0.92 ± 0.09 | 0.6 ± 0.06 | 22.82 ± 0.4 | 9.5 ± 0.6 | 77.82 ± 1 | 36.61 ± 3.5 | 3.72 ± 0.2 |
| CS-1b | 79 ± 1.1 | 1895.25 ± 55.8 | 24.56 ± 0.8 | 0.88 ± 0.02 | 0.59 ± 0.01 | 24.63 ± 0.2 | 9.39 ± 0.3 | 75 ± 2.7 | 35.31 ± 0.7 | 4.74 ± 0.1 |
| CS-4 | 77.75 ± 1.2 | 1500.25 ± 56.3 | 25 ± 1.5 | 0.9 ± 0.06 | 0.59 ± 0.04 | 23.29 ± 0.3 | 10.21 ± 0.4 | 74.04 ± 0.7 | 35.96 ± 2.4 | 3.75 ± 0.1 |
| CS-3b | 77 ± 0.6 | 1664.25 ± 74.3 | 27 ± 1.1 | 0.99 ± 0.05 | 0.59 ± 0.03 | 23.63 ± 0.3 | 10.36 ± 0.2 | 77.86 ± 1.2 | 39.56 ± 2 | 4.16 ± 0.2 |
| CS-1h | 80 ± 0.4 | 1631.25 ± 131.5 | 28.85 ± 2.9 | 0.94 ± 0.09 | 0.57 ± 0.05 | 24.47 ± 0.3 | 9.93 ± 0.6 | 81.79 ± 1.1 | 37.6 ± 3.6 | 4.08 ± 0.3 |
| CS WT | 76.75 ± 1.1 | 1760 ± 78.6 | 24.28 ± 1.2 | 0.9 ± 0.05 | 0.6 ± 0.03 | 23.16 ± 0.4 | 10.04 ± 0.8 | 76.1 ± 1.1 | 36.2 ± 1.9 | 4.4 ± 0.2 |

**Table 2:** Transgene evaluation of GE’d lines according to transgene analyses.

| GE line | PCR<br>Cas9 | PCR<br>vector | WGS | DNA<br>blot | Overlap<br>PCR | Hyg<sup>R</sup> | Transfer |
| --- | --- | --- | --- | --- | --- | --- | --- |
| IR64-5d | nd | nd | nd | nd | nd | R |  |
| IR64-7a | nd | nd | 273 bp *Cas9* | nt | nt | nt |  |
| IR64-9a | nd | nd | nd | nd | nd | R | Kenya |
| IR64-134a | nd | nd | Recurring<br>singletons | nt | nt | nt |  |
| IR64-134b | nd | nd | nd | nd | nd | R |  |
| IR64-136a | nd | nd | nd | nd | nd | R | Kenya |
| IR64-7b | nd | nd | nd | nd | nd | R |  |
| IR64-134d | nd | nd | Recurring<br>singletons | nt | nt | nt |  |
| IR64-C3 | nd | nd | nd | nt | nt | nt | Kenya |
| CS-3b | nd | nd | nd | nd | nd | R |  |
| CS-1h | nd | nd | Recurring<br>singletons | nt | nt | nt |  |
| CS-4 | nd | nd | nd | nd | nd | R |  |
| CS-1b | nd | nd | Recurring<br>singletons | nd | nt | nt |  |
| CS-6d | nd | nd | Recurring<br>singletons | nd | nt | nt |  |
| KOM-14-19 | nd | nd | nd | nt | nd | R | Kenya |
| KOM-14-65 | nd | nd | nd | nt | nd | R | Kenya |

Singletons: Sequenced fragments with only one read aligned to vector
Recurring singletons: Singletons found in more than one biological replicate
nd: not determined
nt: not tested

## DISCUSSION

We generated broad-spectrum resistance using multiplex editing of SWEET promoter elements in four rice varieties: Kitaake, IR64, Ciherang-Sub1, and Komboka. Here, we report the next critical steps, i.e., a comprehensive laboratory and field analysis of GE’d lines before transferring the GE’d lines to interested partners from countries that have established suitable regulations to exclude GE’d products from their national GMO regulations (www.editagenome)(Buchholzer and Frommer, 2023). We tested for the presence of foreign DNA and transgenes (used in the T0 generation to introduce edits) after segregation and subsequently evaluated the performance of GE’d lines in experimental field plots, under government-approved permits. We provide a set of GE’d lines in three elite varieties that are compatible with the criteria set by government agencies in India and Kenya, required for import, further characterization, and possibly variety registration under regulations that would consider these lines as equivalent to classical breeding lines.

### Segregation of foreign DNA/transgenes to obtain GE’d elite lines that can be exempted from country-specific GM regulations

Stable *Agrobacterium*-mediated transformation was used to introduce editing enzymes and gRNAs, thus the original transformants (before segregation) are classified as GMOs. T-DNA insertions can be eliminated by segregation. Since *Agrobacterium*-mediated transformation can lead to multiple T-DNA integrations, partial T-DNA insertions, as well as vector fragment insertions, it can be challenging to eliminate the T-DNA and to test for effective outcrossing (Norris *et al*., 2020). For instance, up to six Cas9 fragments were detected in the GE’d Komboka lines, requiring large populations and multiple generations to identify lines in which Cas9 was not detected anymore (Figure S7a). Until March 2025, 21 countries had implemented regulations that exempt GE’d plants that do not contain foreign DNA from respective national GM regulations, e.g. India and Kenya (www.editagenome.org). In 2023, the Department of Biotechnology (DBT), India, published guidelines with detailed requirements for the evaluation of the absence of ‘exogenous’ DNA from GE’d plants (https://dbtindia.gov.in/latest-announcement/sops-regulatory-review-genome-edited-plants-under-sdn-1-and-sdn-2-categories). For deregulation and import of GE’d seeds into India, the DBT guidelines requires validation on the successful removal of the selection markers, and overlap PCR on genomic DNA for vector sequences used for generating the DNA modification. In Kenya, GE’d crops are not subject to GMO regulation if the genetic modification was introduced using SDN1 or SDN2 approaches and if no foreign DNA or recombinant sequences are present in the final product. According to the "<u>Guidelines for Determining</u> <u>the Regulatory Process of Genome Edited Organisms and Products in Kenya</u>", applicants must submit a notification dossier to the National Biosafety Authority (NBA), including a scientific description of the editing process and sufficient evidence demonstrating the absence of foreign DNA. The data generated for import into India for overlap PCR and loss of herbicide resistance were accepted by the NBA in Kenya as sufficient for import.

DBT regulations stipulate that assays need to be performed side-by-side with a segregant line that retained the T-DNA as positive controls and untransformed parental lines as negative controls. Our analysis demonstrates that while the T-DNA containing parental line is herbicide tolerant, the negative control and the segregant edited lines are sensitive. We note, however, that we had to analyze multiple parental lines since some of the T-DNA containing lines had lost herbicide tolerance likely due to gene silencing (Figure S5b). The absence of herbicide tolerance, therefore, cannot serve as reliable evidence for the absence of the T-DNA. The results also do not exclude potential vector backbone integration into the genome (De Buck *et al*., 2000; Lange *et al*., 2006). Overlap PCR with respective positive and negative controls did not detect vector fragments (Figures S2, S7b), although together with the herbicide tolerance tests, the data comply with the regulations. However, trying to learn from the identification of a 3.9 kb fragment derived from vector backbone in the genomes of GE’d hornless cows by WGS previously undetected via diagnostic PCR (Carlson *et al*., 2016; Young *et al*., 2020), we intimate that fragments, either from partial T-DNA insertions or vector backbone can only be detected if both primers anneal to the inserted fragment (Figure S6b). We also note that the design and testing of the primer set is challenging and costly, since there are currently no suitable tools for overlap primer design. Each primer pair has to be hand-curated and validated. Here, 49, 50 and 59 primer pairs were designed to amplify IRS1132 in Ciherang-Sub1, IR64, and pMUGW5, respectively, and the total cost, excluding failed primers, was €870. As an additional and complementary test, DNA gel blot using overlapping probes was performed. The assay could detect ≤ 0.25 copies of vector fragment per haploid genome in positive controls but not detected in GE’d lines (Figure S2). We can, however, not exclude that the sensitivity for detecting small insertions is sufficient. Since none of the assays performed were 100% reliable, we explored whether WGS might be a superior tool for detecting foreign DNA/transgenes.

### WGS for monitoring segregation and detection of transgenes

Initially, we used Illumina technology for WGS. The standard approach uses enzymatic fragmentation. Since enzymatic digestion of genomic DNA is not perfectly unspecific, it is conceivable that insertions will not be detected for a fragment that remains undigested. We therefore performed two independent sets of WGS using different DNA fragmentation methods (enzymatic digestion and mechanical shearing via sonication). Genome assembly and screening for vector backbone with the sequences from the two independent sets of WGS detected *Cas9* fragment insertion in IR64-7a, and recurring singletons in IR64-134a, IR64-134d, CS-1b, CS-1h, and CS-6d. These lines were subsequently not considered as a potential vector-free GE’d lines for export to target countries. The two protocols for short-read WGS also detected false positive hits for vector integration, potentially due to index-hopping or from contaminations, e.g., pollen from transgene-containing plants grown in parallel (Li *et al*., 2019). Reads mapping to the vector were underrepresented and not consistently detected in all biological samples, supporting an origin from index hopping or contamination. Thus, as an improved measure, we grew plants individually in sterile culture and performed long-read WGS without barcoding, providing us with an ultra-sensitive alignment (seed alignment as short as 13 bp). Using long-read WGS, foreign DNA/transgene insertions could not be detected, further supporting the hypothesis that the low copy reads for vector derived from index hopping or contaminations (e.g., pollen from other plants in the greenhouse). It may be useful to evaluate both coverage depth and breadth (i.e., uniformity of genome sequence coverage) in WGS, since focusing only on the average sequencing depth can obscure blind spots in coverage. Among the tested library preparations, HIFI-PacBio has the highest and most consistent breadth coverage even at moderate sequencing depths (∼30×). Standard Illumina sequencing and NanoBall sequencing of enzymatically digested or mechanically sheared DNA samples exhibited minimal differences in sequencing breadth coverage (Figure S8). One may thus consider using long-read PacBio HiFi sequencing (cost ∼750€ per rice line) combined with sensitive mapping parameters that allow the detection of short sequence matches. Long-read sequencing provided the highest confidence for detecting T-DNA or vector backbone fragment insertions.

### SNPs and off-target mutations

An additional advantage of WGS is the ability to screen for potential mutations (naturally occurring mutations that manifest in each generation, potential off-target and somaclonal mutations), and to predict whether such mutations might impact performance. Here, we set a threshold of five mismatches for a stringent screen for potential off-target mutations in the GE’d lines. If we allow three mismatches, the single base insertion in position 6314414 on chromosome 1 (in IR64-9a, IR64-136a, and IR64-7b) and position 15429591 in chromosome 5 (in IR64-136a) could be potential off-target mutations that might have been caused by Cas9 by editing gRNA targeting the EBE for PthXo2 in three independent lines: IR64-9a, IR64-136a, and IR64-7b (Table S4). While it cannot be excluded that the SNP traced to intergenic regions may have epigenetic effects, it is unlikely to directly affect the function of a gene. We traced SNPs across five generations in three GE’d lineages by comparing their genomes to the WT genome (Figure 3, Figure S4). The sum of SNPs was in the range of 100 - 300 for GE’d lines in the respective elite backgrounds, which is crudely in the range of the expected natural mutation rate after 5-7 generations. Notably, more than 75% of the SNPs found in GE’d lines were intergenic and up to 24% were located in the intron, with less than 10% of the SNPs in exons. The presence of high-density SNPs in chromosomes 8, 11, and 12 where the EBEs were located, across three generations of GE’d line, suggests linkage drag. However, severe traits linked to the edits introduced in the GE’d line were not observed. The lack of agronomic penalties from the GE’d line pointed to the non-detrimental effects of the exonic SNPs and indels (Figure 4, Tables S6-10). GE’d lines that underwent and passed the approaches used for transgene analysis have been transferred to KALRO, Kenya for seed amplification, field trials in the presence of BB, and potential registration as new breeding material (Figure 1). The transfer required the execution of a Material Transfer Agreement (MTA) and a plant health inspection by the German Chamber of Agriculture to provide a plant health certificate that declares the plants to be free from pests (www.landwirtschaftskammer.de/landwirtschaft/pflanzenschutz/psd/index.htm). Once suitable agreements are in place for ICAR, the edited lines that underwent and passed the transgene analysis protocol could be transferred to India. Transfer of the lines to farmers may require freedom to operate (evaluation of possible infringement of intellectual property). We transferred the lines for humanitarian purposes and do not request royalties from KALRO or ICAR. The lines can also be transferred to other countries that have appropriate regulations for GE’d crops after the execution of the MTA.

### Results for GE’d lines in multi-season and multi-location EFPs

To evaluate whether the promoter edits, off-target or other unintended mutations might affect the performance of the GE’d IR64 and Ciherang-Sub1lines, we performed multi-year multi-site EFPs in the Philippines and Colombia. EFP conditions tested included wet and dry harvest seasons from 2020 to 2023, and different nitrogen regimes. Multispectral imaging and individual agronomic traits analyses indicate that the GE’d lines behave similarly to unedited parental controls, in particular, the most important trait, i.e. yield was unaffected. It may be noteworthy that GE’d IR64 lines exhibited a reduced 1000-grain weight (reduction of 4 - 8%) compared to unedited IR64 plants, while the 1000-grain weight of GE’d Ciherang-Sub1 lines was not impacted. Similarly, the 1000-grain weight of other GE’d Zhonghua elite rice variety was not impacted (Zeng *et al*., 2020). Compensatory increases in fertility and/or spikelet number may explain why total yield was unaffected. Since the edits introduced in the EBEs of GE’d Ciherang-Sub1 and IR64 lines were comparable, it is unlikely that the effect on 1000-grain weight in the IR64 lines was due to the edits in the promoters or off-target mutations (Table S6-10). The GE’d IR64 lines tested in this study were independent transformants, thus second site mutations are also not likely causes. The most parsimonious explanation might be that the IR64 control acquired natural mutation(s) that impacts 1000-grain weight. Since total yield, which is the trait breeders and farmers are interested in was unchanged, the reduced 1000-grain weight was not considered a concern. In sum, large-scale multi-location and multi-season EFPs with different nitrogen regimes validated that GE’d IR64 and Ciherang-Sub1 cultivars were not affected in agronomic performance compared to respective WT plants.

### Summary

After careful analysis for transgenes/foreign DNA and muti-location multi-year field trials, we identified nine lines in three elite varieties that are suitable for transfer to rice growing countries that have instated regulations for GE’d crops.

## MATERIALS AND METHODS

### DNA gel blots for detecting transgenes

Probes were generated using primers listed in Table S1 and labelled using the PCR DIG probe synthesis kit (Roche, Mannheim, Germany) according to the manufacturer’s instructions. For each sample, 1 mg of genomic DNA was digested with 20 units *BmtI* at 37°C for 1 hr followed by enzyme inactivation at 65°C for 20 mins. Digested genomic DNA were separated via electrophoresis at 90 V for 90 mins using 1% (w/v) UltraPure Agarose (Thermo Fischer Scientific, Waltham, Massachusetts, USA) before blotting onto nylon membrane overnight. DNA crosslinking was performed using UV Stratalynker followed by probe hybridization using the DIG Easy Hybridization kit (Roche, Mannheim, Germany) according to manufacturer’s instructions. Hybridized membrane was washed with low stringency buffer for 5 mins at room temperature followed by washing with high stringency buffer for 15 mins at 70°C. Washed membrane was treated with blocking solution for 30 mins with shaking at 60 rpm and Anti-Digoxigenin-AP antibody for 30 mins with shaking at 60 rpm. The labeled membrane was washed and treated with CDP-Star (Roche, Mannheim, Germany) for signal detection using ImaGE’dQuant LAS4000 at high-binning sensitivity. For analyzing the sensitivity of the probes, a 359 bp Cpf1 probe labeled with digoxygenin (DIG) was used for detecting Cpf1 fragments in genomic DNA of WT Komboka spiked with 0.1 to 3.5 copies of digested pMUGW5 (Figure S7c, Table S1). Copy number of T-DNA insertions was calculated by molar ratio in the genome of Komboka according to https://brcf.medicine.umich.edu/cores/transgenic-animal-model/training-education/protocols/spike/.

### Whole genome sequencing of GE’d lines

The genomes of the GE’d lines were subjected to nanoball sequencing using the T7 platform (DNBSEQ-T7 PE150) (BGI Genomics) (Drmanac et al., 2010). 100 mg tissues were harvested from young leaves of plants grown in greenhouse (GE’d IR64 and Ciherang-Sub1 lines) or MS medium under sterile conditions (GE’d Komboka lines) to avoid cross-contamination from parental lines. Leaves from two plants were pooled as one sample per GE’d line. DNA was extracted from selected lines using DNeasy Plant Mini Kits (Qiagen, Hilde, Germany). The DNA quality was evaluated using Qubit 4 fluorimeter (Thermo Fisher Scientific, Waltham, Massachusetts, USA). Two independent DNA libraries were prepared, i.e., enzymatic and mechanical fragmentation for each line. For enzymatic fragmentation, libraries were prepared with NEBNext Ultra II FS DNA Library Prep with Sample Purification Beads. For mechanical fragmentation, DNA was sonicated with LE220 Focused Ultrasonicator (Covaris LLC, US) using default parameters for 400-500 bp fragments. DNA fragment size was validated with DNF-474 high sensitivity NGS fragment analysis kit. NEBNext® Ultra™ II DNA Library Prep Kit (Germany) for Illumina was used to generate the libraries. Barcoding was done using NEBNext® Multiplex Oligos for Illumina® (96 Unique Dual Index Primer Pairs).

### WGS for reference genome from IR64, Ciherang-Sub1 and Komboka

While assembled reference genomes for the rice variety IR64 are readily available (GCA_009914875 and GCA_011764405 submitted by the Arizona State University and Institute of Crop Science-NARO, respectively), 142,456 SNPs were identified between the two IR64 assemblies. To ensure accurate analysis for the EBE edited lines, we performed whole genome sequencing and assembled three reference genomes, i.e., IR64, Ciherang-Sub1, and Komboka. The genomes were sequenced using Oxford Nanopore Technology (ONT) R9 and R10. Read calling was performed with the dorado tool V0.5 (https://github.com/nanoporetech/dorado). Adapters were removed using porechop V0.2 (https://github.com/rrwick/Porechop). Filtering and cleaning processes were conducted with filtlong V0.2 (https://github.com/rrwick/Filtlong). Filtered and corrected long reads were assembled with the NECAT V0.01 assembler (Ying Chen *et al*., 2021). The resulting contigs were inspected and corrected using the Inspector tool (Yu Chen *et al*., 2021), then polished with High quality (PHRED score > 30) Illumina reads using NextPolish v1.4 (https://github.com/Nextomics/NextPolish). Finally, the contigs were ordered and scaffolded with the Ragtag tool V2.1 (Alonge *et al*., 2022), guided by the indica cultivar IR64 (IRRI) assembly (GenBank accession GCA_009914875). This process yielded a pseudomolecule representing the chromosomes of IR64, Ciherang-Sub1 and Komboka. The assembly for IR64 yielded 1,508 contigs with a total length of 368,972,140 bp and a N_50_ of 415,664 bp whereas 124 contigs with a total length of 393,489,778 bp and N_50_ of 13,319,595 bp for Ciherang-sub-1. We provide the full assembly of IR64, Ciherang-Sub1, and Komboka with the accession numbers in Table S12. Analysis of assembly completeness using BUSCO V1.0 (doi: 10.1002/cpz1.323. PMID: 34936221.) indicated that more than 97 % of the core genes in *Poales* and between 95-98% from GCA_009914875 were recovered in the genome assembly of IR64 and Ciherang-Sub1, indicating a good completeness (Wingett and Andrews, 2018)(Martin, 2011).

### Detection of vector integration in GE’d lines

For each sequence sample, filtered and cleaned reads were mapped (aligned) against the vector sequence reference IRS11345 using the Bowtie2 V1.21 (doi: 10.1186/gb-2009-10-3-r25), for IR64 and Chiherang-Sub1 samples. The samtools V1.19 and picardTools packages V3.2 (Danecek *et al*., 2021) (https://broadinstitute.github.io/picard/) were used to post-process aligned reads, sort for optical duplicates, sort by coordinate, and recall mapped reads. A seed sequence of 19 bases was used for screening of vector in lines sequenced with the Illumina (short-read) platform and a seed sequence of 13 for genomes sequenced with PacBio (long-read) platform. We also conducted sequence alignments of IRS1132 with the WT rice lines to identify identical regions between the vectors and the WT lines that could cause confounding results due to the use of native rice promoters in gene expression constructs in the vectors. The alignment tool blastn (doi: 10.1093/nar/25.17.3389. PMID: 9254694; PMCID: PMC146917) was used to identify identical regions between vector sequences and rice genome sequences. The Artemis genome browser V18.2 (Carver *et al*., 2008) was used to: 1) inspect visually the vector sequence and the bam file containing the aligned reads; 2) load the areas between the vector and the genomes of the rice lines; and 3) inspect the covered regions of the vector and detect integration events.

### Detection of off-target mutations in edited lines

Detection of potential off-target mutation was conducted following previously reported protocols with some modifications (Liu *et al*., 2021). Briefly Reads were filtered by Phred score using TrimGalore V0.6. Alignments were performed using Bowtie2 V1.2 (doi: 10.1186/gb-2009-10-3-r25), following by a Indel Realignment with GATK3 V3.8 SNVs were detected using in-house reference genome assemblies with the sequence variation callers, Loqfreq V2.1, Varscan V2.3, Mutect2-GATK V4.0) and pindel V0.2 (Chen et al., 2020; Koboldt et al., 2009; Wilm et al., 2012; Ye et al., 2009). Consensus SNV calls were obtained using bedtools V2.30 (doi: 10.1093/bioinformatics/btq033), generating vcf files for the agreement of the three SNV callers. Potential off-target predictions were conducted using the Cas-OFFinder tool V2.4 (Bae et al., 2014), allowing for up to 5 mismatches. The Artemis genome browser V18.2 was employed to visually inspect predicted off-targets. Additionally, SNVs were recorded and annotated using SnpEff V5.2 (Cingolani et al., 2012).

### Overlap PCR

Genomic DNA from rice leaves and plasmid vector propagated in *E. coli* were extracted using the DNeasy Plant Mini Kit (Qiagen, Hilden, Germany) and NucleoSpin Plasmid mini kit for plasmid DNA (Machery Nagel, Düren, Germany), respectively, according to the manufacturer’s instructions. PCR was performed with GoTaq G2 Green Master Mix (Promega, Madison, Wisconsin, USA) using primers with their corresponding annealing temperatures as listed in Table S1. The thermocycler was set to 98°C for 5 mins; 25 cycles of 94°C for 30 s, annealing for 30 s, and 72°C for 30 s; and one cycle of 72°C for 2 mins. Electrophoresis was performed on 1% (w/v) agarose in TBA buffer. Sizes of amplicons were determined using GeneRuler 1 kb Plus DNA ladder (Thermo Fischer Scientific, Waltham, Massachusetts, USA).

### Agronomic traits from multi-environmental experimental field plots

For India, IR64-21 (single plant selection from IR64, IRGC 117268) and Ciherang-Sub1 (IRGC 141193) served as both control and parental line for editing (Mackill and Khush, 2018). The edited rice lines had been described (Eom *et al*., 2019; Oliva *et al*., 2019). For Kenya, Komboka (IRGC 141192) was used both as control and parental line for editing (Kitilu *et al*., 2019). Edited Komboka lines have been described (Schepler-Luu *et al*., 2023).

To evaluate the agronomic performance of IR64 and Ciherang-Sub1 with edited EBE lines, three experimental field plots (EFPs) were conducted in two geographically separated environments (CIAT in Colombia, and IRRI in the Philippines). Two EFPs were conducted at CIAT over a dry season from June 2020 to October 2020, and a wet season from January 2021 to May 2021. An EPF was conducted at IRRI during the wet season from December 2020 to March 2021. Each EFP was performed using complete randomized block design with three to four replicates. The plots were 3.96 m^2^ with 20 cm between hills and 25 cm between rows. At the four-leaf stage, (approximately 21 days after sowing), each seedling was transplanted from soil trays onto a hill. In total, 120 plants were planted in each plot. The plants were treated with 60 kg/ha of nitrogen two-, ten-, and thirty-days post-transplant. Integrated agronomic practices were adopted to control pests and weeds throughout the trials. Seven plants per replicate from each EFP were sampled for agronomic performance. Plant height, tiller number, panicle number, and panicle length were quantified. Tissues were dried at 60°C for four days to obtain dry biomass. Grain moisture was adjusted to 14% prior to quantification of single plant grain weight and 100 or 1000 grain weight. Field grain yield (kg/ha) was calculated based on the yield from a single plant multiplied by plant density in the plot. All statistical analyses were performed using the Statistical Analysis System SAS 9.4 (SAS Institute Inc., Cary, NC, USA). ANOVA was performed using a Linear mixed model and further verified with Dunnett’s Multiple Comparison. For nitrogen use efficiency, the EFP was conducted according to (Selvaraj *et al*., 2020). The experimental design utilized a randomized block design, incorporating three replications to facilitate the evaluation of three distinct nitrogen treatment levels. The plants were not treated with 0 kg N ha^-1^ nitrogen fertilizers (0%), 90 kg N ha^-1^ (50%), or 180 kg N ha^-1^ (100%). Drone-based crop monitoring in paddy field experiments was conducted using vegetation indices (VIs) according to protocols by Selvaraj et al. (2020). A DJI MATRICE 600 drone equipped with a MicaSense Altum multispectral camera (MicaSense, Seattle, Washington, USA) captured aerial images. Ground control points (GCPs) were placed at the edges of each plot for accurate georeferencing. Regions of interest (ROIs) were identified and marked using QGIS software (https://www.qgis.org). The vegetation indices, NDVI and NDRE, were analyzed using the CIAT’s Pheno-i platform (http://pheno-i.ciat.cgiar.org). NDVI and NDRE were measured in one- to two-week intervals between 37 and 89 days after planting (DAP).

## Acknowledgements

This work was supported in parts by the Gates Foundation to HHU (investment ID INV-008733), Deutsche Forschungsgemeinschaft (DFG, German Research Foundation) under Germanýs Excellence Strategy – EXC-2048/1 – project ID 390686111 (CEPLAS), and an Alexander von Humboldt Professorship (WF). We dedicate this work to the late Umar Singh, who dedicated his career to helping small-scale producers by developing improved varieties, and who supported us tremendously as a member of the SAB of Healthy Crops.

## Conflict of interest

The authors declare no conflict of interest.

## Author contributions

WBF, BY, FW, BS developed the concept. PC, MS, JT performed EFPs; JHT, TH, VSL performed WGS analyses; EL, MSt., BK, SVG performed transgene analysis; EL, MIRS compiled data; EL, WBF, FW, BY, MB have written the manuscript. All authors have given approval to the final version of the manuscript.

## Notes

### Competing Interest Statement

The authors have declared no competing interest.

