## Supplementary for "From lab to field: analyses of genome-edited bacterial blight resistant rice"

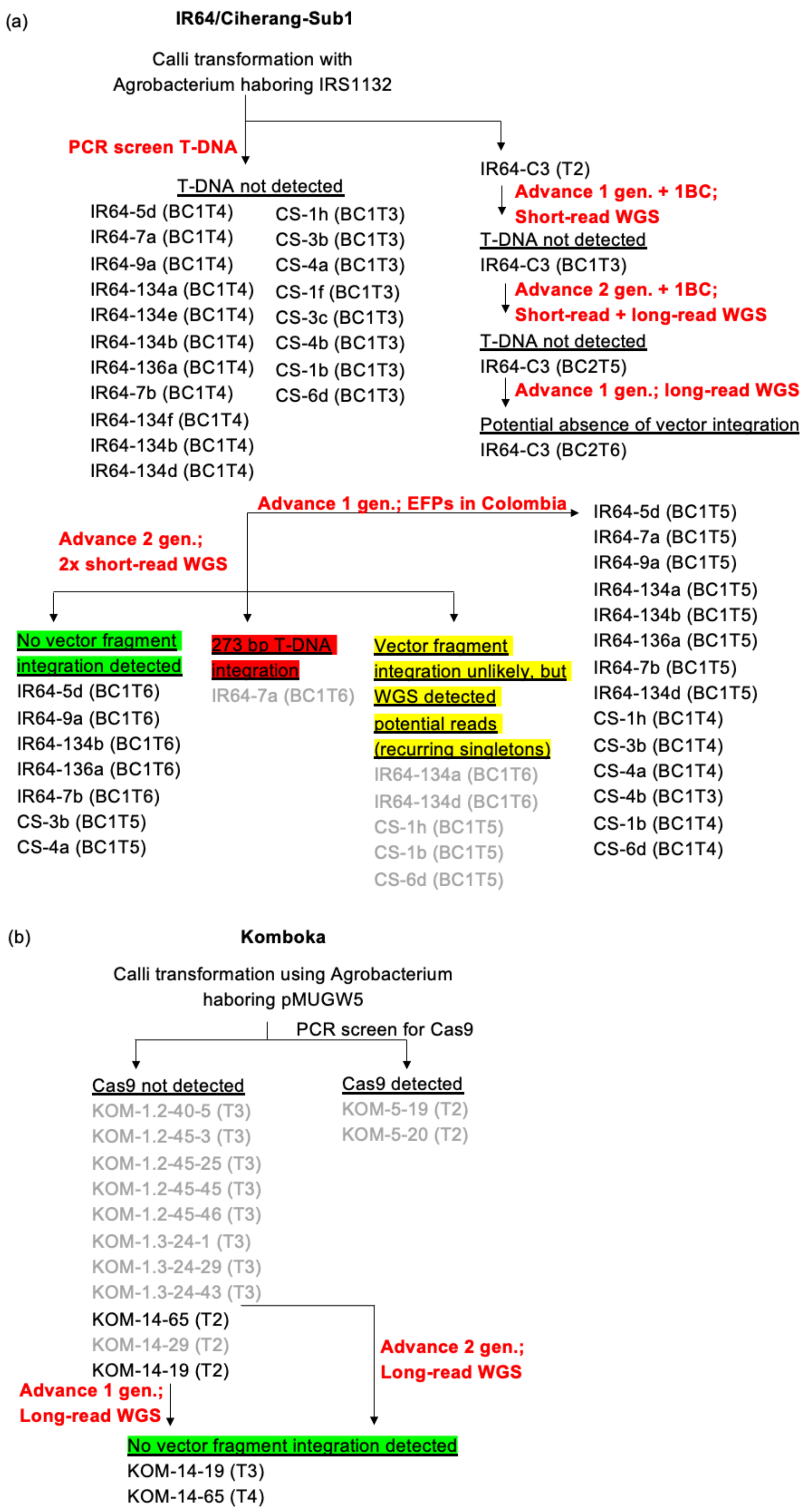


Figure S1: Overview of tests performed in different generations of (a) GE'd IR64 and Ciherang-Sub1 lines; and B) Komboka lines. GE'd lines in grey indicate lines not further pursued for export to interested partners.

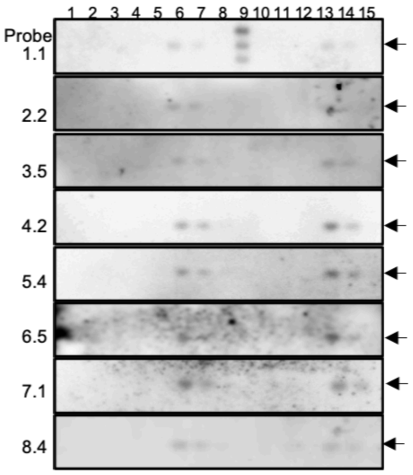


| Lane | Line ID |
|---|---|
| 1 | IR64-9a |
| 2 | IR64-136a |
| 3 | IR64-C3 |
| 4 | IR64-H12 |
| 5 | IR64 WT |
| 6 | IR64 WT + 12 pg IRS1132 |
| 7 | IR64 WT + 6 pg IRS1132 |
| 8 | IR64 WT + 1.2 pg IRS1132 |

| Lane | Line ID |
|---|---|
| 9 | DIG ladder |
| 10 | CS-3b |
| 11 | CS-4 |
| 12 | CS WT |
| 13 | CS WT + 12 pg IRS1132 |
| 14 | CS WT + 6 pg IRS1132 |
| 15 | CS WT + 1.2 pg IRS1132 |

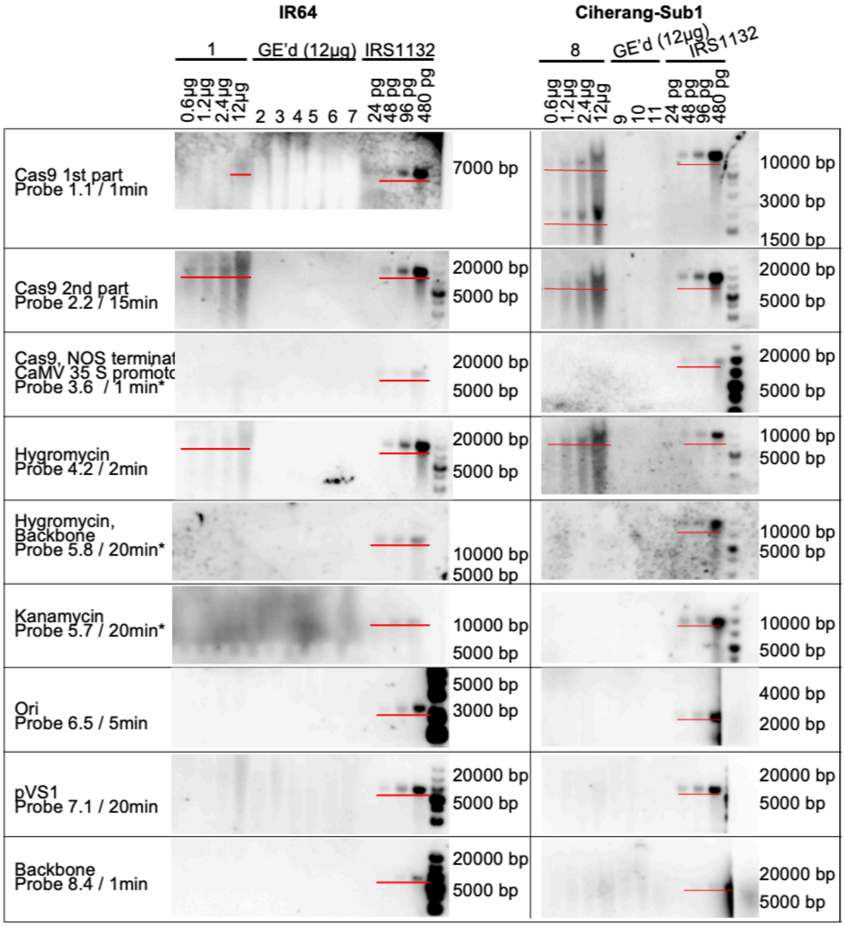


| Lane | Line ID |
|---|---|
| 1 | IR64-136b |
| 2 | IR64-5d |
| 3 | IR64-7b |
| 4 | IR64-9a |
| 5 | IR64-136a |
| 6 | IR64-C3 |
| 7 | IR64 WT |
| 8 | CS-3b |
| 9 | CS-3b |
| 10 | CS-4 |
| 11 | CS WT |

Figure S2: DNA gel blot with overlapping probe for the detection of transgene in GE'd lines. DNA gel blot of GE'd IR64 and Ciherang-Sub1 lines using the overlapping probes (Table S1). IR64-136b and CS-3b are GE'd parental lines with vector integration as positive controls for GE'd lines of respective cultivars. IRS1132 indicate the vector (positive) control. Arrows and red lines indicate fragment of expected size. DNA samples used for each lane in the DNA gel blots indicated in table on the right of the blots.

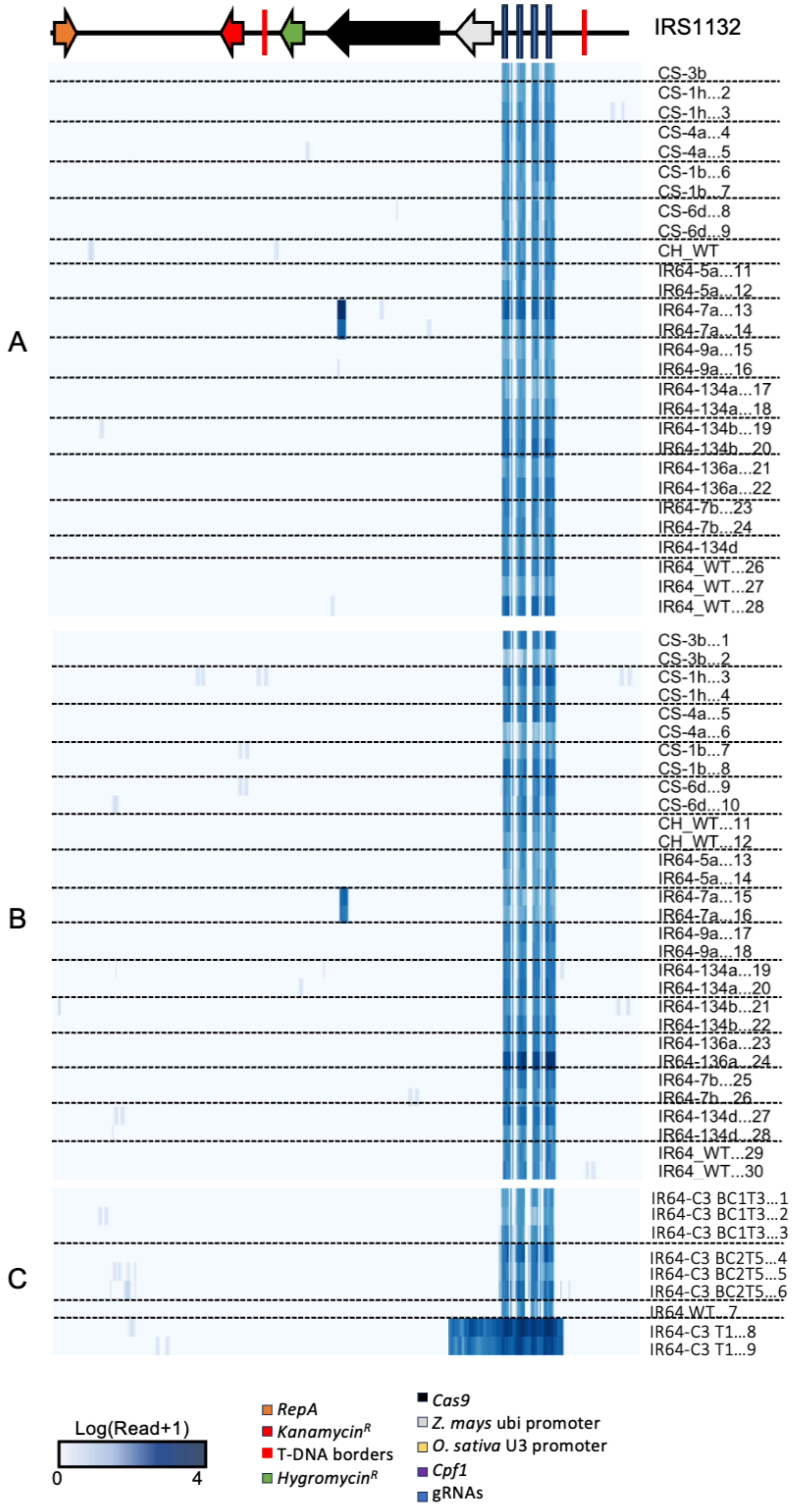


Figure S3: Analysis for the presence of transgenes in GE'd lines.Genome sequence of WT and GE'd IR64 and Ciherang-Sub1 lines against vector IRS1132. Linearized map of each vector indicated at the top genome alignment. Letters on right indicate protocol used for WGS: A- short-read WGS with enzymatic digestion for library preparation, B- short-read WGS with sonication for library preparation, C- short-read WGS using standard Illumina library preparation. Numbers on the right of heatmap indicate ID of GE'd lines and sample number.

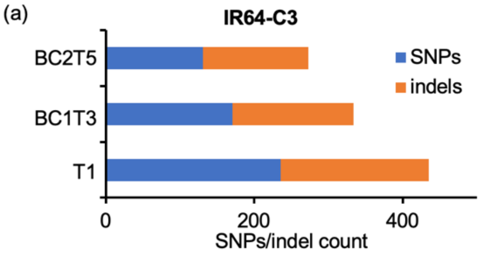


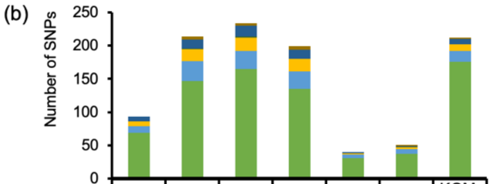


| | IR64-C3 | IR64-9a | IR64-136a | IR64-7b | CS-3b | CS-4 | KOM-14-19-2 |
|---|---|---|---|---|---|---|---|
| Silent | 0 | 5 | 4 | 5 | 0 | 2 | 2 |
| Missense | 7 | 13 | 16 | 14 | 2 | 1 | 8 |
| Splice site | 0 | 1 | 2 | 0 | 0 | 1 | 0 |
| Exon | 7 | 18 | 20 | 19 | 2 | 3 | 10 |
| Intron | 10 | 30 | 27 | 26 | 5 | 7 | 16 |
| Intergenic | 69 | 147 | 165 | 135 | 31 | 37 | 176 |

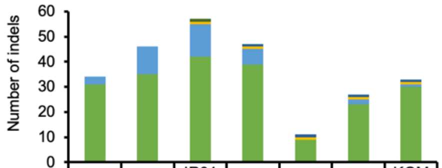


| | IR64-C3 | IR64-9a | IR64-136a | IR64-7b | CS-3b | CS-4 | KOM-14-19-2 |
|---|---|---|---|---|---|---|---|
| Frameshift | 0 | 0 | 0 | 1 | 1 | 1 | 1 |
| In-frame | 0 | 0 | 1 | 0 | 0 | 0 | 0 |
| Exon | 0 | 0 | 1 | 1 | 1 | 1 | 1 |
| Intron | 3 | 11 | 13 | 6 | 0 | 2 | 1 |
| Intergenic | 31 | 35 | 42 | 39 | 9 | 23 | 30 |

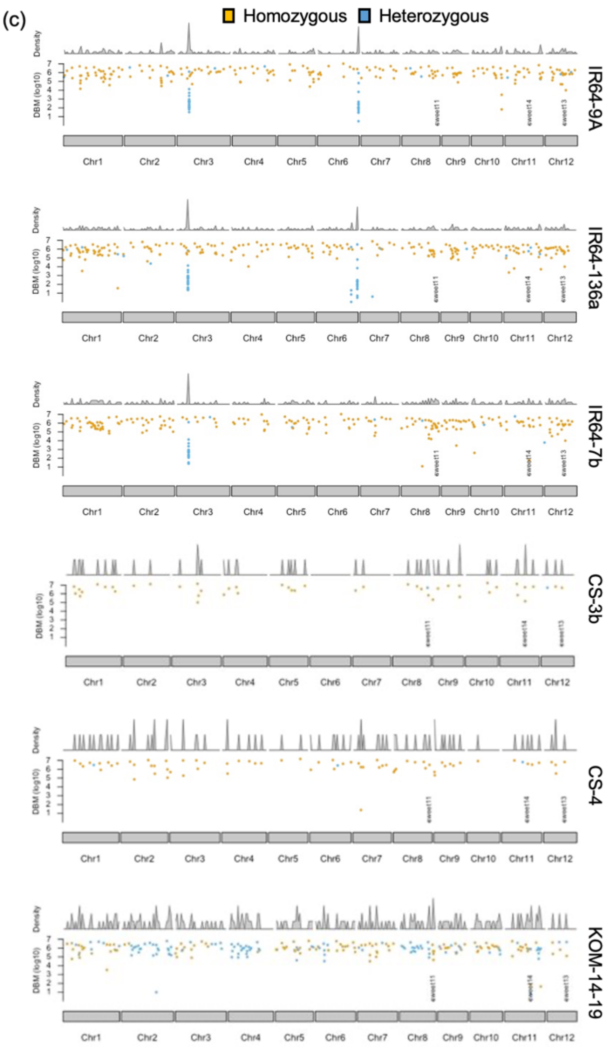


Figure S4: GE'd lines do not carry detrimental indels or SNPs. (a)Number of SNPs and indels found in three generations of GE'd IR64-C3. (b) SNPs and indels detected in the consensus genome sequence of indicated GE'd lines. Exact number of SNPs/indels indicated in the table below bar plots. (c) Density and distribution of SNPs and indels detected in each chromosome of GE'd line. Peaks above the dotplots indicate the density of SNPs/indels. Color of the dots indicate heterogenous SNPs (blue), or homogenous SNPs (yellow). Position of SNPs (chromosome) indicate on the X-axis.

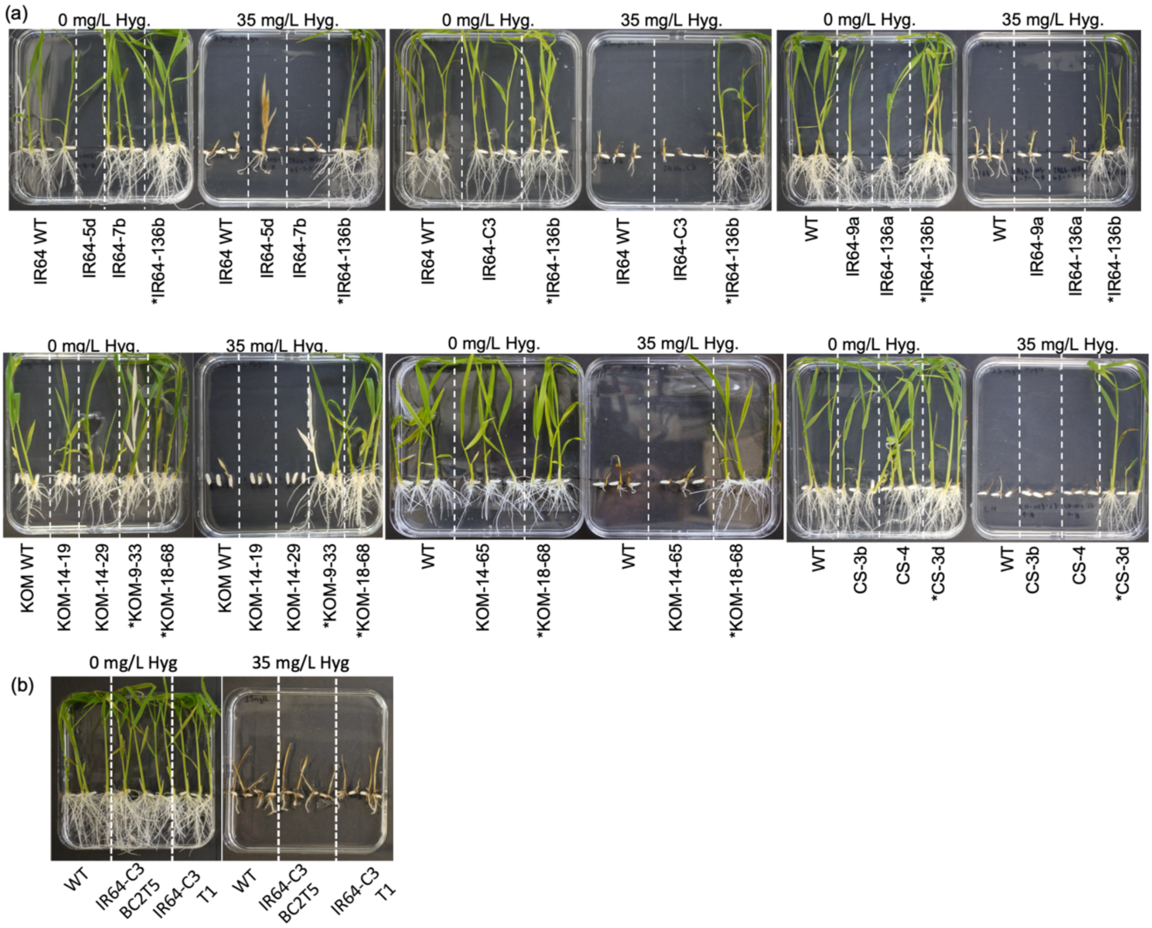


Figure S5: Absence of herbicide tolerance in GE'd lines. (a) Hygromycin (Hyg.) test on GE'd, parental (positive control, indicated with *), and WT plants (negative control). (b) Hygromycin-tolerant gene silencing resulted in loss of hygromycin tolerance in transgene-positive GE'd IR64-C3 T1. WT: wild-type plant, IR64-C3 BC2T5: GE'd IR64 without vector integration, IR64-C3 T1: GE'd IR64 with carrying T-DNA. (c) Gel electrophoresis images for overlap PCR on GE'd IR64 for the detection of vector IRS1132. (d) Gel electrophoresis images for overlap PCR on GE'd Ciherang-Sub1 for the detection of vector IRS1132. (e) Gel electrophoresis images for overlap PCR on GE'd Komboka for the detection of vector pMUGW5.

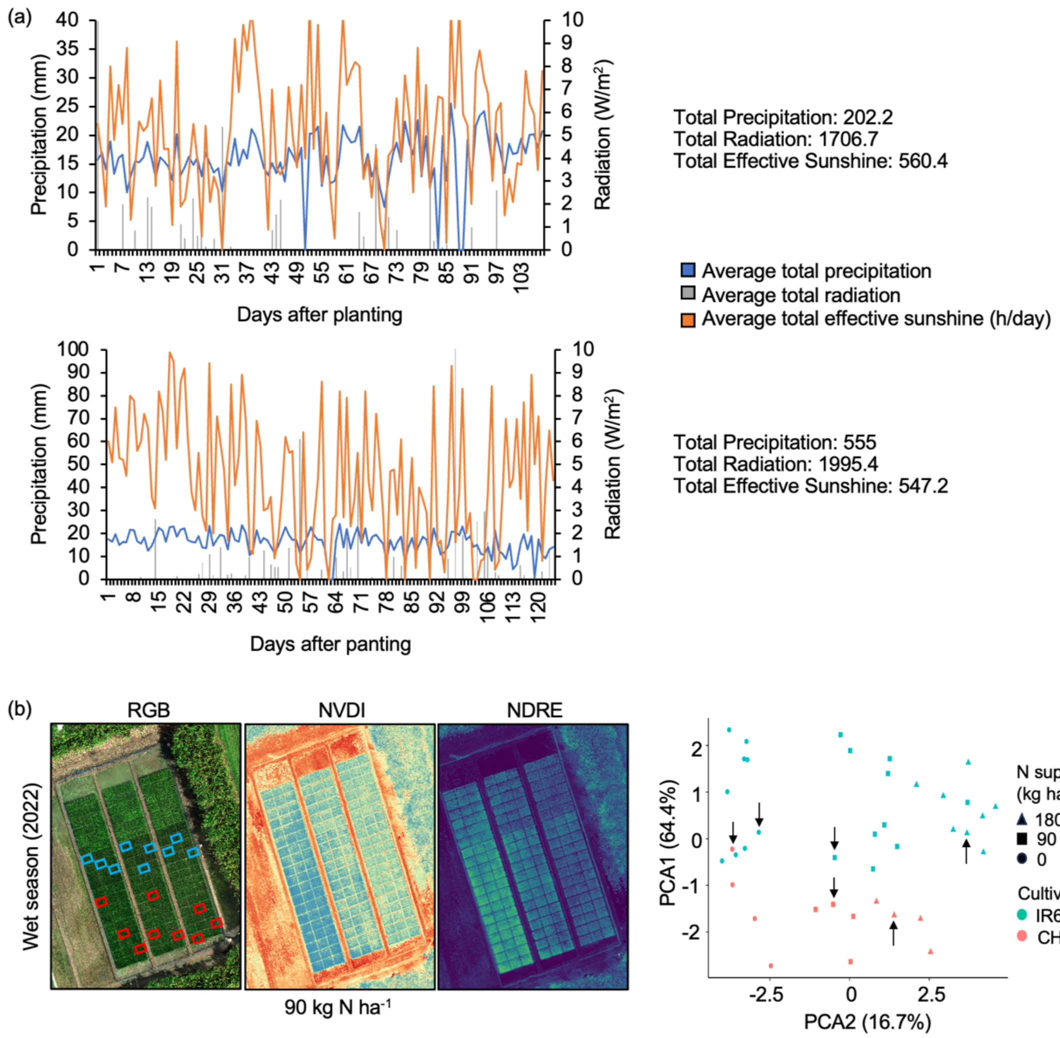


Figure S6: Metreological data during EFPs and agronomic performance of GE'd lines in EFPs. (a) Meteorological information from the CIAT, Colombia June 2020 dry season (top) and January 2021 rainy season (bottom) of EFPs. (b) RGB and multispectral images of the EFPs conducted in Colombia in the wet season of 2021 and its corresponding PCA analysis of EBE-edited and WT IR64 and Ciherang-Sub1 lines under 0%, 50%, and 100% nitrogen treatment. Arrows indicate datapoint for WT plants. Boxed in red indicate WT IR64 plants, boxed in blue indicates WT Ciherang-Sub1 plants.

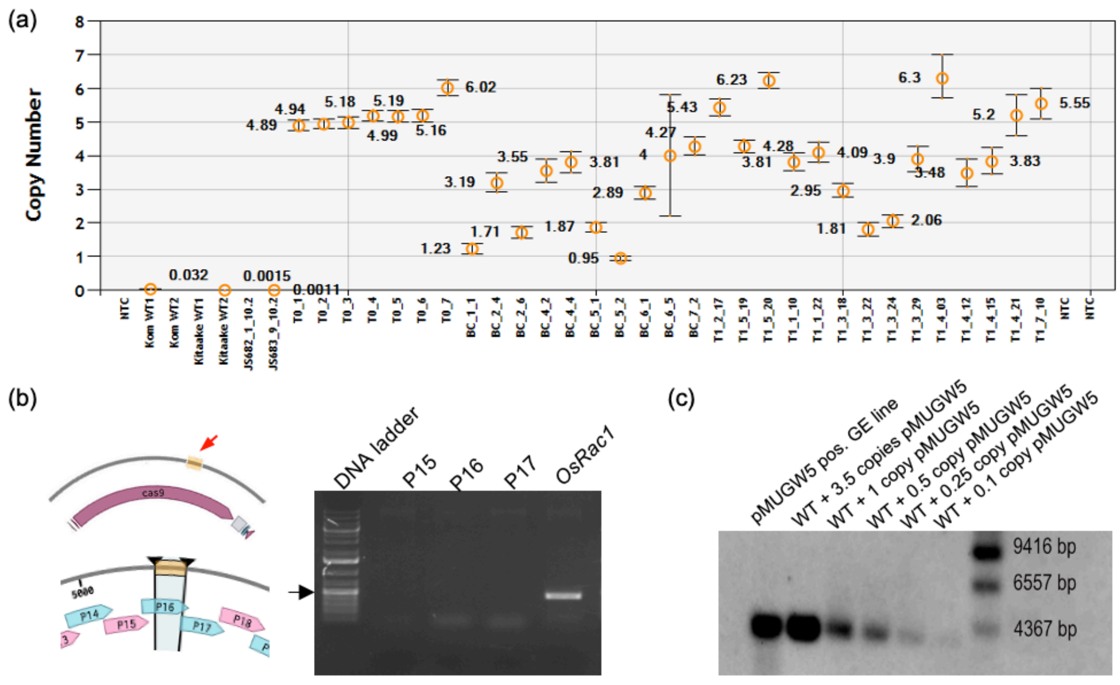


Figure S7: Evaluation of the efficiency of transgene analysis. (a) Determination of Cas9 copy number in T0, T1, and BC1T1 GE'd Komboka via ddPCR. NTC- No template control, Komboka (Kom) and Kitaake WT as negative controls. (b) Overlap PCR failed to detect the presence of Cas9 fragment integrated into the plant's genome. Red arrow and region highlighted in orange indicates the 289 bp fragment from Cas9 incorporated into the genome of IR64-7a. Blue and pink arrows indicate the amplicon produced by indicated primer pairs. Gel electrophoresis image of PCR amplification of IR64-7a using P15, P16, and P17 primers (see scheme on left). *OsRac1* was used as a positive control. Black arrow indicates DNA ladder position for 500 bp. (c) Cpf1-positive transgenic Komboka and WT Komboka spiked with dilutions of pMUGW5 vector blot against DIG-labeled Cpf1 probe.

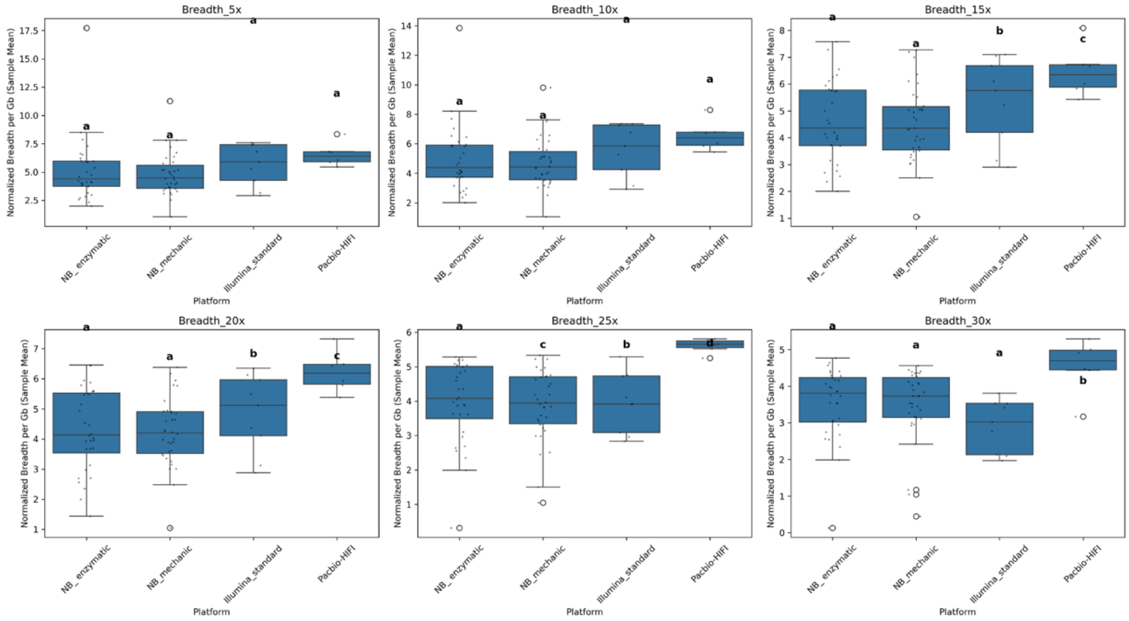


Figure S8: Evaluation of the coverage depth and breadth for the various protocols used for WGS of GE'd lines. An analysis of genome coverage uniformity for each WGS library. The fraction of bases meeting or exceeding certain depth thresholds (5x, 10x, 15x, 20x, 25x, and 30x) indicate the coverage breadth. NB_enzymatic: enzymatically digested genomic DNA using Nanoball platform; NB_mechanic: sonicated genomic DNA using Nanoball platform; Illumina_standard: standard Illumina sequencing library preparation and sequenced using Illumina platform; Pacbio-Hifi: long-read WGS using PacBio platform.

**WGS vector integration**

WGS on DNA samples prepared via the enzymatic digest protocol revealed 178 and 74 reads from the two biological replicates of IR64-7a, respectively, that mapped to *Cas9* (Table S2). In the sonication-based WGS, 57 and 42 reads of IR64-7a (from the replicates) mapped to the same region in *Cas9,* indicating true integration of a 272 bp fragment of *Cas9* in the chromosome 12 of IR64-7a. Samples from IR64-134a contained two/one read (from the replicates). IR64-134b showed four reads and IR64-7b seven reads from two different positions that were that were not present in the biological replicates (Table S2).

Since samples were pooled prior to sequencing, we posit that false positives might be caused by index hopping, i.e., an artifact generated by misassignment of reads during demultiplexing (Kircher *et al.*, 2012). For GE'd Ciherang-Sub1 lines prepared via the enzymatic digest protocol, one biological sample of CS-1h exhibited two reads whereas one read (each) from one biological sample of CS-1b, and CS-6d mapped to regions outside the promoter and gRNAs. The same GE'd Ciherang-Sub1 lines exhibited 6 reads (one biological sample in CS-1h), 2 reads (one biological sample in CS-1b), and 2 and 3 reads from two biological samples of CS-6d (Table S2).

To trace the heritance of SNPs (described below), the whole genomes derived from three generations of IR64-C3, i.e., T1, BC1T3, and BC2T5, were sequenced using short-read WGS (enzymatic fragmentation). A 3546 bp fragment comprising the ZmUBI10 promoter and all four gRNAs was detected in IR64-T1, indicating the presence of a partial insertion of T-DNA in the T1 generation (Figure S2). The line was advanced and backcrossed before being subjected to another round of short-read WGS. Vector integration was not detected in the genome of IR64-C3 BC1T3, an observation further validated in IR64-C3 BC2T5.
