## Supplementary material for "From lab to field: analyses of genome-edited bacterial blight resistant rice": Table S1

Table S1: Primers used in this study

| Application | Region amplified | Direction | 5'-3' Sequence | Product size (bp) | Annealing temp (℃) |
|---|---|---|---|---|---|
| Initial transgene-free screening (IRS1132) | 35S promoter | Forward | CAGAAGACCAAAGGGCTATTGA | 728/401 | 50 |
| | | Reverse | GAGATAGATTTGTAGAGAGAG | | |
| | Right border | Forward | AAGGCGGGAAACGACAAT | 536 | 54 |
| | | Reverse | GTAGGCGAGGTGGTACTAAAC | | |
| | gRNA SWEET14 | Forward | GAATGGTCACGATGGATGAT | 250 | 54 |
| | | Reverse | GGGGTTATAAGGTGGGTTCT | | |
| | gRNA SWEET11/13 | Forward | CCTGGAGCCCATAGTCTAGTAA | 248 | 54 |
| | | Reverse | GCTGGGCAATGTGGTACTAA | | |
| | Ubi promoter | Forward | AGTGTGCATGTGTTCTCCT | 1180 | 58 |
| | | Reverse | GGCACACACATACAGATCCA | | |
| | Cas9 5'end | Forward | TGACAGGCACTCTATAAAGAAGA | 444 | 52 |
| | | Reverse | ATAAAAAGCTTGTCAACATCTGA | | |
| | Cas9 mid | Forward | ATGACTAGAAAGTCCGAAGAAAC | 464 | 52 |
| | | Reverse | ATGTCCTCCAGTATATCTTCGTT | | |
| | Cas9 3' end | Forward | GGGTAATGAACTCGCTCTGC | 431 | 53 |
| | | Reverse | TATAGCGCTTTCTGTCTATGGTA | | |
| | Nos Ter | Forward | TCGTTCAAACATTTGGCAAT | 212 | 52 |
| | | Reverse | CCTAGTTTGCGCGCTATATTT | | |
| DNA gel blot | 1.1 | Forward | GACTATAAGGATGACGACGATAAGATGG | 1941 | 55 |
| | | Reverse | CAAAGTCAACACAATGTCCTC | | |
| | 2.2 | Forward | GAGGACATTGTGTTGACTTTG | 1669 | 60 |
| | | Reverse | ACGACCGCTCCATAATCGTGAT | | |
| | 3.5 | Forward | TCACGATTATGGAGCGGTCG | 1969 | 55 |
| | | Reverse | ATCTGCGAAAGCTCGAGAGAGATAGATT | | |
| | 4.2 | Forward | GCAAGTGGATTGATGTGATA | 1210 | 55 |
| | | Reverse | CTATTTCTTTGCCCTCGGACGAGTG | | |
| | 5.4 | Forward | TGTCGGGCGTACACAAATC | 2659 | 60 |
| | | Reverse | CTTTATCTGTTCGGGCAATACGATGCC | | |
| | 6.5 | Forward | ATCAGACCAGTCAACAAGTACAATGGGC | 2330 | 55 |
| | | Reverse | CTAGCTGATTGGATGTACCGCGAGA | | |
| | 7.1 | Forward | CGCGGTACATCCAATCAGCTAGC | 2244 | 60 |
| | | Reverse | TACCGCGCACTTGAG | | |
| | 8.4 | Forward | TGAAAAACGGTTCGTCCTGGCC | 1264 | 60 |
| | | Reverse | AACGTCGTGACTGGGAAAACCC | | |
| Overlap PCR for IRS1132 | 1 | Forward | tcgacggtatcgata | 379 | 55 |
| | | Reverse | tagctctaaaacgctgctgaca | | |
| | 2 | Forward | cgcttgtgtgtatataaacc | 446 | 50 |
| | | Reverse | gtcctttagaccatgtc | | |
| | 3 | Forward | cacacttgtttgaagtgcag | 494 | 57 |
| | | Reverse | caggctggcattatctactcg | | |
| | 4 | Forward | gcctctaaattaagaaaactaaaactc | 497 | 55 |
| | | Reverse | tctatttattacggcgggcga | | |
| | 5 | Forward | ggattcctttcccac | 500 | 55 |
| | | Reverse | gatgacccgacaaacaagtg | | |
| | 6 | Forward | tagccctgccttcatacgcta | 480 | 55 |
| | | Reverse | cctattcttccttcttgtgtacct | | |
| | 7 | Forward | gcactctataaagaagaacctg | 500 | 55 |
| | | Reverse | ggattctcttcaaacagc | | |
| | 8 | Forward | gcttagcgcatccatgatcaaga | 485 | 55 |
| | | Reverse | cgaccacttcctcaaaattc | | |
| | 9 | Forward | tcgcatggatgactagaa | 500 | 55 |
| | | Reverse | caaagtcaacacaatgtcctc | | |
| | 10 | Forward | cgtatcatgacctgctta | 496 | 55 |
| | | Reverse | aatgttttccggcttatgac | | |
| | 11 | Forward | cgagcatatcgctaacttg | 495 | 55 |
| | | Reverse | gagttgcctccaatagttcttc | | |
| | 12 | Forward | gtaagtcggataatgtcccgtcg | 500 | 55 |
| | | Reverse | ctatttcctgctcagatttcgc | | |
| | 13 | Forward | tgagttcgtctatggcgactat | 504 | 55 |
| | | Reverse | acgaccgctccataatcgtgat | | |
| | 14 | Forward | caaagtggagaagggcaagt | 480 | 55 |
| | | Reverse | aataatgttctcggcctgctc | | |
| | 15 | Forward | gcaaatcttgataaggtcct | 440 | 55 |
| | | Reverse | atgtataattgcgggactc | | |
| | 16 | Forward | cggtcttgcgatgattatc | 431 | 55 |
| | | Reverse | atgttggcaagctgctc | | |
| | 17 | Forward | gaaacctgtcgtgccag | 462 | 55 |
| | | Reverse | gagagtgtcgtgctccaccatgttc | | |
| | 18 | Forward | ccacgtcttcaaagcaagtggattgatgtgaaca | 447 | 55 |
| | | Reverse | tagagagagactggtgatt | | |
| | 19 | Forward | gcaagtggattgatgtgata | 452 | 55 |
| | | Reverse | gtcaggctctcgctaaact | | |
| | 20 | Forward | tgtttatcggcactttgcatcg | 462 | 55 |
| | | Reverse | ctgttatgcggccattgtc | | |
| | 21 | Forward | cgatgagctgatgctttgg | 480 | 55 |
| | | Reverse | cgtcggtttccactatcg | | |
| | 22 | Forward | tgtcgggcgtacacaaatc | 420 | 55 |
| | | Reverse | tgattttgtgccgagctgc | | |
| | 23 | Forward | cgtccgcaatgtgttattaagttgtc | 480 | 55 |
| | | Reverse | ctccaattcactgttccttgcattcta | | |
| | 24 | Forward | ccgtcgatactatgttatacg | 463 | 55 |
| | | Reverse | tcatcttcatactcttccgag | | |
| | 25 | Forward | gcactttgaacggcatgatg | 494 | 55 |
| | | Reverse | atcctccctgatcgaccg | | |
| | 26 | Forward | ctttattgatcttgggagaagc | 498 | 55 |
| | | Reverse | ctcagagcgttttgatatcatgct | | |
| | 27 | Forward | ccaaccttgtttctgagca | 496 | 55 |
| | | Reverse | tggttaagtcgagtaagag | | |
| | 28 | Forward | gagtgactagatgacatatc | 500 | 55 |
| | | Reverse | gtaacctgccaaccaaa | | |
| | 29 | Forward | ggtgacgattacctaacaat | 517 | 55 |
| | | Reverse | tgaagccagttaccttcg | | |
| | 30 | Forward | tgcgcgtaatctgctgctt | 509 | 55 |
| | | Reverse | ggactataaagataccaggcg | | |
| | 31 | Forward | gacaggtatccggtaagcg | 393 | 55 |
| | | Reverse | cagtcacgtagcgatagc | | |
| | 32 | Forward | tgcactctcagtacaatctg | 448 | 55 |
| | | Reverse | gacagttatgcacaggc | | |
| | 33 | Forward | ctggtagattgcctgg | 483 | 55 |
| | | Reverse | atcctagtcatgcgcta | | |
| | 34 | Forward | atttgccctagcatctgc | 484 | 55 |
| | | Reverse | tacgaagaaggccaagaac | | |
| | 35 | Forward | ttcgctctttacgatcttgt | 493 | 55 |
| | | Reverse | cgacgagcaaccagatt | | |
| | 36 | Forward | catgatgctgcgactat | 498 | 55 |
| | | Reverse | gctaccagagtaaatgagca | | |
| | 37 | Forward | ttccatgccgcctcctt | 497 | 55 |
| | | Reverse | atcaaaggcacgcgcatc | | |
| | 38 | Forward | gactcaagaatgggca | 492 | 55 |
| | | Reverse | atgaaggttatcgctgt | | |
| | 39 | Forward | gttgcgatggtcgtcttg | 470 | 55 |
| | | Reverse | taccgcgcacttgag | | |
| | 40 | Forward | aatgcgtcctcacggaa | 482 | 55 |
| | | Reverse | aggatgcttgaccaccta | | |
| | 41 | Forward | caatgtccagtaggtcg | 248 | 55 |
| | | Reverse | tcgtccatttgtatgtgc | | |
| | 42 | Forward | tgaacgtcagaagccgac | 338 | 55 |
| | | Reverse | gcttgagcttggatcagattg | | |
| | 43 | Forward | cgccaatatatcctgt | 379 | 55 |
| | | Reverse | ggcttagttcgttcttaagcttggatccagtcgactgaa | | |
| | 44 | Forward | atgggaaccaattcagtcga | 484 | 55 |
| | | Reverse | cctactgtacaccaccaa | | |
| | 45 | Forward | gaacaatgaagaacccac | 483 | 55 |
| | | Reverse | caactcccttcgttcctctc | | |
| | 46 | Forward | agtacgtgcacggtga | 418 | 55 |
| | | Reverse | tagtacacgctcgtcg | | |
| | 47 | Forward | aagaacccaccttataaccccgcgcgctgccgcttgtgttgagggca | 488 | 55 |
| | | Reverse | gctatttctagctctaaaacggttg | | |
| Overlap PCR for pMUGW5 | 1 | Forward | tggtcatagctgtttcctgtgtga | 499 | 55 |
| | | Reverse | ctcgtgggtgggggtccatctt | | |
| | 2 | Forward | ccatcattgcgataaaggaaaggct | 487 | 55 |
| | | Reverse | ccttacgtcagtggagatatcacatc | | |
| | 3 | Forward | aagatggacccccacccacgag | 394 | 62.5 |
| | | Reverse | cgcaggacatatccacgccctc | | |
| | 4 | Forward | acctgatgcagctctcgga | 439 | 60 |
| | | Reverse | acggtgtcgtccatcacagtttgc | | |
| | 5 | Forward | tgcgcgattgctgatccccatg | 413 | 60 |
| | | Reverse | gcgcccaagctgcatcatcgaa | | |
| | 6 | Forward | gtcttgaccaactctatc | 497 | 55 |
| | | Reverse | agacaacttaataacacattgcggacg | | |
| | 7 | Forward | ccgaattaattcggcgttaattcag | 456 | 55 |
| | | Reverse | ctccaattcactgttccttgcattc | | |
| | 8 | Forward | ggctccgtcgatactatgttatacgc | 486 | 55 |
| | | Reverse | gttcatcttcatactcttccgagc | | |
| | 9 | Forward | gcaatctgctcatgagtgagg | 492 | 55 |
| | | Reverse | caggcttgatccccagtaagtc | | |
| | 10 | Forward | tatcggggaagaacagtatgtcg | 497 | 55.7 |
| | | Reverse | ctgcgccttatccggtaac | | |
| | 11 | Forward | cgataagtcgtgtcttacc | 469 | 55 |
| | | Reverse | agcggtatcagctcactcaaa | | |
| | 12 | Forward | acggttcctggccttttgctgg | 409 | 62.5 |
| | | Reverse | tctgacacatgcagctcccgga | | |
| | 13 | Forward | tcccggcatccgcttacagaca | 407 | 55 |
| | | Reverse | gggaaccggtcacacatgtaagt | | |
| | 14 | Forward | cagacagttatgcacaggccagg | 481 | 62.5 |
| | | Reverse | ggcaaggcagaagccagatg | | |
| | 15 | Forward | cgctgccactgcgttcgtagat | 480 | 60 |
| | | Reverse | cccatctaaccgaatccatgaaccg | | |
| | 16 | Forward | gcaacgtgtggacggaa | 470 | 62.5 |
| | | Reverse | ctgggtgatgacctggtggagaagt | | |
| | 17 | Forward | ctttgcggattcgatcagcg | 477 | 62.5 |
| | | Reverse | atgaagcgggtcaactttcag | | |
| | 18 | Forward | ttcagatagcagctcggtaatggtc | 460 | 62.5 |
| | | Reverse | tcacggatggaaggctacaagc | | |
| | 19 | Forward | ctcaagaatgggcagctcgta | 498 | 62.5 |
| | | Reverse | cgcatgaaggttatcgctgtact | | |
| | 20 | Forward | gctagatgggttgcgatggt | 441 | 62.5 |
| | | Reverse | ttccgtgaggacgcattgac | | |
| | 21 | Forward | ctggtgcggtttcatgcttgttc | 473 | 60 |
| | | Reverse | agcacccgcgacctactg | | |
| | 22 | Forward | gaacacggtcaacaccatgc | 487 | 62.5 |
| | | Reverse | gtcttctgaaaacgacatgtcgca | | |
| | 23 | Forward | atgcgactaaaacacgcgacaag | 480 | 62.5 |
| | | Reverse | ctggcgttacccaacttaatcg | | |
| | 24 | Forward | cctcttcgctattacgccag | 484 | 60 |
| | | Reverse | ccctaaaccctaaatggatgtacta | | |
| | 25 | Forward | gagtattttgacaacaggactctacag | 479 | 60 |
| | | Reverse | acggtggagcggaactctc | | |
| | 26 | Forward | tcgacgagtctaacggacac | 474 | 60 |
| | | Reverse | gaagtagaactaccgggccctaa | | |
| | 27 | Forward | ttcaaggtacgccgctc | 445 | 62.5 |
| | | Reverse | caggtagtttgaaacagtattctactccg | | |
| | 28 | Forward | ggttgtgatgatgtggtgtggttg | 498 | 62.5 |
| | | Reverse | tccttatagtccatggccggattcg | | |
| | 29 | Forward | tgctcaccctgttgtttggtgt | 479 | 55 |
| | | Reverse | cgccatctcgttactgaagat | | |
| | 30 | Forward | ccgcaggtacacaagaaggaag | 489 | 55 |
| | | Reverse | tcaccgggaagctgtgcgat | | |
| | 31 | Forward | atgctaaggccattttgtccgc | 478 | 55 |
| | | Reverse | ccaccgtcaatatagccagcat | | |
| | 32 | Forward | gcttgttcgtcagcagctt | 481 | 55 |
| | | Reverse | attggtcatcctttcgatgaagctc | | |
| | 33 | Forward | cgtttcgcatggatgactaga | 491 | 62.5 |
| | | Reverse | gtcaaagtcaacacaatgtcctcc | | |
| | 34 | Forward | cgtatcatgacctgcttaaaattatcaag | 453 | 55 |
| | | Reverse | caacttttaccgtttgaagaatgcc | | |
| | 35 | Forward | tgcacgagcatatcgctaact | 500 | 55 |
| | | Reverse | atgaagaactattggaggcaactc | | |
| | 36 | Forward | ctgacgcgcagcgataaa | 488 | 55 |
| | | Reverse | ctaacgtcgtagaccttatagtcgcc | | |
| | 37 | Forward | ctaagctcgaatctgagttcgtc | 511 | 55 |
| | | Reverse | gaccgctccataatcgtgatc | | |
| | 38 | Forward | tgcgtactccgtgcttgtc | 481 | 60 |
| | | Reverse | gtctctatgcttgttgtaggcg | | |
| | 39 | Forward | tagcaagcgcgttatactggcag | 474 | 55 |
| | | Reverse | cccatctcataaataacgtcatgc | | |
| | 40 | Forward | atcgacctttcgcagttg | 472 | 55 |
| | | Reverse | cttcgaagggcaattctg | | |
| | 41 | Forward | gcttatcgataccgtcg | 488 | 55 |
| | | Reverse | agatttccaaccaggtcc | | |
| | 42 | Forward | gcaattccagcacactg | 493 | 55 |
| | | Reverse | aagaacgaactaagccggac | | |
| | 43 | Forward | cccatcatccatcgtgacca | 499 | 55 |
| | | Reverse | tctgtgccaagtgtgagctgc | | |
| | 44 | Forward | ggcagtagtagcttgatgcct | 558 | 62.5 |
| | | Reverse | agcaaattactgatgagtccgtg | | |
| | 45 | Forward | tgccgaagcatgttgcc | 561 | 62.5 |
| | | Reverse | ggctgggcaacatgctt | | |
| | 46 | Forward | cacaaattatcagcacgctcac | 592 | 62.5 |
| | | Reverse | gcatggatcttggaggaatcagatg | | |
| | 47 | Forward | gtagcaccagtagaagacgc | 307 | 55 |
| | | Reverse | attactgatgagtccgtgagg | | |
| | 48 | Forward | ctcatctcccccactgta | 496 | 55.7 |
| | | Reverse | atgcaagctagagacgg | | |
| | 49 | Forward | ctgtgtgaagattattgccact | 481 | 55 |
| | | Reverse | acaggctgatgtaattgttc | | |
| | 50 | Forward | atcaacgacgtgctgcacag | 468 | 62.5 |
| | | Reverse | atcttctccttgatctcctgc | | |
| | 51 | Forward | ctaatatggacatcttcgagaagg | 497 | 55 |
| | | Reverse | atccttggagattgtgctgatg | | |
| | 52 | Forward | tactctagcgccggcatctt | 520 | 62.5 |
| | | Reverse | gatgtcgtaggccagcacaa | | |
| | 53 | Forward | ggcaaggagacaaacagggac | 497 | 55 |
| | | Reverse | ctatccttaaagaagtcgatcagc | | |
| | 54 | Forward | atggcacattcaagaaggg | 504 | 55 |
| | | Reverse | ccttatacacgtcgtaggaca | | |
| | 55 | Forward | ccaacaagaatccagataatcc | 492 | 55 |
| | | Reverse | cagagttcaggtcctccag | | |
| | 56 | Forward | tatcaaggagctgaaggc | 477 | 55 |
| | | Reverse | cgagagaagttcttatagtccaggg | | |
| | 57 | Forward | acaggatcatgtacgtgc | 485 | 62.5 |
| | | Reverse | ccacagcacctttctgg | | |
| | 58 | Forward | cgatagccggaactatgag | 472 | 55 |
| | | Reverse | ggggaaattcgagctcacta | | |
| | 59 | Forward | gcagatatccagcacagtg | 403 | 55 |
| | | Reverse | tttacactttatgcttccggctcg | | |
