## Supplementary material for "From lab to field: analyses of genome-edited bacterial blight resistant rice": Table S2

Table S2: Whole genome mapping of GE'd lines to vectors IRS1132

| Library preparation | Sample ID | Replicate | Non-singletons | Singletons |
|---|---|---|---|---|
| Enzymatic digestion | IR64-5d | 1 | N.D. | 77 |
| | IR64-5d | 2 | N.D. | N.D. |
| | IR64-7a | 1 | 273 | 132 |
| | IR64-7a | 2 | 273 | 210 |
| | IR64-9a | 1 | N.D. | 64 |
| | IR64-9a | 2 | N.D. | N.D. |
| | IR64-134a | 1 | N.D. | N.D. |
| | IR64-134a | 2 | N.D. | 82 |
| | IR64-134b | 1 | N.D. | N.D. |
| | IR64-134b | 2 | N.D. | N.D. |
| | IR64-136a | 1 | N.D. | N.D. |
| | IR64-136a | 2 | N.D. | N.D. |
| | IR64-7b | 1 | N.D. | N.D. |
| | IR64-7b | 2 | N.D. | N.D. |
| | IR64-134d | 1 | N.D. | N.D. |
| | IR64 WT | 1 | N.D. | N.D. |
| | IR64 WT | 2 | N.D. | N.D. |
| | IR64 WT | 3 | N.D. | N.D. |
| | CS-3b | 1 | N.D. | N.D. |
| | CS-1h | 1 | N.D. | N.D. |
| | CS-1h | 2 | N.D. | 232 |
| | CS-4 | 1 | N.D. | N.D. |
| | CS-4 | 2 | N.D. | N.D. |
| | CS-1b | 1 | N.D. | N.D. |
| | CS-1b | 2 | N.D. | N.D. |
| | CS-6d | 1 | N.D. | 62 |
| | CS-6d | 2 | N.D. | 229 |
| | Ciherang-Sub1 WT | 1 | N.D. | N.D. |
| | Ciherang-Sub1 WT | 2 | N.D. | N.D. |
| | Ciherang-Sub1 WT | 3 | N.D. | N.D. |
| Mechanical shearing | IR64-5d | 1 | N.D. | N.D. |
| | IR64-5d | 2 | N.D. | N.D. |
| | IR64-7a | 1 | 273 | N.D. |
| | IR64-7a | 2 | 273 | N.D. |
| | IR64-9a | 1 | N.D. | N.D. |
| | IR64-9a | 2 | N.D. | N.D. |
| | IR64-134a | 1 | N.D. | 224 |
| | IR64-134a | 2 | N.D. | 135 |
| | IR64-134b | 1 | N.D. | 218 |
| | IR64-134b | 2 | N.D. | 135 |
| | IR64-136a | 1 | N.D. | N.D. |
| | IR64-136a | 2 | N.D. | N.D. |
| | IR64-7b | 1 | N.D. | N.D. |
| | IR64-7b | 2 | N.D. | 270 |
| | IR64-134d | 1 | N.D. | 270 |
| | IR64-134d | 2 | N.D. | 44 |
| | IR64 WT | 1 | N.D. | N.D. |
| | IR64 WT | 2 | N.D. | 237 |
| | CS-3b | 1 | N.D. | N.D. |
| | CS-3b | 2 | N.D. | N.D. |
| | CS-1h | 1 | N.D. | 807 |
| | CS-1h | 2 | N.D. | N.D. |
| | CS-4 | 1 | N.D. | N.D. |
| | CS-4 | 2 | N.D. | N.D. |
| | CS-1b | 1 | N.D. | 269 |
| | CS-1b | 2 | N.D. | N.D. |
| | CS-6d | 1 | N.D. | N.D. |
| | CS-6d | 2 | N.D. | 262 |
| Standard Illumina | IR64-C3 T1 | 1 | 3646 | 211 |
| | IR64-C3 T1 | 2 | 3646 | 248 |
| | IR64-C3 BC1T3 | 1 | N.D. | 249 |
| | IR64-C3 BC1T3 | 2 | N.D. | N.D. |
| | IR64-C3 BC1T3 | 3 | N.D. | 230 |
| | IR64-C3 BC2T5 | 1 | N.D. | 257 |
| | IR64-C3 BC2T5 | 2 | N.D. | 692 |
| | IR64-C3 BC2T5 | 3 | N.D. | 676 |
| | IR64 WT | 1 | N.D. | N.D. |

N.D- None detected
