## Supplementary material for "From lab to field: analyses of genome-edited bacterial blight resistant rice": Table S3

**Table S3: Genes harbouring indels in genome of GE'd rice lines**

| GE'd line | GeneBank protein ID | Annotation in rice genome database | Locus name | Position |
|---|---|---|---|---|
| IR64-136a | EEC73301.1 | SNF2 family N-terminal domain containing protein, expressed | LOC_Os02g32570.1 | Chr.2 position 267950-267952 |
| IR64-7b | Not determined | Short hypothetical protein | Not determined | Chr.? position 33191418-33191420 |
| CS-3b | EAY73218.1 | dnaJ domain containing protein, expressed | LOC_Os01g13760.1 | Chr.1 position 7719897-7719899 |
| CS-4 | EAY75494.1 | surp module family protein, putative, expressed | LOC_Os01g50320.2 | Chr.1 position 30040733-30040735 |
| KOM-14-19 | KAF2941743.1 | Aldehyde oxidase 2, putative, expressed | LOC_Os03g57720.1 | Chr.3 position 35212264-35212266 |
