## Supplementary material for "From lab to field: analyses of genome-edited bacterial blight resistant rice": Table S4

Table S4: Off-target mutation analysis

| GE line | Chromosom | Chromosom | SNP/Indel p | Sequence in | Sequence in | Zygousity of | gRNA bindin | gRNA binding pos | gRNA match | DNA strand | Number of mismacth | gRNA | Distance fro | Note |
|---|---|---|---|---|---|---|---|---|---|---|---|---|---|---|
| IR64-C3 | 8 | CM020883.1 | 25660109 | T | TAA | Homozygous | 25660095 | 25660115 | TTGGTGGT | + | 0 | gRNA1 in IRS1132 | 0 | On target |
| | 12 | CM020887.1 | 14149934 | GGAGTTGT | G | Homozygous | 14149923 | 14149943 | GAGGAACG | + | 0 | gRNA2 in IRS1132 | 0 | On target |
| | 11 | CM020886.1 | 17855327 | CTGCTGA | C | Homozygous | 17855323 | 17855343 | AGGGCATG | - | 0 | gRNA3 in IRS1132 | 0 | On target |
| | 11 | CM020886.1 | 17855266 | TTGGAGGG | T | Homozygous | 17855261 | 17855281 | TATATAAAC | - | 0 | gRNA4 in IRS1132 | 0 | On target |
| | 11 | CM020886.1 | 17855266 | TTGGAGGG | T | Homozygous | 17855248 | 17855268 | GATGAGCTT | + | 0 | gRNA6 in IRS1132 | 0 | On target |
| IR64-9a | 8 | CM020883.1 | 25660108 | G | GT | Homozygous | 25660095 | 25660115 | TTGGTGGT | + | 0 | gRNA1 in IRS1132 | 0 | On target |
| | 1 | CM020876.1 | 6314428 | G | GT | Homozygous | 6314414 | 6314434 | GAGGAACG | + | 3 | gRNA2 in IRS1132 | 0 | Off target |
| | 12 | CM020887.1 | 14149937 | G | GT | Homozygous | 14149923 | 14149943 | GAGGAACG | + | 0 | gRNA2 in IRS1132 | 0 | On target |
| | 11 | CM020886.1 | 17855260 | . | . | . | 17855248 | 17855268 | GATGAGCTT | + | 0 | gRNA6 in IRS1132 | 0 | On target |
| | 11 | CM020886.1 | 17855260 | . | . | . | 17855261 | 17855281 | TATATAAAC | - | 0 | gRNA4 in IRS1132 | 1 | On target |
| IR64-136a | 8 | CM020883.1 | 25660106 | . | . | . | 25660095 | 25660115 | TTGGTGGT | + | 0 | gRNA1 in IRS1132 | 0 | On target |
| | 1 | CM020876.1 | 6314428 | G | GT | Homozygous | 6314414 | 6314434 | GAGGAACG | + | 3 | gRNA2 in IRS1132 | 0 | Off target |
| | 5 | CM020880.1 | 15429580 | AGGGAGGC | A | Homozygous | 15429571 | 15429591 | GAGGAACG | + | 3 | gRNA2 in IRS1132 | 0 | Off target |
| | 12 | CM020887.1 | 5385079 | G | GA | Homozygous | 5385065 | 5385085 | GAGGAACG | + | 3 | gRNA2 in IRS1132 | 0 | On target |
| | 12 | CM020887.1 | 14149937 | G | GA | Homozygous | 14149923 | 14149943 | GAGGAACG | + | 0 | gRNA2 in IRS1132 | 0 | On target |
| | 11 | CM020886.1 | 17855266 | T | TGCTGACAT | Heterozygou | 17855261 | 17855281 | TATATAAAC | - | 0 | gRNA4 in IRS1132 | 0 | On target |
| | 11 | CM020886.1 | 17855266 | T | TGCTGACAT | Heterozygou | 17855248 | 17855268 | GATGAGCTT | + | 0 | gRNA6 in IRS1132 | 0 | On target |
| IR64-7b | 8 | CM020883.1 | 25660109 | T | TA | Homozygous | 25660095 | 25660115 | TTGGTGGT | + | 0 | gRNA1 in IRS1132 | 0 | On target |
| | 1 | CM020876.1 | 6314428 | G | GT | Homozygous | 6314414 | 6314434 | GAGGAACG | + | 3 | gRNA2 in IRS1132 | 0 | Off target |
| | 12 | CM020887.1 | 14149937 | G | GTT | Homozygous | 14149923 | 14149943 | GAGGAACG | + | 0 | gRNA2 in IRS1132 | 0 | On target |
| | 11 | CM020886.1 | 17855325 | AGCT | A | Homozygous | 17855323 | 17855343 | AGGGCATG | - | 0 | gRNA3 in IRS1132 | 0 | On target |
| | 11 | CM020886.1 | 17855264 | GGTTGGA | G | Homozygous | 17855261 | 17855281 | TATATAAAC | - | 0 | gRNA4 in IRS1132 | 0 | On target |
| | 11 | CM020886.1 | 17855264 | GGTTGGA | G | Homozygous | 17855248 | 17855268 | GATGAGCTT | + | 0 | gRNA6 in IRS1132 | 0 | On target |
| CS-3b | 8 | CM020883.1 | 28510258 | G | GT | Homozygous | 28510245 | 28510265 | TTGGTGGT | + | 0 | gRNA1 in IRS1132 | 0 | On target |
| | 12 | CM020887.1 | 16544475 | AG | A | Homozygous | 16544462 | 16544482 | GAGGAACG | + | 0 | gRNA2 in IRS1132 | 0 | On target |
| | 11 | CM020886.1 | 20258753 | ATGACCAGC | A | Homozygous | 20258757 | 20258777 | AGGGCATG | - | 0 | gRNA3 in IRS1132 | 4 | On target |
| CS-4 | 8 | CM020883.1 | 28510253 | GTACAGTAC | G | Homozygous | 28510245 | 28510265 | TTGGTGGT | + | 0 | gRNA1 in IRS1132 | 0 | On target |
| | 12 | CM020887.1 | 16544476 | G | GT | Homozygous | 16544462 | 16544482 | GAGGAACG | + | 0 | gRNA2 in IRS1132 | 0 | On target |
| | 11 | CM020886.1 | 20258700 | . | . | . | 20258695 | 20258715 | TATATAAAC | - | 0 | gRNA4 in IRS1132 | 0 | On target |
| | 11 | CM020886.1 | 20258700 | . | . | . | 20258682 | 20258702 | GATGAGCTT | + | 0 | gRNA6 in IRS1132 | 0 | On target |
| KOM-14-19 | 8 | CM020883.1 | 25762485 | AGGGGGAG | A | Homozygous | 25762468 | 25762488 | CATCTCCCCC | - | 0 | gRNA1 in pMUGW5 | 0 | On target |
| | 11 | CM020886.1 | 20025933 | T | TA | Homozygous | 20025931 | 20025951 | GCTGCTGAC | + | 0 | gRNA4 in pMUGW5 | 0 | On target |
| | 11 | CM020886.1 | 20025847 | AGGCTTGAT | A | Heterozygou | 20025841 | 20025861 | GCTAAGCTC | - | 0 | gRNA5 in pMUGW5 | 0 | On target |
| | 11 | CM020886.1 | 20025854 | A | AG | Heterozygou | 20025841 | 20025861 | GCTAAGCTC | - | 0 | gRNA5 in pMUGW5 | 0 | On target |
| | 12 | CM020887.1 | 14966587 | AGGGAGTT | A | Homozygous | 14966589 | 14966609 | GGAGTTGT | + | 0 | gRNA2 in pMUGW5 | 2 | On target |
