## Supplementary material for "From lab to field: analyses of genome-edited bacterial blight resistant rice": Table S5

Table S5: Modification of EBEs across two generations

| EBE | Line ID | Generation |
|---|---|---|
| PthXo1 | IR64-7b | T3 |
| PthXo1 | IR64-7b | T4/BC1 |
| PthXo1 | IR64-9a | T3 |
| PthXo1 | IR64-9a | T4/BC1 |
| PthXo1 | IR64-136a | T3 |
| PthXo1 | IR64-136a | T4/BC1 |
| PthXo1 | IR64-C3 | BC1T3 |
| PthXo1 | IR64-C3 | BC2T5 |
| PthXo1 | CS-3c | T2 (Heterozygous) |
| PthXo1 | CS-3c | T3/BC1 |
| PthXo1 | CS-4a | T2 |
| PthXo1 | CS-4a | T3BC1 |
| PthXo1 | KOM-14-19 | T1 |
| PthXo1 | KOM-14-19 | T3 |
| PthXo1 | KOM-14-65 | T1 |
| PthXo1 | KOM-14-65 | T3 |
| PthXo2 | IR64-7b | T3 |
| PthXo2 | IR64-7b | T4/BC1 |
| PthXo2 | IR64-9a | T3 |
| PthXo2 | IR64-9a | T4/BC1 |
| PthXo2 | IR64-136a | T3 |
| PthXo2 | IR64-136a | T4/BC1 |
| PthXo2 | IR64-C3 | BC1T3 |
| PthXo2 | IR64-C3 | BC2T5 |
| PthXo2 | CS-3c | T2 |
| PthXo2 | CS-3c | T3BC1 |
| PthXo2 | CS-4a | T2 |
| PthXo2 | CS-4a | T3BC1 |
| PthXo2 | KOM-14-19 | T1 |
| PthXo2 | KOM-14-19 | T3 |
| PthXo2 | KOM-14-65 | T1 (Biallelic) |
| PthXo2 | KOM-14-65 | T3 |
| TalF, AvrXa7, PthXo3 | IR64-7b | T3 |
| TalF, AvrXa7, PthXo3 | IR64-7b | T4/BC1 |
| TalF, AvrXa7, PthXo3 | IR64-9a | T3 |
| TalF, AvrXa7, PthXo3 | IR64-9a | T4/BC1 |
| TalF, AvrXa7, PthXo3 | IR64-136a | T3 |
| TalF, AvrXa7, PthXo3 | IR64-136a | T4/BC1 |
| TalF, AvrXa7, PthXo3 | IR64-C3 | BC1T3 |
| TalF, AvrXa7, PthXo3 | IR64-C3 | BC2T5 |
| TalF, AvrXa7, PthXo3 | CS-3c | T2 |
| TalF, AvrXa7, PthXo3 | CS-3c | T3BC1 |
| TalF, AvrXa7, PthXo3 | CS-4a | T2 |
| TalF, AvrXa7, PthXo3 | CS-4a | T3BC1 |
| TalF, AvrXa7, PthXo3 | KOM-14-19 | T1 (Biallelic) |
| TalF, AvrXa7, PthXo3 | KOM-14-19 | T3 |
| TalF, AvrXa7, PthXo3 | KOM-14-65 | T1 (Biallelic) |
| TalF, AvrXa7, PthXo3 | KOM-14-65 | T3 |
