## Supplementary material for "From lab to field: analyses of genome-edited bacterial blight resistant rice": Table S6

Table S6: NDRE and NDVI measurements for EFP in Colobia during wet season of 2020.

| Spectra | Genotype | 48DAP | SE | P-value | 56DAP | SE | P-value | 84DAP | SE | P-value | 87DAP | SE | P-value |
|---|---|---|---|---|---|---|---|---|---|---|---|---|---|
| NDRE | Ciherang-Sub1 WT | 0.483 | 0.024 | | 0.508 | 0.027 | | 0.427 | 0.028 | | 0.366 | 0.032 | |
| | CS-1h | 0.504 | 0.025 | ns | 0.536 | 0.022 | ns | 0.44 | 0.021 | ns | 0.371 | 0.022 | ns |
| | CS-3b | 0.445 | 0.008 | ns | 0.467 | 0.007 | ns | 0.396 | 0.003 | ns | 0.326 | 0.003 | ns |
| | CS-4 | 0.48 | 0.02 | ns | 0.501 | 0.016 | ns | 0.432 | 0.014 | ns | 0.367 | 0.016 | ns |
| | CS-1i | 0.496 | 0.023 | ns | 0.52 | 0.016 | ns | 0.438 | 0.022 | ns | 0.374 | 0.023 | ns |
| | CS-6d | 0.488 | 0.02 | ns | 0.511 | 0.014 | ns | 0.433 | 0.019 | ns | 0.371 | 0.021 | ns |
| | IR 64 WT | 0.453 | 0.018 | | 0.486 | 0.017 | | 0.375 | 0.013 | | 0.302 | 0.007 | |
| | IR64-5d | 0.494 | 0.01 | ns | 0.545 | 0.011 | * | 0.405 | 0.018 | ns | 0.325 | 0.015 | ns |
| | IR64-7a | 0.433 | 0.009 | ns | 0.487 | 0.012 | ns | 0.363 | 0.013 | ns | 0.293 | 0.014 | ns |
| | IR64-9a | 0.48 | 0.009 | ns | 0.52 | 0.005 | ns | 0.398 | 0.016 | ns | 0.316 | 0.012 | ns |
| | IR64-134a | 0.487 | 0.007 | ns | 0.532 | 0.008 | ns | 0.396 | 0.004 | ns | 0.315 | 0.003 | ns |
| | IR64-134b | 0.461 | 0.011 | ns | 0.515 | 0.013 | ns | 0.371 | 0.022 | ns | 0.299 | 0.02 | ns |
| | IR64-136a | 0.459 | 0.011 | ns | 0.508 | 0.012 | ns | 0.378 | 0.012 | ns | 0.302 | 0.007 | ns |
| | IR64-7b | 0.445 | 0.011 | ns | 0.5 | 0.014 | ns | 0.368 | 0.012 | ns | 0.289 | 0.01 | ns |
| | IR64-134d | 0.471 | 0.005 | ns | 0.522 | 0.008 | ns | 0.388 | 0.013 | ns | 0.303 | 0.011 | ns |
| NDVI | Ciherang-Sub1 WT | 0.852 | 0.011 | | 0.885 | 0.011 | | 0.871 | 0.012 | | 0.83 | 0.015 | |
| | CS-1h | 0.869 | 0.015 | ns | 0.896 | 0.011 | ns | 0.877 | 0.008 | ns | 0.828 | 0.01 | ns |
| | CS-3b | 0.835 | 0.008 | ns | 0.862 | 0.006 | ns | 0.857 | 0.008 | ns | 0.8 | 0.003 | ns |
| | CS-4 | 0.844 | 0.022 | ns | 0.875 | 0.005 | ns | 0.873 | 0.006 | ns | 0.829 | 0.008 | ns |
| | CS-1i | 0.865 | 0.01 | ns | 0.886 | 0.006 | ns | 0.877 | 0.008 | ns | 0.833 | 0.014 | ns |
| | CS-6d | 0.863 | 0.01 | ns | 0.88 | 0.006 | ns | 0.873 | 0.008 | ns | 0.829 | 0.011 | ns |
| | IR 64 WT | 0.863 | 0.011 | | 0.891 | 0.01 | | 0.859 | 0.003 | | 0.809 | 0.003 | |
| | IR64-5d | 0.881 | 0.008 | ns | 0.906 | 0.008 | ns | 0.86 | 0.005 | ns | 0.816 | 0.009 | ns |
| | IR64-7a | 0.887 | 0.008 | ns | 0.911 | 0.007 | ns | 0.866 | 0.009 | ns | 0.82 | 0.01 | ns |
| | IR64-9a | 0.859 | 0.009 | ns | 0.886 | 0.006 | ns | 0.856 | 0.008 | ns | 0.809 | 0.01 | ns |
| | IR64-134a | 0.887 | 0.009 | ns | 0.901 | 0.006 | ns | 0.86 | 0.004 | ns | 0.814 | 0.007 | ns |
| | IR64-134b | 0.886 | 0.013 | ns | 0.902 | 0.008 | ns | 0.862 | 0.001 | ns | 0.812 | 0.002 | ns |
| | IR64-136a | 0.875 | 0.009 | ns | 0.901 | 0.007 | ns | 0.856 | 0.007 | ns | 0.807 | 0.01 | ns |
| | IR64-7b | 0.869 | 0.013 | ns | 0.894 | 0.009 | ns | 0.858 | 0.006 | ns | 0.807 | 0.005 | ns |
| | IR64-134d | 0.863 | 0.012 | ns | 0.89 | 0.01 | ns | 0.851 | 0.004 | ns | 0.802 | 0.006 | ns |

DAP: days after planting; SE: standard error; *: significant at $P<0.05$; **: significant at $P<0.01$; ns = not significant ($P>0.05$)
