## Supplementary material for "From lab to field: analyses of genome-edited bacterial blight resistant rice": Table S7

Table S7: NDRE and NDVI measurements of EPF in Colombia during wet season of 2021

| Spectra | Genotype | 37DAP | SE | P-value | 52DAP | SE | P-value | 76DAP | SE | P-value | 82DAP | SE | P-value | 85DAP | SE | P-value | 89DAP | SE | P-value |
|---|---|---|---|---|---|---|---|---|---|---|---|---|---|---|---|---|---|---|---|
| NDRE | Ciherang-Sub1 WT | 0.009 | 0.006 | | 0.212 | 0.016 | | 0.493 | 0.007 | | 0.411 | 0.006 | | 0.402 | 0.008 | | 0.353 | 0.008 | |
| | CS-1h | 0.001 | 0.006 | ns | 0.224 | 0.02 | ns | 0.495 | 0.017 | ns | 0.433 | 0.02 | ns | 0.419 | 0.014 | ns | 0.38 | 0.013 | ns |
| | CS-3b | -0.015 | 0.007 | * | 0.223 | 0.006 | ns | 0.454 | 0.011 | ns | 0.384 | 0.007 | ns | 0.376 | 0.005 | ns | 0.327 | 0.002 | ns |
| | CS-4 | -0.016 | 0.007 | * | 0.2 | 0.014 | ns | 0.462 | 0.025 | ns | 0.389 | 0.023 | ns | 0.386 | 0.018 | ns | 0.344 | 0.017 | ns |
| | CS-1i | 0.008 | 0.005 | ns | 0.205 | 0.005 | ns | 0.495 | 0.008 | ns | 0.423 | 0.008 | ns | 0.413 | 0.006 | ns | 0.368 | 0.009 | ns |
| | CS-6d | -0.013 | 0.007 | ns | 0.224 | 0.019 | ns | 0.449 | 0.023 | ns | 0.38 | 0.02 | ns | 0.374 | 0.019 | ns | 0.332 | 0.016 | ns |
| | IR 64 WT | -0.022 | 0.005 | | 0.213 | 0.011 | | 0.432 | 0.025 | | 0.347 | 0.028 | | 0.335 | 0.025 | | 0.304 | 0.026 | |
| | IR64-5d | 0.013 | 0.005 | ** | 0.244 | 0.005 | ns | 0.455 | 0.007 | ns | 0.352 | 0.006 | ns | 0.331 | 0.007 | ns | 0.281 | 0.007 | ns |
| | IR64-7a | -0.011 | 0.007 | ns | 0.243 | 0.014 | ns | 0.429 | 0.024 | ns | 0.341 | 0.021 | ns | 0.329 | 0.018 | ns | 0.285 | 0.016 | ns |
| | IR64-9a | 0.005 | 0.005 | * | 0.232 | 0.021 | ns | 0.436 | 0.026 | ns | 0.335 | 0.023 | ns | 0.316 | 0.02 | ns | 0.267 | 0.012 | ns |
| | IR64-134a | -0.008 | 0.002 | ns | 0.234 | 0.016 | ns | 0.418 | 0.02 | ns | 0.322 | 0.018 | ns | 0.307 | 0.015 | ns | 0.264 | 0.01 | ns |
| | IR64-134b | -0.014 | 0.006 | ns | 0.234 | 0.005 | ns | 0.431 | 0.023 | ns | 0.338 | 0.02 | ns | 0.328 | 0.019 | ns | 0.298 | 0.016 | ns |
| | IR64-136a | 0.002 | 0.002 | * | 0.236 | 0.015 | ns | 0.459 | 0.01 | ns | 0.362 | 0.009 | ns | 0.344 | 0.012 | ns | 0.292 | 0.01 | ns |
| | IR64-7b | -0.006 | 0.005 | ns | 0.227 | 0.011 | ns | 0.444 | 0.023 | ns | 0.351 | 0.022 | ns | 0.334 | 0.023 | ns | 0.291 | 0.02 | ns |
| | IR64-134d | -0.014 | 0.006 | ns | 0.226 | 0.008 | ns | 0.433 | 0.012 | ns | 0.353 | 0.013 | ns | 0.342 | 0.012 | ns | 0.3 | 0.01 | ns |
| NDVI | Ciherang-Sub1 WT | 0.377 | 0.014 | | 0.704 | 0.025 | | 0.879 | 0.006 | | 0.819 | 0.005 | | 0.823 | 0.007 | | 0.813 | 0.008 | |
| | CS-1h | 0.357 | 0.012 | ns | 0.72 | 0.029 | ns | 0.878 | 0.007 | ns | 0.828 | 0.011 | ns | 0.827 | 0.006 | ns | 0.829 | 0.006 | ns |
| | CS-3b | 0.327 | 0.018 | ns | 0.721 | 0.009 | ns | 0.867 | 0.005 | ns | 0.811 | 0.007 | ns | 0.817 | 0.004 | ns | 0.804 | 0.003 | ns |
| | CS-4 | 0.332 | 0.015 | ns | 0.697 | 0.023 | ns | 0.867 | 0.011 | ns | 0.804 | 0.016 | ns | 0.812 | 0.009 | ns | 0.801 | 0.01 | ns |
| | CS-1i | 0.367 | 0.017 | ns | 0.688 | 0.009 | ns | 0.873 | 0.004 | ns | 0.813 | 0.004 | ns | 0.826 | 0.001 | ns | 0.816 | 0.006 | ns |
| | CS-6d | 0.341 | 0.015 | ns | 0.72 | 0.031 | ns | 0.86 | 0.012 | ns | 0.804 | 0.014 | ns | 0.807 | 0.012 | ns | 0.798 | 0.007 | ns |
| | IR 64 WT | 0.345 | 0.012 | ns | 0.733 | 0.015 | ns | 0.864 | 0.013 | ns | 0.801 | 0.018 | ns | 0.794 | 0.017 | ns | 0.793 | 0.012 | ns |
| | IR64-5d | 0.417 | 0.007 | ns | 0.77 | 0.006 | ns | 0.868 | 0.004 | ns | 0.803 | 0.003 | ns | 0.797 | 0.003 | ns | 0.785 | 0.005 | ns |
| | IR64-7a | 0.36 | 0.019 | ns | 0.738 | 0.045 | ns | 0.859 | 0.013 | ns | 0.799 | 0.016 | ns | 0.791 | 0.016 | ns | 0.788 | 0.01 | ns |
| | IR64-9a | 0.409 | 0.02 | * | 0.756 | 0.028 | ns | 0.861 | 0.013 | ns | 0.797 | 0.017 | ns | 0.791 | 0.015 | ns | 0.78 | 0.01 | ns |
| | IR64-134a | 0.358 | 0.009 | ns | 0.754 | 0.017 | ns | 0.853 | 0.011 | ns | 0.787 | 0.013 | ns | 0.784 | 0.012 | ns | 0.775 | 0.01 | ns |
| | IR64-134b | 0.353 | 0.025 | ns | 0.756 | 0.005 | ns | 0.863 | 0.011 | ns | 0.802 | 0.014 | ns | 0.798 | 0.012 | ns | 0.788 | 0.01 | ns |
| | IR64-136a | 0.388 | 0.017 | ns | 0.742 | 0.018 | ns | 0.868 | 0.005 | ns | 0.808 | 0.006 | ns | 0.803 | 0.009 | ns | 0.791 | 0.005 | ns |
| | IR64-7b | 0.358 | 0.009 | ns | 0.746 | 0.014 | ns | 0.864 | 0.011 | ns | 0.803 | 0.013 | ns | 0.795 | 0.015 | ns | 0.788 | 0.01 | ns |
| | IR64-134d | 0.35 | 0.017 | | 0.73 | 0.021 | | 0.87 | 0.008 | | 0.808 | 0.01 | | 0.804 | 0.009 | | 0.797 | 0.006 | |

DAP: days after planting; SE: standard error; *: significant at $P<0.05$; **: significant at $P<0.01$; ns = not significant ($P>0.05$)
