## Supplementary material for "From lab to field: analyses of genome-edited bacterial blight resistant rice": Table S8

Table S8: Quantification of agronomic traits for GE lines from EFPs in Colombia and the Philippines

| Season | Genotype | 100% Flowering (DAP) | SE | P-Value | 1000 Grain Weight (g) | SE | P-Value | 100 Grain Weight (g) | SE | P-Value | Bulk Grain weight (g) | SE | P-Value | Dry Biomass (g) | SE | P-Value | Grain Yield (kg/m2) | SE | P-Value | Harvest Index | SE | P-Value | Panicle Length (cm) | SE | P-Value | Panicle Number | SE | P-Value | Plant height (cm) | SE | P-Value | Single Plant grain Yield (g) | SE | P-Value | Yield (Ton/Ha) | SE | P-Value |
|---|---|---|---|---|---|---|---|---|---|---|---|---|---|---|---|---|---|---|---|---|---|---|---|---|---|---|---|---|---|---|---|---|---|---|---|---|---|
| Dry 2020 (Col) | Ciherang-Sub1 WT | 74 | 1.22 | | 26.05 | 0.2 | | (n.d.) | | | 2355.61 | 73.42 | | 27.61 | 1.07 | | 1.14 | 0 | | 0.62 | 0.003 | | 24.34 | 0.21 | | 11.43 | 0.39 | | 92.75 | 0.71 | | 45.58 | 1.7 | | 8.2 | 0.26 | |
| | CS-1h | 73.25 | 1.03 | ns | 26.83 | 0.4 | ns | (n.d.) | | | 2177.6 | 104.52 | ns | 31.18 | 1.23 | * | 1.13 | 0 | ns | 0.59 | 0.007 | ** | 24.99 | 0.24 | ns | 10.75 | 0.3 | ns | 98 | 0.84 | ** | 45.16 | 1.7 | ns | 7.72 | 0.37 | ns |
| | CS-3b | 71.25 | 0.25 | ns | 27.25 | 0.2 | * | (n.d.) | | | 2252.32 | 26.31 | ns | 26.21 | 0.73 | ns | 1.07 | 0 | ns | 0.62 | 0.002 | ns | 23.42 | 0.23 | * | 11 | 0.33 | ns | 92.86 | 0.52 | ns | 42.97 | 1.2 | ns | 7.85 | 0.09 | ns |
| | CS-4 | 73.25 | 1.03 | ns | 26.59 | 0.2 | ns | (n.d.) | | | 2187.94 | 128.41 | ns | 25.51 | 0.94 | ns | 1.03 | 0 | ns | 0.62 | 0.003 | ns | 23.91 | 0.17 | ns | 11.64 | 0.36 | ns | 91.46 | 0.76 | ns | 41.17 | 1.5 | ns | 7.61 | 0.45 | ns |
| | CS-1b | 75.25 | 1.18 | ns | 27.47 | 0.2 | ** | (n.d.) | | | 2347.33 | 107.12 | ns | 31.41 | 1.38 | * | 1.12 | 0 | ns | 0.59 | 0.01 | ** | 25.08 | 0.24 | ns | 10.68 | 0.46 | ns | 95.07 | 0.84 | ns | 44.86 | 1.7 | ns | 8.25 | 0.38 | ns |
| | CS-6d | 72 | 1 | ns | 26.4 | 0.2 | ns | (n.d.) | | | 2146.73 | 93.12 | ns | 23.4 | 0.55 | * | 0.96 | 0 | * | 0.62 | 0.004 | ns | 23.23 | 0.17 | ** | 10.14 | 0.26 | ns | 92.54 | 0.57 | ns | 38.49 | 0.8 | * | 7.48 | 0.32 | ns |
| | IR64 WT | 68.25 | 0.75 | | 28.28 | 0.1 | | (n.d.) | | | 2332.83 | 111.46 | | 29.06 | 1.25 | | 1.09 | 0 | | 0.6 | 0.004 | | 26.26 | 0.21 | | 14.21 | 0.58 | | 91.71 | 0.62 | | 43.74 | 1.6 | | 8.16 | 0.39 | |
| | IR64-5d | 65.5 | 0.29 | ns | 27.82 | 0.1 | ns | (n.d.) | | | 2356.84 | 45.83 | ns | 29.03 | 0.91 | ns | 1.13 | 0 | ns | 0.61 | 0.004 | ns | 25.99 | 0.2 | ns | 13.75 | 0.42 | ns | 91.75 | 0.63 | ns | 45.39 | 1.3 | ns | 8.25 | 0.16 | ns |
| | IR64-7a | 68.25 | 0.75 | ns | 27.39 | 0.5 | ns | (n.d.) | | | 2645.54 | 65.07 | * | 29.11 | 1.14 | ns | 1.21 | 0.1 | ns | 0.62 | 0.005 | ns | 25.31 | 0.18 | * | 14.64 | 0.68 | ns | 93.82 | 0.56 | ns | 48.41 | 1.8 | ns | 9.25 | 0.23 | * |
| | IR64-9a | 67.5 | 0.87 | ns | 28.4 | 0.2 | ns | (n.d.) | | | 2577.87 | 55.83 | ns | 31.73 | 1.18 | ns | 1.26 | 0.1 | * | 0.61 | 0.003 | ns | 26.05 | 0.25 | ns | 16.64 | 0.86 | ** | 94.32 | 0.49 | ns | 50.27 | 1.9 | 0.05 | 8.97 | 0.19 | ns |
| | IR64-134a | 68.25 | 0.75 | ns | 27.46 | 0.5 | ns | (n.d.) | | | 2441.92 | 94.52 | ns | 30.89 | 1.27 | ns | 1.15 | 0.1 | ns | 0.6 | 0.006 | ns | 25.51 | 0.18 | ns | 14.21 | 0.52 | ns | 93.11 | 0.78 | ns | 45.88 | 2.2 | ns | 8.56 | 0.33 | ns |
| | IR64-134b | 66.5 | 0.87 | ns | 26.94 | 0.3 | ** | (n.d.) | | | 2087.31 | 39.65 | ns | 26.38 | 0.85 | ns | 0.91 | 0 | * | 0.58 | 0.011 | * | 25.17 | 0.21 | ** | 12.32 | 0.47 | ns | 87.46 | 0.6 | ** | 36.58 | 1.5 | * | 7.3 | 0.14 | ns |
| | IR64-136a | 65.5 | 0.29 | ns | 26.13 | 0.2 | ** | (n.d.) | | | 2350.57 | 118.7 | ns | 28.52 | 1.28 | ns | 1.07 | 0.1 | ns | 0.6 | 0.005 | ns | 24.84 | 0.27 | ** | 15.79 | 0.98 | ns | 89.43 | 0.6 | ns | 42.93 | 1.9 | ns | 8.22 | 0.42 | ns |
| | IR64-7b | 66.5 | 0.87 | ns | 27.92 | 0.1 | ns | (n.d.) | | | 2443.75 | 139.82 | ns | 27.65 | 0.98 | ns | 1.18 | 0 | ns | 0.63 | 0.004 | * | 26.11 | 0.22 | ns | 13.57 | 0.61 | ns | 90.68 | 1.3 | ns | 47.03 | 1.8 | ns | 8.62 | 0.49 | ns |
| | IR64-134c | 69 | 0 | ns | 26.16 | 0.1 | ** | (n.d.) | | | 2578.72 | 66.74 | ns | 30.79 | 1.3 | ns | 1.14 | 0.1 | ns | 0.6 | 0.004 | ns | 25.06 | 0.21 | ** | 13.86 | 0.67 | ns | 92.54 | 0.59 | ns | 45.67 | 1.8 | ns | 9.05 | 0.23 | ns |
| Wet 2020 (Phil) | IR64 WT | 67.75 | | | (n.d.) | | | 2.738 | | | (n.d.) | | | 30.323 | | | (n.d.) | | | (n.d.) | | | 27.073 | | | 20.361 | | | 98.249 | | | 39.95 | | | 6.994 | | |
| | IR64-5d | 64.25 | | | (n.d.) | | | 2.661 | | | (n.d.) | | | 31.158 | | | (n.d.) | | | (n.d.) | | | 25.88 | | | 18.588 | | | 91.778 | | | 36.837 | | | 6.729 | | |
| | IR64-7a | 66.5 | | | (n.d.) | | | 2.525 | | | (n.d.) | | | 33.323 | | | (n.d.) | | | (n.d.) | | | 26.457 | | | 18.768 | | | 99.922 | | | 40.367 | | | 7.413 | | |
| | IR64-7b | 65.25 | | | (n.d.) | | | 2.629 | | | (n.d.) | | | 30.966 | | | (n.d.) | | | (n.d.) | | | 26.396 | | | 19.745 | | | 94.061 | | | 46.01 | | | 8.305 | | |
| | IR64-9a | 64.75 | | | (n.d.) | | | 2.678 | | | (n.d.) | | | 32.467 | | | (n.d.) | | | (n.d.) | | | 26.563 | | | 20.732 | | | 97.026 | | | 41.085 | | | 6.956 | | |
| | IR64-134b | 65.75 | | | (n.d.) | | | 2.543 | | | (n.d.) | | | 33.227 | | | (n.d.) | | | (n.d.) | | | 26.286 | | | 21.037 | | | 93.565 | | | 41.892 | | | 7.434 | | |
| | IR64-134a | 65.5 | | | (n.d.) | | | 2.514 | | | (n.d.) | | | 34.816 | | | (n.d.) | | | (n.d.) | | | 26.242 | | | 19.672 | | | 96.015 | | | 40.987 | | | 7.052 | | |
| | IR64-134d | 65.25 | | | (n.d.) | | | 2.541 | | | (n.d.) | | | 29.76 | | | (n.d.) | | | (n.d.) | | | 26.869 | | | 19.503 | | | 96.253 | | | 42.107 | | | 7.3 | | |
| | IR64-136a | 65.25 | | | (n.d.) | | | 2.51 | | | (n.d.) | | | 26.336 | | | (n.d.) | | | (n.d.) | | | 26.56 | | | 20.794 | | | 93.603 | | | 43.651 | | | 6.952 | | |
| | Ciherang-Sub1 WT | 65.059 | | | (n.d.) | | | 2.683 | | | (n.d.) | | | 35.149 | | | (n.d.) | | | (n.d.) | | | 25.383 | | | 16.65 | | | 98.921 | | | 44.413 | | | 7.783 | | |
| | CS-1i | 66.386 | | | (n.d.) | | | 2.804 | | | (n.d.) | | | 33.771 | | | (n.d.) | | | (n.d.) | | | 27.155 | | | 16.2 | | | 105.945 | | | 49.367 | | | 7.751 | | |
| | CS-1h | 65.649 | | | (n.d.) | | | 2.745 | | | (n.d.) | | | 40.704 | | | (n.d.) | | | (n.d.) | | | 26.498 | | | 16.6 | | | 104.139 | | | 48.352 | | | 7.642 | | |
| | CS-3b | 64.941 | | | (n.d.) | | | 2.687 | | | (n.d.) | | | 32.801 | | | (n.d.) | | | (n.d.) | | | 25.555 | | | 17.1 | | | 98.796 | | | 45.511 | | | 7.623 | | |
| | CS-6d | 63.472 | | | (n.d.) | | | 2.67 | | | (n.d.) | | | 32.298 | | | (n.d.) | | | (n.d.) | | | 25.338 | | | 15.275 | | | 99.214 | | | 43.102 | | | 8.038 | | |
| Wet 2021 (Col) | Ciherang-Sub1 WT | 76.75 | 1.109 | | 26.25 | 0.2 | | (n.d.) | | | 1760.00 | 78.60 | | 24.28 | 1.16 | | 0.9 | 0.05 | | 0.6 | 0.03 | | 23.16 | 0.37 | | 10.04 | 0.78 | | 76.11 | 1.06 | | 36.20 | 1.94 | | 4.40 | 0.20 | |
| | CS-1h | 80.00 | 0.408 | * | 28.02 | 0.2 | ** | (n.d.) | | | 1631.25 | 131.47 | ns | 28.85 | 2.94 | * | 0.94 | 0.09 | ns | 0.57 | 0.05 | * | 24.47 | 0.31 | * | 9.93 | 0.59 | ns | 81.79 | 1.10 | * | 37.60 | 3.56 | ns | 4.08 | 0.33 | ns |
| | CS-3b | 77.00 | 0.577 | ns | 27.57 | 0.2 | ** | (n.d.) | | | 1664.25 | 74.28 | ns | 27.00 | 1.05 | ns | 0.99 | 0.05 | ns | 0.59 | 0.03 | ns | 23.63 | 0.33 | ns | 10.36 | 0.22 | ns | 77.86 | 1.17 | ns | 39.56 | 2.01 | ns | 4.16 | 0.19 | ns |
| | CS-4 | 77.75 | 1.181 | ns | 27.06 | 0.1 | ** | (n.d.) | | | 1500.25 | 56.26 | * | 25.00 | 1.48 | ns | 0.9 | 0.06 | ns | 0.59 | 0.04 | ns | 23.29 | 0.25 | ns | 10.21 | 0.43 | ns | 74.04 | 0.68 | ns | 35.96 | 2.37 | ns | 3.75 | 0.14 | * |
| | CS-1b | 79.00 | 1.08 | ns | 28.24 | 0.2 | ** | (n.d.) | | | 1895.25 | 55.79 | ns | 24.56 | 0.76 | ns | 0.88 | 0.02 | ns | 0.59 | 0.01 | ns | 24.63 | 0.20 | ** | 9.39 | 0.31 | ns | 75.00 | 2.67 | ns | 35.31 | 0.69 | ns | 4.74 | 0.14 | ns |
| | CS-6d | 78.00 | 0.707 | ns | 26.33 | 0.1 | ns | (n.d.) | | | 1486.75 | 62.79 | * | 24.25 | 2.06 | ns | 0.92 | 0.09 | ns | 0.6 | 0.06 | ns | 22.82 | 0.37 | ns | 9.50 | 0.60 | ns | 77.82 | 1.01 | ns | 36.61 | 3.52 | ns | 3.72 | 0.16 | * |
| | IR64 WT | 74.75 | 1.377 | | 28.07 | 0.2 | | (n.d.) | | | 1403.00 | 70.30 | | 30.35 | 2.99 | | 0.74 | 0.01 | | 0.49 | 0.01 | | 24.41 | 0.27 | | 12.61 | 0.57 | | 81.61 | 1.30 | | 33.21 | 0.38 | | 3.51 | 0.16 | |
| | IR64-5d | 71.00 | 1 | * | 27.71 | 0.1 | ns | (n.d.) | | | 1517.50 | 48.09 | ns | 32.53 | 1.59 | ns | 0.74 | 0.04 | ns | 0.48 | 0.02 | ns | 24.91 | 0.19 | ns | 11.86 | 0.12 | ns | 81.54 | 0.88 | ns | 29.46 | 1.51 | ns | 3.79 | 0.19 | ns |
| | IR64-7a | 72.00 | 0.816 | ns | 26.25 | 0.2 | ** | (n.d.) | | | 1475.00 | 173.81 | ns | 28.77 | 2.49 | ns | 0.81 | 0.05 | ns | 0.53 | 0.03 | ns | 24.63 | 0.30 | ns | 11.43 | 0.89 | ns | 82.71 | 0.95 | ns | 29.56 | 1.75 | ns | 3.69 | 0.12 | ns |
| | IR64-9a | 70.25 | 0.25 | * | 28.41 | 0.2 | ns | (n.d.) | | | 1440.25 | 110.78 | ns | 30.79 | 3.31 | ns | 0.74 | 0.08 | ns | 0.49 | 0.05 | ns | 25.05 | 0.55 | ns | 12.86 | 0.81 | ns | 82.00 | 0.95 | ns | 32.35 | 3.35 | ns | 3.60 | 0.42 | ns |
| | IR64-134a | 71.25 | 0.75 | * | 25.70 | 0.2 | ** | (n.d.) | | | 1396.25 | 71.57 | ns | 32.00 | 4.73 | ns | 0.71 | 0.04 | ns | 0.48 | 0.03 | ns | 24.41 | 0.37 | ns | 11.14 | 0.74 | ns | 79.64 | 0.94 | ns | 29.44 | 1.79 | ns | 3.49 | 0.27 | ns |
| | IR64-134b | 73.25 | 0.75 | ns | 25.56 | 0.1 | ** | (n.d.) | | | 1458.50 | 151.29 | ns | 28.55 | 5.18 | ns | 0.84 | 0.07 | ns | 0.54 | 0.05 | ns | 23.98 | 0.33 | ns | 12.75 | 1.15 | ns | 76.71 | 1.14 | ns | 28.32 | 2.41 | ns | 3.65 | 0.19 | ns |
| | IR64-136a | 71.75 | 0.854 | * | 24.74 | 0.1 | ** | (n.d.) | | | 1485.00 | 64.85 | ns | 35.24 | 3.24 | ns | 0.89 | 0.13 | ns | 0.5 | 0.08 | ns | 24.10 | 0.33 | ns | 14.00 | 0.64 | ns | 75.89 | 1.06 | ** | 33.62 | 5.06 | ns | 3.71 | 0.38 | ns |
| | IR64-7b | 79.50 | 8.51 | ns | 27.45 | 0.1 | * | (n.d.) | | | 1561.50 | 125.18 | ns | 32.78 | 4.10 | ns | 0.98 | 0.10 | * | 0.55 | 0.06 | ns | 24.34 | 0.38 | ns | 12.68 | 0.69 | ns | 76.79 | 1.08 | * | 39.07 | 4.11 | * | 3.90 | 0.17 | ns |
| | IR64-134c | 75.50 | 0.5 | ns | 25.51 | 0.1 | ** | (n.d.) | | | 1658.75 | 77.65 | ns | 30.78 | 1.01 | ns | 0.83 | 0.04 | ns | 0.52 | 0.02 | ns | 24.31 | 0.09 | ns | 12.18 | 0.24 | ns | 82.43 | 0.85 | ns | 35.49 | 1.70 | ns | 4.15 | 0.33 | ns |

SE: standard error; *: significant at P<0.05; **: significant at P<0.01;  ns = not significant (P>0.05)

N.A: Not applicable; n.d.: not determined
