## Supplementary material for "From lab to field: analyses of genome-edited bacterial blight resistant rice": Table S9

Table S9: NDRE and NDVI measurements of EPF in Colombia during wet season of 2022-2023

| | Spectra | Genotype | 39DAP | SE | P-value | 47DAP | SE | P-value | 53DAP | SE | P-value | 64DAP | SE | P-value | 77DAP | SE | P-value | 83DAP | SE | P-value | 89DAP | SE | P-value |
|---|---|---|---|---|---|---|---|---|---|---|---|---|---|---|---|---|---|---|---|---|---|---|---|
| 0% N | NDRE | Ciherang-Sub1 WT | 0.301 | 0.011 | | 0.383 | 0.008 | | 0.387 | 0.007 | | 0.365 | 0.015 | | 0.214 | 0.027 | | 0.400 | 0.021 | | 0.320 | 0.006 | |
| | | CS-3b | 0.296 | 0.012 | ns | 0.379 | 0.009 | ns | 0.388 | 0.014 | ns | 0.358 | 0.020 | ns | 0.213 | 0.053 | ns | 0.393 | 0.008 | ns | 0.312 | 0.005 | ns |
| | | CS-1b | 0.303 | 0.010 | ns | 0.393 | 0.012 | ns | 0.396 | 0.016 | ns | 0.380 | 0.015 | ns | 0.252 | 0.040 | ns | 0.408 | 0.003 | ns | 0.327 | 0.006 | ns |
| | | CS-6d | 0.286 | 0.004 | ns | 0.379 | 0.003 | ns | 0.379 | 0.003 | ns | 0.353 | 0.009 | ns | 0.227 | 0.026 | ns | 0.384 | 0.016 | ns | 0.312 | 0.005 | ns |
| | | IR64 WT | 0.263 | 0.010 | | 0.345 | 0.004 | | 0.365 | 0.008 | | 0.357 | 0.015 | | 0.206 | 0.043 | | 0.361 | 0.005 | | 0.290 | 0.008 | |
| | | IR64-5d | 0.266 | 0.010 | ns | 0.343 | 0.010 | ns | 0.354 | 0.019 | ns | 0.344 | 0.038 | ns | 0.138 | 0.093 | ns | 0.340 | 0.029 | ns | 0.270 | 0.029 | ns |
| | | IR64-7a | 0.263 | 0.003 | ns | 0.343 | 0.004 | ns | 0.351 | 0.008 | ns | 0.321 | 0.014 | ns | 0.127 | 0.060 | ns | 0.333 | 0.016 | ns | 0.267 | 0.010 | ns |
| | | IR64-9a | 0.269 | 0.014 | ns | 0.339 | 0.012 | ns | 0.353 | 0.021 | ns | 0.318 | 0.020 | ns | 0.060 | 0.016 | ns | 0.321 | 0.006 | ns | 0.249 | 0.008 | ns |
| | | IR64-134a | 0.262 | 0.010 | ns | 0.339 | 0.007 | ns | 0.359 | 0.013 | ns | 0.330 | 0.013 | ns | 0.125 | 0.039 | ns | 0.346 | 0.008 | ns | 0.270 | 0.008 | ns |
| | | IR64-134b | 0.275 | 0.010 | ns | 0.356 | 0.009 | ns | 0.379 | 0.012 | ns | 0.349 | 0.009 | ns | 0.114 | 0.053 | ns | 0.339 | 0.011 | ns | 0.275 | 0.005 | ns |
| | | IR64-136a | 0.277 | 0.007 | ns | 0.356 | 0.005 | ns | 0.370 | 0.014 | ns | 0.351 | 0.020 | ns | 0.158 | 0.055 | ns | 0.352 | 0.012 | ns | 0.277 | 0.012 | ns |
| | | IR64-7b | 0.264 | 0.007 | ns | 0.341 | 0.006 | ns | 0.353 | 0.009 | ns | 0.315 | 0.009 | ns | 0.108 | 0.058 | ns | 0.325 | 0.010 | ns | 0.262 | 0.004 | ns |
| | | IR64-134d | 0.266 | 0.006 | ns | 0.344 | 0.006 | ns | 0.361 | 0.008 | ns | 0.341 | 0.005 | ns | 0.087 | 0.007 | ns | 0.344 | 0.010 | ns | 0.267 | 0.007 | ns |
| | NDVI | Ciherang-Sub1 WT | 0.787 | 0.007 | | 0.845 | 0.004 | | 0.892 | 0.002 | | 0.839 | 0.006 | | 0.688 | 0.006 | | 0.799 | 0.012 | | 0.759 | 0.011 | |
| | | CS-3b | 0.784 | 0.005 | ns | 0.846 | 0.003 | ns | 0.898 | 0.007 | ns | 0.841 | 0.006 | ns | 0.692 | 0.011 | ns | 0.803 | 0.010 | ns | 0.764 | 0.012 | ns |
| | | CS-1b | 0.782 | 0.004 | ns | 0.844 | 0.004 | ns | 0.892 | 0.006 | ns | 0.843 | 0.005 | ns | 0.688 | 0.012 | ns | 0.796 | 0.009 | ns | 0.760 | 0.007 | ns |
| | | CS-6d | 0.776 | 0.001 | ns | 0.839 | 0.002 | ns | 0.886 | 0.001 | ns | 0.831 | 0.004 | ns | 0.677 | 0.002 | ns | 0.791 | 0.007 | ns | 0.757 | 0.011 | ns |
| | | IR64 WT | 0.770 | 0.002 | | 0.843 | 0.001 | | 0.886 | 0.003 | | 0.840 | 0.001 | | 0.689 | 0.009 | | 0.798 | 0.004 | | 0.738 | 0.003 | |
| | | IR64-5d | 0.766 | 0.002 | ns | 0.836 | 0.005 | ns | 0.883 | 0.005 | ns | 0.831 | 0.011 | ns | 0.665 | 0.029 | ns | 0.781 | 0.016 | ns | 0.727 | 0.013 | ns |
| | | IR64-7a | 0.768 | 0.002 | ns | 0.837 | 0.001 | ns | 0.884 | 0.001 | ns | 0.830 | 0.006 | ns | 0.659 | 0.013 | ns | 0.782 | 0.010 | ns | 0.727 | 0.008 | ns |
| | | IR64-9a | 0.769 | 0.004 | ns | 0.838 | 0.005 | ns | 0.887 | 0.007 | ns | 0.827 | 0.007 | ns | 0.660 | 0.013 | ns | 0.780 | 0.009 | ns | 0.726 | 0.006 | ns |
| | | IR64-134a | 0.765 | 0.005 | ns | 0.840 | 0.003 | ns | 0.888 | 0.005 | ns | 0.833 | 0.004 | ns | 0.671 | 0.011 | ns | 0.794 | 0.006 | ns | 0.735 | 0.003 | ns |
| | | IR64-134b | 0.770 | 0.002 | ns | 0.840 | 0.003 | ns | 0.888 | 0.004 | ns | 0.824 | 0.003 | ns | 0.658 | 0.010 | ns | 0.780 | 0.008 | ns | 0.732 | 0.006 | ns |
| | | IR64-136a | 0.772 | 0.005 | ns | 0.840 | 0.005 | ns | 0.886 | 0.006 | ns | 0.833 | 0.004 | ns | 0.670 | 0.016 | ns | 0.788 | 0.008 | ns | 0.732 | 0.006 | ns |
| | | IR64-7b | 0.764 | 0.003 | ns | 0.836 | 0.001 | ns | 0.884 | 0.003 | ns | 0.826 | 0.002 | ns | 0.658 | 0.006 | ns | 0.784 | 0.008 | ns | 0.724 | 0.003 | ns |
| | | IR64-134d | 0.769 | 0.004 | ns | 0.843 | 0.003 | ns | 0.889 | 0.004 | ns | 0.834 | 0.001 | ns | 0.678 | 0.002 | ns | 0.794 | 0.003 | ns | 0.736 | 0.005 | ns |

| | Spectra | Genotype | 39DAP | SE | P-value | 47DAP | SE | P-value | 53DAP | SE | P-value | 64DAP | SE | P-value | 77DAP | SE | P-value | 83DAP | SE | P-value | 89DAP | SE | P-value |
|---|---|---|---|---|---|---|---|---|---|---|---|---|---|---|---|---|---|---|---|---|---|---|---|
| 50% N | NDRE | Ciherang-Sub1 WT | 0.384 | 0.013 | | 0.467 | 0.016 | | 0.482 | 0.017 | | 0.457 | 0.016 | | 0.252 | 0.023 | | 0.432 | 0.004 | | 0.332 | 0.004 | |
| | | CS-3b | 0.380 | 0.006 | ns | 0.463 | 0.007 | ns | 0.478 | 0.007 | ns | 0.447 | 0.005 | ns | 0.248 | 0.029 | ns | 0.418 | 0.007 | ns | 0.328 | 0.005 | ns |
| | | CS-1b | 0.378 | 0.015 | ns | 0.466 | 0.017 | ns | 0.483 | 0.021 | ns | 0.463 | 0.020 | ns | 0.260 | 0.021 | ns | 0.441 | 0.018 | ns | 0.342 | 0.019 | ns |
| | | CS-6d | 0.360 | 0.017 | ns | 0.441 | 0.020 | ns | 0.461 | 0.023 | ns | 0.433 | 0.018 | ns | 0.221 | 0.024 | ns | 0.421 | 0.007 | ns | 0.326 | 0.011 | ns |
| | | IR64 WT | 0.336 | 0.008 | | 0.416 | 0.011 | | 0.424 | 0.015 | | 0.393 | 0.015 | | 0.107 | 0.007 | | 0.354 | 0.002 | | 0.274 | 0.007 | |
| | | IR64-5d | 0.369 | 0.005 | ns | 0.451 | 0.004 | ns | 0.471 | 0.003 | ns | 0.428 | 0.003 | ns | 0.134 | 0.009 | ns | 0.364 | 0.007 | ns | 0.282 | 0.001 | ns |
| | | IR64-7a | 0.344 | 0.014 | ns | 0.444 | 0.011 | ns | 0.461 | 0.015 | ns | 0.430 | 0.014 | ns | 0.130 | 0.029 | ns | 0.372 | 0.015 | ns | 0.295 | 0.010 | ns |
| | | IR64-9a | 0.368 | 0.017 | ns | 0.447 | 0.018 | ns | 0.460 | 0.025 | ns | 0.419 | 0.029 | ns | 0.123 | 0.035 | ns | 0.366 | 0.018 | ns | 0.281 | 0.020 | ns |
| | | IR64-134a | 0.360 | 0.020 | ns | 0.439 | 0.026 | ns | 0.454 | 0.034 | ns | 0.429 | 0.038 | ns | 0.150 | 0.032 | ns | 0.379 | 0.025 | ns | 0.291 | 0.033 | ns |
| | | IR64-134b | 0.324 | 0.003 | ns | 0.421 | 0.004 | ns | 0.436 | 0.003 | ns | 0.399 | 0.003 | ns | 0.113 | 0.020 | ns | 0.359 | 0.006 | ns | 0.295 | 0.002 | ns |
| | | IR64-136a | 0.353 | 0.015 | ns | 0.437 | 0.020 | ns | 0.451 | 0.028 | ns | 0.416 | 0.024 | ns | 0.118 | 0.009 | ns | 0.370 | 0.008 | ns | 0.280 | 0.014 | ns |
| | | IR64-7b | 0.356 | 0.015 | ns | 0.448 | 0.021 | ns | 0.461 | 0.026 | ns | 0.430 | 0.022 | ns | 0.128 | 0.003 | ns | 0.376 | 0.011 | ns | 0.296 | 0.015 | ns |
| | | IR64-134d | 0.354 | 0.010 | ns | 0.440 | 0.011 | ns | 0.456 | 0.014 | ns | 0.422 | 0.010 | ns | 0.135 | 0.013 | ns | 0.376 | 0.006 | ns | 0.290 | 0.007 | ns |
| | NDVI | Ciherang-Sub1 WT | 0.837 | 0.012 | | 0.883 | 0.007 | | 0.923 | 0.005 | | 0.867 | 0.005 | | 0.730 | 0.005 | | 0.812 | 0.003 | | 0.773 | 0.003 | |
| | | CS-3b | 0.836 | 0.015 | ns | 0.882 | 0.004 | ns | 0.921 | 0.003 | ns | 0.869 | 0.002 | ns | 0.731 | 0.003 | ns | 0.812 | 0.000 | ns | 0.782 | 0.004 | ns |
| | | CS-1b | 0.832 | 0.015 | ns | 0.880 | 0.010 | ns | 0.921 | 0.007 | ns | 0.865 | 0.007 | ns | 0.729 | 0.012 | ns | 0.810 | 0.008 | ns | 0.775 | 0.009 | ns |
| | | CS-6d | 0.830 | 0.015 | ns | 0.874 | 0.008 | ns | 0.914 | 0.007 | ns | 0.859 | 0.006 | ns | 0.717 | 0.008 | ns | 0.805 | 0.005 | ns | 0.767 | 0.008 | ns |
| | | IR64 WT | 0.823 | 0.010 | | 0.872 | 0.007 | | 0.912 | 0.006 | | 0.858 | 0.005 | | 0.691 | 0.003 | | 0.785 | 0.002 | | 0.740 | 0.002 | |
| | | IR64-5d | 0.845 | 0.003 | ns | 0.890 | 0.002 | ns | 0.926 | 0.002 | ns | 0.869 | 0.002 | ns | 0.704 | 0.006 | ns | 0.794 | 0.001 | ns | 0.746 | 0.002 | ns |
| | | IR64-7a | 0.837 | 0.008 | ns | 0.887 | 0.006 | ns | 0.924 | 0.003 | ns | 0.872 | 0.007 | ns | 0.703 | 0.011 | ns | 0.797 | 0.008 | ns | 0.754 | 0.005 | ns |
| | | IR64-9a | 0.838 | 0.019 | ns | 0.888 | 0.010 | ns | 0.926 | 0.010 | ns | 0.872 | 0.014 | ns | 0.711 | 0.014 | ns | 0.799 | 0.012 | ns | 0.752 | 0.011 | ns |
| | | IR64-134a | 0.832 | 0.012 | ns | 0.882 | 0.011 | ns | 0.922 | 0.010 | ns | 0.874 | 0.015 | ns | 0.715 | 0.016 | ns | 0.805 | 0.016 | ns | 0.758 | 0.015 | ns |
| | | IR64-134b | 0.817 | 0.004 | ns | 0.873 | 0.003 | ns | 0.912 | 0.002 | ns | 0.852 | 0.001 | ns | 0.681 | 0.006 | ns | 0.789 | 0.003 | ns | 0.749 | 0.002 | ns |
| | | IR64-136a | 0.831 | 0.014 | ns | 0.884 | 0.012 | ns | 0.919 | 0.010 | ns | 0.861 | 0.008 | ns | 0.698 | 0.004 | ns | 0.793 | 0.006 | ns | 0.744 | 0.006 | ns |
| | | IR64-7b | 0.827 | 0.018 | ns | 0.884 | 0.013 | ns | 0.922 | 0.008 | ns | 0.873 | 0.008 | ns | 0.705 | 0.006 | ns | 0.799 | 0.007 | ns | 0.754 | 0.006 | ns |
| | | IR64-134d | 0.836 | 0.006 | ns | 0.889 | 0.006 | ns | 0.925 | 0.004 | ns | 0.876 | 0.002 | ns | 0.710 | 0.005 | ns | 0.800 | 0.002 | ns | 0.755 | 0.001 | ns |

| | Spectra | Genotype | 39DAP | SE | P-value | 47DAP | SE | P-value | 53DAP | SE | P-value | 64DAP | SE | P-value | 77DAP | SE | P-value | 83DAP | SE | P-value | 89DAP | SE | P-value |
|---|---|---|---|---|---|---|---|---|---|---|---|---|---|---|---|---|---|---|---|---|---|---|---|
| 100% N | NDRE | Ciherang-Sub1 WT | 0.451 | 0.006 | | 0.55 | 0.01 | | 0.574 | 0.010 | | 0.538 | 0.011 | | 0.298 | 0.024 | | 0.474 | 0.017 | | 0.389 | 0.014 | |
| | | CS-3b | 0.451 | 0.010 | ns | 0.54 | 0.00 | ns | 0.571 | 0.002 | ns | 0.539 | 0.003 | ns | 0.266 | 0.021 | ns | 0.483 | 0.006 | ns | 0.397 | 0.004 | ns |
| | | CS-1b | 0.453 | 0.012 | ns | 0.56 | 0.00 | ns | 0.582 | 0.002 | ns | 0.551 | 0.013 | ns | 0.299 | 0.003 | ns | 0.516 | 0.022 | ns | 0.440 | 0.028 | * |
| | | CS-6d | 0.422 | 0.002 | ns | 0.52 | 0.003 | * | 0.538 | 0.003 | ns | 0.503 | 0.005 | * | 0.248 | 0.026 | ns | 0.455 | 0.005 | ns | 0.379 | 0.006 | ns |
| | | IR64 WT | 0.451 | 0.018 | | 0.56 | 0.01 | | 0.584 | 0.007 | | 0.580 | 0.005 | | 0.265 | 0.019 | | 0.486 | 0.010 | | 0.416 | 0.007 | |
| | | IR64-5d | 0.474 | 0.012 | ns | 0.58 | 0.01 | ns | 0.598 | 0.008 | ns | 0.579 | 0.006 | ns | 0.229 | 0.013 | ns | 0.464 | 0.010 | ns | 0.397 | 0.008 | ns |
| | | IR64-7a | 0.463 | 0.018 | ns | 0.56 | 0.01 | ns | 0.590 | 0.017 | ns | 0.585 | 0.021 | ns | 0.246 | 0.036 | ns | 0.471 | 0.016 | ns | 0.415 | 0.019 | ns |
| | | IR64-9a | 0.466 | 0.029 | ns | 0.57 | 0.02 | ns | 0.594 | 0.015 | ns | 0.579 | 0.010 | ns | 0.250 | 0.011 | ns | 0.467 | 0.003 | ns | 0.411 | 0.008 | ns |
| | | IR64-134a | 0.449 | 0.014 | ns | 0.55 | 0.01 | ns | 0.577 | 0.012 | ns | 0.555 | 0.016 | ns | 0.231 | 0.041 | ns | 0.453 | 0.023 | ns | 0.393 | 0.025 | ns |
| | | IR64-134b | 0.426 | 0.021 | ns | 0.55 | 0.01 | ns | 0.571 | 0.011 | ns | 0.550 | 0.014 | ns | 0.220 | 0.030 | ns | 0.474 | 0.029 | ns | 0.407 | 0.025 | ns |
| | | IR64-136a | 0.456 | 0.016 | ns | 0.56 | 0.01 | ns | 0.580 | 0.005 | ns | 0.563 | 0.004 | ns | 0.224 | 0.006 | ns | 0.469 | 0.004 | ns | 0.405 | 0.004 | ns |
| | | IR64-7b | 0.465 | 0.007 | ns | 0.57 | 0.00 | ns | 0.593 | 0.004 | ns | 0.565 | 0.019 | ns | 0.208 | 0.024 | ns | 0.444 | 0.018 | ns | 0.384 | 0.017 | ns |
| | | IR64-134d | 0.451 | 0.015 | ns | 0.54 | 0.01 | ns | 0.563 | 0.008 | ns | 0.540 | 0.012 | ns | 0.210 | 0.018 | ns | 0.465 | 0.010 | ns | 0.393 | 0.011 | ns |
| | NDVI | Ciherang-Sub1 WT | 0.859 | 0.002 | | 0.91 | 0.00 | | 0.939 | 0.003 | | 0.898 | 0.004 | | 0.771 | 0.010 | | 0.838 | 0.008 | | 0.797 | 0.010 | |
| | | CS-3b | 0.855 | 0.006 | ns | 0.91 | 0.00 | ns | 0.939 | 0.002 | ns | 0.908 | 0.002 | ns | 0.796 | 0.009 | ns | 0.849 | 0.001 | ns | 0.806 | 0.001 | ns |
| | | CS-1b | 0.852 | 0.010 | ns | 0.91 | 0.00 | ns | 0.940 | 0.001 | ns | 0.901 | 0.008 | ns | 0.795 | 0.024 | ns | 0.851 | 0.012 | ns | 0.813 | 0.014 | ns |
| | | CS-6d | 0.836 | 0.003 | ns | 0.89 | 0.00 | ns | 0.926 | 0.002 | *** | 0.888 | 0.001 | ns | 0.759 | 0.012 | ns | 0.832 | 0.002 | ns | 0.789 | 0.004 | ns |
| | | IR64 WT | 0.887 | 0.011 | | 0.93 | 0.00 | | 0.952 | 0.001 | | 0.935 | 0.002 | | 0.820 | 0.017 | | 0.869 | 0.006 | | 0.821 | 0.005 | |
| | | IR64-5d | 0.896 | 0.010 | ns | 0.93 | 0.00 | ns | 0.953 | 0.002 | ns | 0.930 | 0.002 | ns | 0.789 | 0.011 | ns | 0.855 | 0.006 | ns | 0.811 | 0.006 | ns |
| | | IR64-7a | 0.894 | 0.008 | ns | 0.93 | 0.00 | ns | 0.949 | 0.002 | ns | 0.932 | 0.005 | ns | 0.801 | 0.020 | ns | 0.860 | 0.008 | ns | 0.820 | 0.010 | ns |
| | | IR64-9a | 0.895 | 0.020 | ns | 0.93 | 0.00 | ns | 0.954 | 0.002 | ns | 0.934 | 0.003 | ns | 0.796 | 0.010 | ns | 0.862 | 0.002 | ns | 0.823 | 0.005 | ns |
| | | IR64-134a | 0.880 | 0.012 | ns | 0.92 | 0.00 | ns | 0.947 | 0.002 | ns | 0.923 | 0.006 | ns | 0.784 | 0.024 | ns | 0.851 | 0.013 | ns | 0.810 | 0.016 | ns |
| | | IR64-134b | 0.869 | 0.010 | ns | 0.92 | 0.00 | ns | 0.945 | 0.004 | ns | 0.923 | 0.005 | ns | 0.789 | 0.030 | ns | 0.864 | 0.015 | ns | 0.822 | 0.013 | ns |
| | | IR64-136a | 0.885 | 0.016 | ns | 0.93 | 0.00 | ns | 0.948 | 0.002 | ns | 0.925 | 0.002 | ns | 0.785 | 0.008 | ns | 0.857 | 0.005 | ns | 0.814 | 0.003 | ns |
| | | IR64-7b | 0.887 | 0.003 | ns | 0.93 | 0.00 | ns | 0.952 | 0.001 | ns | 0.926 | 0.007 | ns | 0.769 | 0.025 | * | 0.847 | 0.010 | ns | 0.804 | 0.010 | ns |
| | | IR64-134d | 0.886 | 0.017 | ns | 0.92 | 0.00 | ns | 0.944 | 0.0003 | * | 0.921 | 0.003 | * | 0.785 | 0.010 | ns | 0.855 | 0.005 | ns | 0.812 | 0.007 | ns |
