## Supplementary material for "From lab to field: analyses of genome-edited bacterial blight resistant rice": Table S10

Table S10: Quantification of agronomic traits if GE plants under 0%, 50%, and 100% nitrogen treatment in EPF in Colombia, 2021

| NUE | Genotype | Dry Biomass (g) | SE | P-Value | Grain Yield (Kg/m2) | SE | P-Value | Harvest Index | SE | P-Value | Panicle Number | SE | P-Value | Plant height (cm) | SE | P-Value |
|---|---|---|---|---|---|---|---|---|---|---|---|---|---|---|---|---|
| | Ciherang-Sub1 WT | 18.99 | 0.75 | | 0.63 | 0.03 | | 0.57 | 0.00 | | 8.10 | 0.29 | | 77.57 | 0.59 | |
| | CS-3b | 19.21 | 0.67 | ns | 0.69 | 0.03 | ns | 0.59 | 0.01 | ns | 8.37 | 0.26 | ns | 76.90 | 0.66 | ns |
| | CS-1b | 19.98 | 0.51 | ns | 0.75 | 0.02 | * | 0.60 | 0.00 | ns | 7.80 | 0.19 | ns | 79.80 | 0.45 | ns |
| | CS-6d | 19.22 | 0.67 | ns | 0.64 | 0.02 | ns | 0.57 | 0.01 | ns | 7.73 | 0.23 | ns | 76.10 | 0.63 | ns |
| | IR64 WT | 18.71 | 0.53 | | 0.61 | 0.02 | | 0.57 | 0.01 | | 9.73 | 0.26 | | 78.53 | 0.69 | |
| | IR64-5d | 15.55 | 0.46 | ** | 0.56 | 0.02 | ns | 0.59 | 0.00 | ns | 9.33 | 0.26 | ns | 74.60 | 0.46 | * |
| 0% | IR64-7a | 17.11 | 0.63 | ns | 0.58 | 0.03 | ns | 0.57 | 0.02 | ns | 9.63 | 0.26 | ns | 77.13 | 0.52 | ns |
| | IR64-9a | 16.68 | 0.43 | ns | 0.60 | 0.02 | ns | 0.59 | 0.00 | ns | 9.50 | 0.19 | ns | 76.63 | 0.53 | ns |
| | IR64-134a | 19.19 | 0.59 | ns | 0.54 | 0.02 | ns | 0.52 | 0.02 | ** | 9.00 | 0.24 | ns | 77.17 | 0.46 | ns |
| | IR64-134b | 18.44 | 0.87 | ns | 0.59 | 0.03 | ns | 0.56 | 0.01 | ns | 10.60 | 0.39 | ns | 77.23 | 0.40 | ns |
| | IR64-136a | 20.00 | 0.80 | ns | 0.58 | 0.03 | ns | 0.53 | 0.01 | * | 10.73 | 0.43 | ns | 74.83 | 0.63 | * |
| | IR64-7b | 15.63 | 0.47 | ** | 0.59 | 0.02 | ns | 0.60 | 0.00 | ns | 9.23 | 0.29 | ns | 73.43 | 0.50 | * |
| | IR64-134d | 20.10 | 0.49 | ns | 0.57 | 0.02 | ns | 0.53 | 0.01 | * | 9.30 | 0.25 | ns | 78.07 | 0.52 | ns |
| | Ciherang-Sub1 WT | 27.45 | 0.93 | | 0.92 | 0.03 | | 0.57 | 0.01 | | 11.93 | 0.39 | | 79.33 | 0.61 | |
| | CS-3b | 28.25 | 0.77 | ns | 1.01 | 0.03 | ns | 0.59 | 0.00 | ns | 11.90 | 0.38 | ns | 81.17 | 0.70 | ns |
| | CS-1b | 25.97 | 0.81 | ns | 0.90 | 0.02 | ns | 0.58 | 0.00 | ns | 9.90 | 0.32 | ** | 84.40 | 0.47 | * |
| | CS-6d | 23.66 | 0.56 | * | 0.84 | 0.03 | ns | 0.58 | 0.01 | ns | 9.97 | 0.29 | * | 78.43 | 0.65 | ns |
| | IR64 WT | 23.30 | 1.06 | | 0.81 | 0.03 | | 0.58 | 0.00 | | 11.73 | 0.47 | | 82.00 | 0.72 | |
| | IR64-5d | 25.95 | 0.88 | ns | 0.90 | 0.03 | ns | 0.58 | 0.01 | ns | 13.17 | 0.36 | ns | 83.33 | 0.51 | ns |
| 50% | IR64-7a | 25.16 | 0.84 | ns | 0.97 | 0.03 | * | 0.60 | 0.01 | ns | 12.83 | 0.43 | ns | 87.80 | 0.59 | * |
| | IR64-9a | 26.71 | 1.00 | ns | 0.90 | 0.05 | ns | 0.57 | 0.02 | ns | 13.33 | 0.72 | ns | 86.70 | 0.60 | * |
| | IR64-134a | 30.28 | 1.22 | ** | 0.94 | 0.06 | ns | 0.54 | 0.02 | * | 13.23 | 0.43 | ns | 88.17 | 0.53 | * |
| | IR64-134b | 27.36 | 0.71 | * | 0.84 | 0.04 | ns | 0.54 | 0.02 | ns | 14.80 | 0.39 | ** | 81.97 | 0.43 | ns |
| | IR64-136a | 28.78 | 1.36 | ** | 0.84 | 0.04 | ns | 0.54 | 0.01 | * | 14.23 | 0.50 | ** | 80.87 | 0.68 | ns |
| | IR64-7b | 25.09 | 0.96 | ns | 1.02 | 0.03 | ** | 0.62 | 0.01 | ns | 13.23 | 0.51 | ns | 82.87 | 0.57 | ns |
| | IR64-134d | 27.84 | 0.82 | ** | 0.83 | 0.03 | ns | 0.54 | 0.01 | * | 12.30 | 0.34 | ns | 88.23 | 0.59 | * |
| | Ciherang-Sub1 WT | 32.92 | 0.86 | | 1.12 | 0.03 | | 0.58 | 0.00 | | 12.97 | 0.37 | | 83.93 | 0.64 | |
| | CS-3b | 33.59 | 1.45 | ns | 1.05 | 0.05 | ns | 0.56 | 0.00 | ns | 13.83 | 0.56 | ns | 84.37 | 0.72 | ns |
| | CS-1b | 35.47 | 1.37 | ns | 1.11 | 0.04 | ns | 0.56 | 0.00 | ns | 12.80 | 0.47 | ns | 88.50 | 0.50 | * |
| | CS-6d | 32.04 | 1.48 | ns | 0.99 | 0.04 | ns | 0.55 | 0.01 | ns | 12.70 | 0.52 | ns | 84.07 | 0.70 | ns |
| | IR64 WT | 38.54 | 1.39 | | 1.01 | 0.04 | | 0.51 | 0.01 | | 16.27 | 0.60 | | 93.00 | 0.82 | |
| | IR64-5d | 40.60 | 0.79 | ns | 1.18 | 0.05 | ns | 0.53 | 0.01 | ns | 17.33 | 0.55 | ns | 94.30 | 0.58 | ns |
| 100% | IR64-7a | 37.94 | 1.59 | ns | 1.20 | 0.06 | * | 0.56 | 0.01 | ** | 17.00 | 0.57 | ns | 96.87 | 0.65 | * |
| | IR64-9a | 36.71 | 1.45 | ns | 1.09 | 0.04 | ns | 0.54 | 0.01 | * | 15.67 | 0.61 | ns | 93.63 | 0.64 | ns |
| | IR64-134a | 36.09 | 1.43 | ns | 1.07 | 0.04 | ns | 0.54 | 0.01 | * | 14.47 | 0.52 | ns | 94.20 | 0.56 | ns |
| | IR64-134b | 42.05 | 1.37 | ns | 1.13 | 0.05 | ns | 0.51 | 0.02 | ns | 19.47 | 0.63 | ** | 89.17 | 0.46 | * |
| | IR64-136a | 36.52 | 1.27 | ns | 1.14 | 0.04 | ns | 0.56 | 0.00 | ** | 18.77 | 0.77 | ** | 86.63 | 0.73 | * |
| | IR64-7b | 35.32 | 1.35 | ns | 1.19 | 0.04 | * | 0.57 | 0.00 | ** | 17.13 | 0.74 | ns | 91.13 | 0.63 | ns |
| | IR64-134d | 34.57 | 0.99 | ns | 1.07 | 0.03 | ns | 0.55 | 0.00 | ** | 15.23 | 0.41 | ns | 89.77 | 0.60 | * |

| NUE | Genotype | Panicle Length (cm) | SE | P-Value | Single Plant grain Yield (g) | SE | P-Value | 1000 Grain Weight (g) | SE | P-Value | 100% Flowering (DAP) | SE | P-Value | Bulk Grain weight (g) | SE | P-Value | Yield (Ton/Ha) | SE | P-Value |
|---|---|---|---|---|---|---|---|---|---|---|---|---|---|---|---|---|---|---|---|
| | Ciherang-Sub1 WT | 21.33 | 0.27 | | 25.36 | 1.16 | | 26.35 | 1.05 | | 77.33 | 1.20 | | 702.33 | 81.93 | | 2.09 | 0.24 | |
| | CS-3b | 22.37 | 0.18 | ns | 27.49 | 1.04 | ns | 28.01 | 1.21 | ns | 79.00 | 0.00 | ns | 794.00 | 74.10 | ns | 2.36 | 0.22 | ns |
| | CS-1b | 23.06 | 0.24 | ** | 29.80 | 0.75 | ** | 28.80 | 0.22 | ** | 81.67 | 2.67 | ns | 825.33 | 41.20 | ns | 2.45 | 0.12 | ns |
| | CS-6d | 21.74 | 0.21 | ns | 25.48 | 0.97 | ns | 27.43 | 0.19 | ns | 78.67 | 0.33 | ns | 696.33 | 41.99 | ns | 2.07 | 0.12 | ns |
| | IR64 WT | 23.32 | 0.25 | | 24.57 | 0.74 | | 27.82 | 0.18 | | 73.67 | 0.88 | | 649.67 | 62.88 | | 2.15 | 0.21 | |
| | IR64-5d | 22.34 | 0.25 | ns | 22.37 | 0.87 | ns | 26.93 | 0.13 | ns | 73.67 | 0.88 | ns | 693.67 | 85.18 | ns | 2.47 | 0.30 | ns |
| 0% | IR64-7a | 22.66 | 0.29 | ns | 23.19 | 1.11 | ns | 25.91 | 0.19 | * | 73.67 | 0.33 | ns | 722.00 | 144.66 | ns | 1.78 | 0.36 | ns |
| | IR64-9a | 22.39 | 0.26 | ns | 24.00 | 0.66 | ns | 27.69 | 0.20 | ns | 73.00 | 0.58 | ns | 601.33 | 73.96 | ns | 2.08 | 0.26 | ns |
| | IR64-134a | 22.08 | 0.35 | * | 21.69 | 0.95 | ns | 25.63 | 0.15 | * | 72.00 | 1.15 | ns | 645.00 | 8.54 | ns | 2.19 | 0.03 | ns |
| | IR64-134b | 21.88 | 0.27 | ** | 23.51 | 1.07 | ns | 25.20 | 0.05 | ** | 71.67 | 0.88 | ns | 640.67 | 37.97 | ns | 2.16 | 0.13 | ns |
| | IR64-136a | 21.03 | 0.31 | ** | 23.02 | 1.29 | ns | 24.57 | 0.38 | ** | 73.00 | 1.53 | ns | 557.00 | 40.26 | ns | 1.94 | 0.14 | ns |
| | IR64-7b | 21.01 | 0.41 | ** | 23.42 | 0.67 | ns | 26.86 | 0.25 | ns | 73.67 | 0.33 | ns | 653.33 | 51.49 | ns | 2.26 | 0.18 | ns |
| | IR64-134d | 22.36 | 0.26 | ns | 22.71 | 0.64 | ns | 25.91 | 0.24 | * | 73.00 | 0.58 | ns | 607.33 | 21.06 | ns | 2.07 | 0.07 | ns |
| | Ciherang-Sub1 WT | 21.42 | 0.25 | | 36.90 | 1.38 | | 27.57 | 0.06 | | 77.67 | 1.33 | | 1041.67 | 40.99 | | 3.10 | 0.12 | |
| | CS-3b | 22.19 | 0.23 | ns | 40.30 | 1.30 | ns | 27.73 | 0.26 | ns | 77.67 | 1.33 | ns | 1093.67 | 66.24 | ns | 3.24 | 0.20 | ns |
| | CS-1b | 22.97 | 0.23 | ** | 36.09 | 0.93 | ns | 29.29 | 0.18 | ** | 79.00 | 0.00 | ns | 1094.33 | 83.98 | ns | 3.22 | 0.25 | ns |
| | CS-6d | 22.02 | 0.31 | ns | 33.68 | 1.31 | ns | 27.45 | 0.18 | ns | 77.67 | 1.33 | ns | 1013.33 | 43.85 | ns | 3.02 | 0.13 | ns |
| | IR64 WT | 23.55 | 0.26 | | 32.42 | 1.33 | | 28.58 | 0.23 | | 70.67 | 0.67 | | 902.00 | 56.66 | | 2.92 | 0.18 | |
| | IR64-5d | 23.46 | 0.14 | ns | 35.91 | 1.26 | ns | 27.52 | 0.16 | ** | 70.00 | 0.00 | ns | 1064.33 | 21.94 | ns | 3.51 | 0.07 | ns |
| 50% | IR64-7a | 23.97 | 0.21 | ns | 38.78 | 1.37 | * | 26.61 | 0.05 | ** | 72.00 | 1.15 | ns | 1133.33 | 20.79 | ns | 3.79 | 0.07 | * |
| | IR64-9a | 23.40 | 0.15 | ns | 36.07 | 1.80 | ns | 27.54 | 0.19 | ** | 70.00 | 0.00 | ns | 1102.33 | 95.73 | ns | 3.54 | 0.31 | ns |
| | IR64-134a | 23.87 | 0.17 | ns | 38.86 | 1.88 | * | 26.12 | 0.05 | ** | 70.00 | 0.00 | ns | 1016.67 | 90.71 | ns | 3.30 | 0.29 | ns |
| | IR64-134b | 22.84 | 0.20 | ns | 33.55 | 1.41 | ns | 25.37 | 0.08 | ** | 72.00 | 1.15 | ns | 937.67 | 56.26 | ns | 2.93 | 0.18 | ns |
| | IR64-136a | 22.15 | 0.26 | ** | 33.67 | 1.53 | ns | 24.91 | 0.28 | ** | 70.67 | 0.67 | ns | 878.00 | 48.56 | ns | 2.90 | 0.16 | ns |
| | IR64-7b | 23.20 | 0.22 | ns | 40.86 | 1.37 | ** | 27.12 | 0.05 | ** | 70.67 | 0.67 | ns | 1080.00 | 95.00 | ns | 3.55 | 0.31 | ns |
| | IR64-134d | 23.84 | 0.18 | ns | 33.19 | 1.26 | ns | 25.75 | 0.24 | ** | 72.67 | 1.33 | ns | 1073.33 | 16.27 | ns | 3.53 | 0.05 | ns |
| | Ciherang-Sub1 WT | 21.85 | 0.25 | | 44.72 | 1.30 | | 27.34 | 0.07 | | 79.00 | 0.00 | | 1157.67 | 51.35 | | 3.43 | 0.15 | |
| | CS-3b | 22.15 | 0.23 | ns | 41.94 | 1.81 | ns | 28.16 | 0.13 | ns | 79.00 | 0.00 | ns | 1295.33 | 37.13 | ns | 3.87 | 0.11 | ns |
| | CS-1b | 23.35 | 0.22 | ** | 44.20 | 1.62 | ns | 29.22 | 0.13 | ** | 81.00 | 1.00 | ns | 1223.33 | 42.12 | ns | 3.64 | 0.13 | ns |
| | CS-6d | 21.45 | 0.23 | ns | 39.74 | 1.72 | ns | 27.63 | 0.32 | ns | 79.00 | 0.00 | ns | 1083.00 | 20.65 | ns | 3.23 | 0.06 | ns |
| | IR64 WT | 23.84 | 0.32 | | 40.49 | 1.55 | | 28.01 | 0.14 | | 76.33 | 1.33 | | 1095.00 | 62.04 | | 3.67 | 0.21 | |
| | IR64-5d | 23.28 | 0.25 | ns | 47.14 | 1.96 | ns | 27.04 | 0.26 | ns | 74.67 | 0.33 | ns | 1184.67 | 10.27 | ns | 4.01 | 0.03 | ns |
| 100% | IR64-7a | 23.94 | 0.28 | ns | 47.90 | 2.29 | * | 25.94 | 0.22 | ** | 74.67 | 0.33 | ns | 1232.67 | 22.15 | ns | 4.25 | 0.08 | ns |
| | IR64-9a | 23.65 | 0.18 | ns | 43.61 | 1.52 | ns | 27.46 | 0.17 | ns | 74.33 | 0.33 | ns | 1189.00 | 45.37 | ns | 3.99 | 0.15 | ns |
| | IR64-134a | 24.20 | 0.20 | ns | 42.78 | 1.61 | ns | 26.49 | 0.71 | * | 73.33 | 0.67 | ns | 1214.67 | 58.12 | ns | 4.02 | 0.19 | ns |
| | IR64-134b | 23.57 | 0.17 | ns | 45.31 | 1.90 | ns | 25.91 | 0.05 | ** | 75.33 | 2.03 | ns | 1092.33 | 87.89 | ns | 3.64 | 0.29 | ns |
| | IR64-136a | 23.20 | 0.16 | ns | 45.73 | 1.65 | ns | 25.31 | 0.98 | ** | 76.33 | 1.33 | ns | 892.67 | 78.85 | ns | 2.99 | 0.26 | ns |
| | IR64-7b | 23.23 | 0.24 | ns | 47.64 | 1.65 | * | 26.81 | 0.20 | ns | 74.33 | 2.33 | ns | 1354.00 | 109.30 | * | 4.46 | 0.36 | * |
| | IR64-134d | 24.18 | 0.22 | ns | 42.61 | 1.38 | ns | 25.61 | 0.34 | ** | 75.00 | 0.00 | ns | 1102.00 | 78.02 | ns | 3.69 | 0.26 | ns |

SE: standard error; *: significant at P<0.05; **: significant at P<0.01;  ns = not significant (P>0.05)

N.A: Not applicable; n.d.: not determined
