## Supplementary material for "From lab to field: analyses of genome-edited bacterial blight resistant rice": Table S11

Table S10: Accession numbers for whole genome sequences of WT and GE'd lines used in this study

| Accession number | SRA | BioSample name |
|---|---|---|
| SAMN47442430 | SRR32777075 | IR64-C3 rep 3 |
| SAMN47442429 | SRR32777076 | IR64-C3 rep 2 |
| SAMN47418954 | SRR32777077 | IR64-C3 rep 1 |
| SAMN47442428 | SRR32838284 | IR64-136a rep 4 |
| SAMN47442427 | SRR32838285 | IR64-136a rep 3 |
| SAMN47442419 | SRR32838286 | IR64 WT rep 6 |
| SAMN47442418 | SRR32838287 | IR64 WT rep 5 |
| SAMN47442437 | SRR32838288 | CS-4 rep 4 |
| SAMN47442436 | SRR32838289 | CS-4 rep 3 |
| SAMN47442434 | SRR32838290 | CS-3b rep 3 |
| SAMN47442433 | SRR32838291 | CS-3b rep 2 |
| SAMN47442438 | SRR32838292 | KOM-14-19 rep 2 |
| SAMN47442443 | SRR32838293 | KOM WT rep 6 |
| SAMN47442442 | SRR32838294 | KOM WT rep 5 |
| SAMN47442441 | SRR32838295 | KOM WT rep 4 |
| SAMN47442422 | SRR32838296 | IR65-5d rep 4 |
| SAMN47442421 | SRR32838297 | IR65-5d rep 3 |
| SAMN47442425 | SRR32838298 | IR64-9a rep 4 |
| SAMN47442424 | SRR32838299 | IR64-9a rep 3 |
| SAMN47442414 | SRR32838300 | IR64-7a rep 4 |
| SAMN47442413 | SRR32838301 | IR64-7a rep 3 |
| SAMN47442432 | SRR32838302 | CS WT rep 3 |
| SAMN47442431 | SRR32838303 | CS WT rep 2 |
| SAMN47442412 | SRR32857392 | IR64-7a rep 2 |
| SAMN47418949 | SRR32857393 | IR64-7a rep 1 |
| SAMN47442426 | SRR32857394 | IR64-136a rep 2 |
| SAMN47418953 | SRR32857395 | IR64-136 rep 1 |
| SAMN47442415 | SRR32857396 | IR64 WT rep 2 |
| SAMN47418950 | SRR32857397 | IR64 WT rep 1 |
| SAMN47442435 | SRR32857398 | CS-4 rep 2 |
| SAMN47418957 | SRR32857399 | CS-4 rep 1 |
| SAMN47418958 | SRR32857400 | KOM-14-1 rep 1 |
| SAMN47442440 | SRR32857401 | KOM WT rep 3 |
| SAMN47442439 | SRR32857402 | KOM WT rep 2 |
| SAMN47418959 | SRR32857403 | KOM WT rep 1 |
| SAMN47442423 | SRR32857404 | IR64-9a rep 2 |
| SAMN47418952 | SRR32857405 | IR64-9a rep 1 |
| SAMN47418956 | SRR32857406 | CS-3b rep 1 |
| SAMN47418955 | SRR32857407 | CS WT rep 1 |
| SAMN47442416 | Need to complete | IR64 WT rep 3 |
| SAMN47442417 | Need to complete | IR64 WT rep 4 |
| SAMN47442420 | Need to complete | IR65-5d rep 2 |
| SAMN47442444 | Need to complete | KOM-14-65 rep 2 |
