## Supplementary material for "From lab to field: analyses of genome-edited bacterial blight resistant rice": Table S12

Table S11: Short IDs and their corresponding line full ID for each GE'd line used in this study

| Cultivar | Short ID | Line full ID |
|---|---|---|
| IR64 | IR64-5a | IR64-IRS1132-5-18-1 |
| | IR64-5b | IR64-IRS1132-5-33-1 |
| | IR64-5c | IR64-IRS1132-5-47-1 |
| | IR64-5d | IR64-IRS1132-5-55-1 |
| | IR64-7a | IR64-IRS1132-7-18-1-2-1 |
| | IR64-7b | IR64-IRS1132-7-45-1 |
| | IR64-9a | IR64-IRS1132-9-32-1 |
| | IR64-9b | IR64-IRS1132-9-38-1 |
| | IR64-106 | IR64-IRS1132-106-4-1 |
| | IR64-134a | IR64-IRS1132-134-4-1 |
| | IR64-134b | IR64-IRS1132-134-37-1 |
| | IR64-134c | IR64-IRS1132-134-53-1-2 |
| | IR64-134d | IR64-IRS1132-134-53-1-8-B |
| | IR64-136a | IR64-IRS1132-136-3-1 |
| | IR64-C3 | IR64-IRS1132-134-003–3-3.3–3B3–C3 |
| | IR64-134e | IR64-IRS1132-134-003 |
| | IR64-136b | IR64-IRS1132-136-38-01 |
| Ciherang-Sub1 | CS-1a | Ciherang-Sub1-IRS1132-1-9 |
| | CS-1b | Ciherang-Sub1-IRS1132-1-11 |
| | CS-1c | Ciherang-Sub1-IRS1132-1-14 |
| | CS-1d | Ciherang-Sub1-IRS1132-1-16 |
| | CS-1e | Ciherang-Sub1-IRS1132-1-18 |
| | CS-1f | Ciherang-Sub1-IRS1132-1-23 |
| | CS-1g | Ciherang-Sub1-IRS1132-1-29 |
| | CS-1h | Ciherang-Sub1-IRS1132-1-30 |
| | CS-1i | Ciherang-Sub1-IRS1132-1-17 |
| | CS-3a | Ciherang-Sub1-IRS1132-3-12 |
| | CS-3b | Ciherang-Sub1-IRS1132-3-23-1-B-1 |
| | CS-4a | Ciherang-Sub1-IRS1132-4-24-1-B |
| | CS-4b | Ciherang-Sub1-IRS1132-4-24-24-B |
| | CS-6a | Ciherang-Sub1-IRS1132-6-6 |
| | CS-6b | Ciherang-Sub1-IRS1132-6-8 |
| | CS-6c | Ciherang-Sub1-IRS1132-6-12 |
| | CS-6d | Ciherang-Sub1-IRS1132-6-19 |
| | CS-6e | Ciherang-Sub1-IRS1132-6-27 |
| | CS1-6f | Ciherang-Sub1-IRS1132-6-28 |
| | CS1-6g | Ciherang-Sub1-IRS1132-6-30 |
| | CS-3c | Ciherang-Sub1-IRS1132-3-23-1-B-7 |
| | CS-3d | Ciherang-Sub1-IRS1132-3-17-1 |
| Komboka | KOM-14-65 | Komboka-pMUGW5-14-65-10-1 |
| | KOM-14-19 | Komboka-pMUGW5-14-19-1 |
| | KOM-18-68 | Komboka-pMUGW5-18-68 |
| | KOM-9-33 | Komboka-pMUGW5-9-33 |
